# C7-substituted Quinolines as Potent Inhibitors of AdeG Efflux Pumps in *Acinetobacter baumannii*

**DOI:** 10.1101/2024.09.07.611778

**Authors:** Yiling Zhu, Charlotte K Hind, Taha Al-Adhami, Matthew E. Wand, Melanie Clifford, J. Mark Sutton, Khondaker Miraz Rahman

**Author notes:** Corresponding authors: MS, KMR.

## Abstract

Efflux, mediated by a series of multidrug efflux pumps, is a major contributor to antibiotic resistance in Gram-negative bacteria. Efflux pump inhibitors (EPIs), which can block efflux, have the potential to be used as adjuvant therapies to re-sensitize bacteria to existing antibiotics. In this study, 36 quinoline-based compounds were synthesized as potential EPIs targeting Resistance Nodulation Division (RND) family pumps in the multidrug-resistant pathogen *Acinetobacter baumannii* . In *A. baumannii* strains with overexpressed AdeFGH (chloramphenicol-adapted) and AdeABC (AYE, Ab5075-UW), these compounds enhanced Hoechst dye accumulation, indicating general efflux inhibition, and potentiated chloramphenicol which is an AdeG substrate. The research focused on two generations of quinoline compounds, with modifications at the C-7 position of first-generation compounds to improve hydrophobic interactions with the Phe loop in the AdeG efflux pump, to generate second-generation compounds. The modified quinolines showed strong pump inhibition and significant chloramphenicol potentiation, with MIC reductions of 4- to 64-fold. Notably, compounds **1.8** and **3.8** exhibited the highest inhibitory activity, while compounds **1.3** and **3.3** showed up to 64-fold potentiation, highlighting the importance of specific structural features at the C-7 position for efflux pump inhibition. The study also revealed selective inhibition of AdeFGH over AdeABC, with no potentiation observed for gentamicin, showing the specificity of these quinoline-based inhibitors. Importantly, the compounds showed no toxicity in a Galleria mellonella model at a 50 mg/kg dose level, highlighting their suitability as potential antibiotic adjuvants for combating bacterial resistance.

---

Antimicrobial resistance (AMR) has been recognized as one of the most critical global health threats. Although AMR occurs as a natural evolutionary process, other factors contribute to its rapid development, including overuse and incorrect use of antimicrobial drugs, use as food additives for food-producing animals, increasing spread of resistant microorganisms between humans and the environment, and lacking of rapid diagnosis methods.[1, 2] With few novel antibiotics being developed in recent decades, AMR has become one of the most critical global health threats.

*Acinetobacter baumannii* is a major nosocomial pathogen involved in epidemic infections. Carbapenem-resistant *A. baumannii* has been rated by the WHO as one of the highest priority species, for which development of new drugs is critically required.[3] Like other Gram-negative bacteria, *A. baumannii* confers resistance through various mechanisms, including decreased penetration, increased efflux, overproduction of drug targets, drug target modification, enzymatic inactivation or modification of drugs, and antibiotics bypass pathway. [4]

In *A. baumannii,* three major RND-type efflux pumps, AdeABC, AdeFGH and AdeIJK, are each associated with extrusion of various antibiotics. AdeABC is the first RND-type efflux pump that was extensively studied in *A. baumannii.* Significant overexpression of AdeB is seen in many MDR *A. baumannii* isolates that are resistant to tigecycline, a major drug for treatment of *A. baumannii* infection. In an AdeABC-overexpressing strain, tigecycline MIC increased by 16-fold compared to the isogenic parental strains.[5] The expression of *adeABC* is positively controlled by a two-component regulatory system AdeRS. Studies have shown that mutations in *adeR* and *adeS* restored the sensitivity to aminoglycosides.[6, 7] Among the three structural deletions of AdeB in the mutants, a 4- to 32-fold decrease in MIC of various antibiotics was observed compared to the parental strains.[6] AdeC is less critical in contributing to MDR resistance, possibly due to the redundancy of OMPs. For instance, AdeK could substitute as the OMP in cooperation with AdeA and AdeB. [8]

AdeIJK contributes to the intrinsic resistance to β-lactams, fluoroquinolones, tetracyclines, chloramphenicol, rifampicin and fusidic acid. [9] High-level overexpression of AdeIJK was previously reported to be toxic for the bacteria itself, but overexpressed AdeIJK can be detected in MDR strains [10]. The expression of AdeIJK is regulated by a TetR-type regulator, AdeN and expression of an intact copy of AdeN can restore the susceptibility to antibiotics in strains with a deleted *adeN* [11]. Recently, specific EPIs have been described for AdeIJK. The 4,6-diaminoquoniline analogues were reported to potentiate erythromycin, tetracycline, and novobiocin in both a lab antibiotic susceptible *A. baumannii* strain and multidrug-resistant clinical isolates AB5075 and AYE.[12]

Efflux pump inhibitors (EPIs) have long been considered as useful adjuvants capable of restoring the activity of efflux-substrate antibiotics.[13] This may include antibiotics, like glycopeptides and macrolides, which have very poor activities against Gram-negative bacteria, partially due to their effective export by efflux pump.[14]. A number of EPIs have been identified, either from the natural or the synthetic sources. For example, Reserpine is an indole plant alkaloid and was found to reduce the efflux of tetracycline in MDR *Staphylococcus aureus* by binding to the NorA transporter.[15] However it is not used in the clinical treatment of bacterial infections due to the effective inhibitory concentration is harmful to the kidneys.[16] The most well-known synthetic EPI is Phe-Arg-β- naphthylamide (PAβN), which is discovered with inhibitory activity against multiple effluc pumps including MexAB-OprM in *P. aeruginosa*, AcrAB-ToLC in *E. coli* and AdeFGH in *A.buamannii*. [17, 18] Even though effects were seen from PAβN on inhibiting efflux or causing OM permeation, the further development of it was abandoned due to toxicity issues. Unfortunately, so far, no stand- alone EPIs have progressed to the clinic, but it remains an attractive approach.

A quinoline-based structure (**1**) was identified by in-silico screening of a large fragment library using the homology model of the AdeB and AdeG efflux pumps in *A. baumannii* as a putative efflux pump inhibitor. Previously, compounds containing a quinoline scaffold were shown to inhibit the AcrAB-TolC pump in *Enterobacter aerogenes*.[19] This suggests that quinoline analogues can be developed as inhibitors of RND-type efflux pumps. Therefore, novel quinoline-type EPIs against *A. baumannii* were designed and evaluated by targeting the three RND-type efflux pumps, AdeABC, AdeFGH, and AdeIJK, using compound **1** as the core scaffold. Several candidates were confirmed as potential EPIs with specificity for potentiation of chloramphenicol by inhibition of AdeG. The development and evaluation of these compounds are described, and the basis for selectivity for AdeG is discussed.

## Results and Discussion

### Antibiotic Specificity against Different RND-type Efflux Pumps

Initially, eight antibiotics were screened against two *A. baumannii* strains (AYE and Ab5075-UW) and respective mutants in the three major RND-family efflux pump, AdeABC, AdeFGH and AdeIJK and their respective regulators (AdeRS, AdeL, AdeN) (Figure 1; Strain information in Table S1). The antibiotics were chosen based on published efflux pump substrate specificity, such as with ciprofloxacin [20] or known potentiation by EPIs, regardless of pump identification. Only two antibiotics could be accurately ascribed as substrates for a particular efflux pump, with at least a 4- fold change in MIC; gentamicin and ciprofloxacin (Table S2), which are known substrates of AdeABC in AYE [21]. All other antibiotics showed 2-fold or less change in MIC irrespective of mutation/transposons in any particular pump. This suggests that these antibiotics are not substrates for any of the efflux pumps studied, that there is redundancy amongst these pumps or that expression of the pumps in these strains is at basal levels masking any effect of the mutation. Clearer results are observed in strains where the efflux pump is overexpressed, either due to a naturally occurring mutation in the regulator, as is the case for AdeABC in these strain backgrounds, or by targeted deletion / gain-of function mutations of the repressor/activator respectively [22]. Strains AYE and Ab5075-UW have endogenous mutations in AdeS, which leads to overexpression of AdeABC, resulting in elevated resistance to gentamicin.

**Figure 1.**
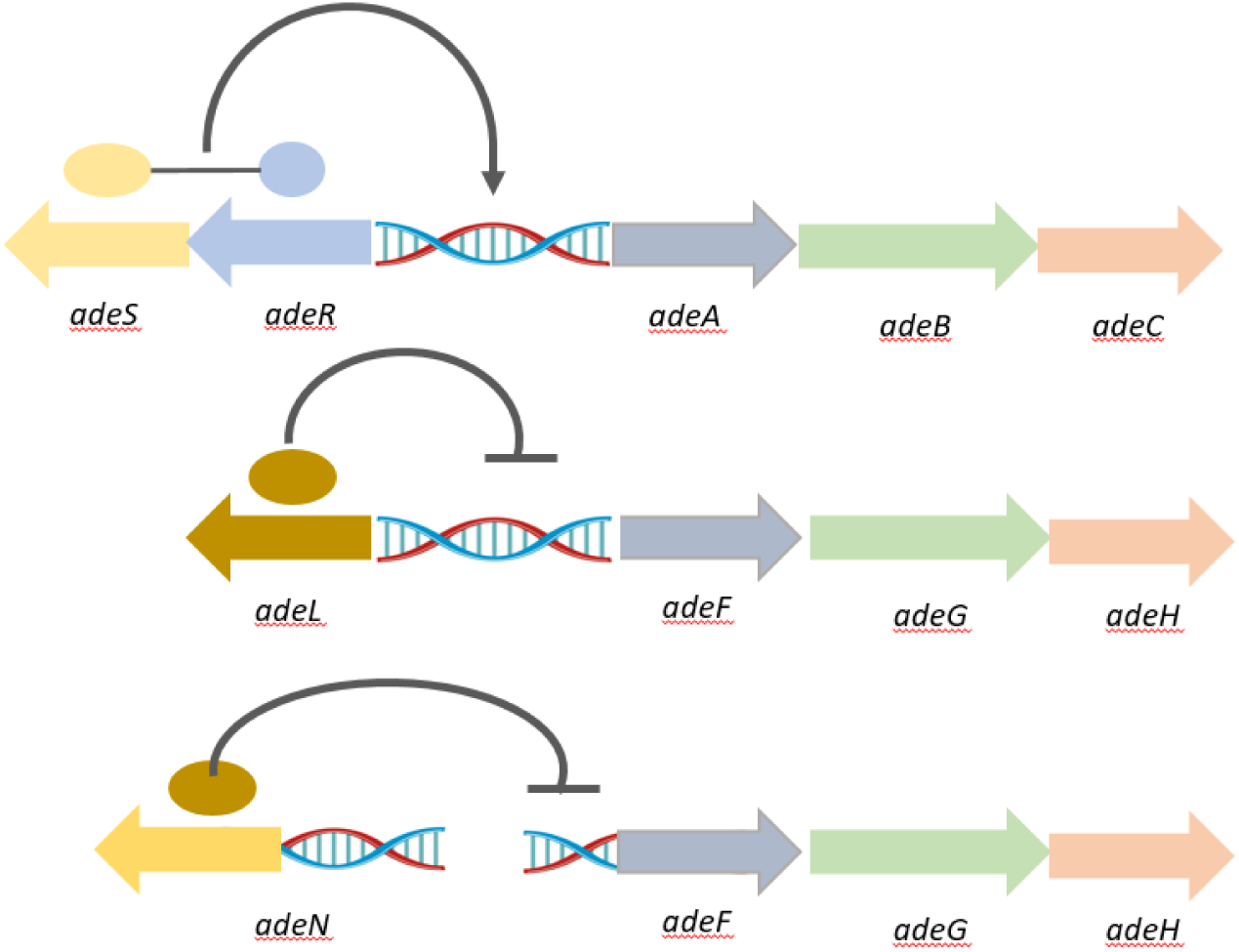
Schematic organization and regulation of three characterized RNA efflux pumps (adeABC- RS, adeFGH-L, and adeIJK-N) genes clusters on A. baumannii chromosome.

Strains with additional overexpressed efflux pumps were generated by adapting WT strains to the antibiotics predicted to be substrates for a particular pump.[23] Looking to overexpress AdeFGH and AdeIJK, strains were adapted on gradient plates (Figure S1) to substrates chloramphenicol and cefotaxime, respectively.[10] Ab5075-UW adapted to chloramphenicol exposure, generating a strain with a 4-fold increased MIC for chloramphenicol. (Table S2) RT-PCR confirmed significant overexpression of AdeG (619-fold, data not shown) and its regulator AdeL (4-fold) and whole genome sequencing confirmed a M270R mutation in the AdeFGH regulator AdeL. No adaptation to cefotaxime was observed for ATCC 17978, which is usually susceptible, perhaps indicating that this is a substrate for more than one pump in the strain tested.

**Chemistry.** Twelve first-generation quinoline-based compounds were synthesized according to Scheme 1, divided into two groups based on the presence (compounds **1** - **6**) or absence (compounds **7** - **12**) of a bromine atom at position 7 of the quinoline ring. Each group featured six different substitutions (R groups) at position C-2, including open chains, aliphatic cyclic rings, and aromatic rings, designed to explore interactions with a target binding pocket.

The synthesis involved an SN2 nucleophilic substitution reaction using two starting materials: 7- bromo-4,8-dimethylquinolin-2-ol and 4,8-dimethylquinolin-2-ol. Depending on the chemical properties of the starting material, three different reaction conditions were employed. Condition 1 (overnight reflux in acetone) successfully synthesized compounds **1**, **2**, and **6** with yields ranging from 36% to 78%. Condition 2 (overnight reflux in DMF) was used when Condition 1 failed, yielding 48% and 19% for compounds **3** and **7**, respectively. Condition 3 (microwave heating in DMF) was applied when previous conditions were unsuccessful, producing the remaining compounds with yields between 14% and 44%.

The first-generation compounds were modelled against both AdeB and AdeG transporter proteins to understand the basis of the observed selectivity. Since the crystal structure of AdeB and AdeG are not available, the molecular models were developed using a homology modelling approach with the Swiss Model webserver using published crystal structures of AcrB (1IWG) and MexB (3W9J) as templates. The poses with the best ChemScore and ΔG value for compounds **1** and **3** against both efflux pump transporters are shown (Table S3). Notably, there are high affinity interactions between both compounds and two Phe (Phe293 and Phe624) residues within the binding pocket of AdeG but only one in AdeB (Phe612).(Figure S2) This could suggest that multiple Phe residues in AdeG are important in determining inhibition and this is consistent with the known importance of a Phe loop in several efflux transporters, including AdeB [24]. Monitoring the location and orientation of the compounds in diverse complexes, showed that both compounds are completely trapped in the distal pocket of the multi binding site of AdeG whereas they are only partially trapped in the case of AdeB. Hence, the compounds are likely to form much more stable pump-inhibitor complexes in AdeG, than in the binding site of AdeB the C-7 site, the Br group of the compound was located in the hydrophobic part of the distal binding pocket of the efflux pump. Based on this, the C-7 site was selected for the introduction of further hydrophobic side chains that might interact with the hydrophobic pockets more efficiently than those in compounds **1** or **3**, with the aim of developing more potent efflux pump inhibitors. We selected a diverse range of hydrophobic C-7 side chains, including a phenyl ring, 5- and 6-membered heterocycles, benzofused heterocycles, a second quinoline ring, a naphthalene ring, and a flexible side chain with a terminal phenyl ring.

A second-generation of compounds was designed based on the best performing first-generation compounds, **1** and **3.** Twenty-four second-generation quinoline-based compounds, designed to increase the interaction with the hydrophobic Phe loop of AdeG, were synthesised using solution phase chemistry (Schemes 2 and 3). Among these, 18 compounds featured two methyl groups at positions C-4 and C-8 of the quinoline ring, while six compounds lacked these methyl groups. The 18 compounds containing C-4 and C-8 methyl groups were derived from the first-generation compounds **1** (**1.1** - **1.9**) and **3** (**3.1** - **3.9**) through a Suzuki coupling reaction (Scheme 2). This reaction involved the use of a boronic acid, an organohalide, and a palladium (0) complex catalyst, specifically Tetrakis (triphenylphosphine) palladium (0) (Pd(PPh3)4) (Scheme 2). The boronic acids used as starting materials were commercially obtained, and the reaction was carried out in a mixture of toluene and methanol under reflux for 6 hours or overnight.

The reaction conditions were consistent across the 18 compounds, where the quinoline starting material was reacted with the boronic acid in a 1:1.2 molar ratio, using 100 mg of the quinoline compound (Scheme 2). The reaction progress was monitored using thin-layer chromatography (TLC) and liquid chromatography-mass spectrometry (LC-MS). Post-reaction, the products were extracted, dried, and purified via flash column chromatography using a dichloromethane/methanol solvent system.

Additionally, six compounds lacking C-4 and C-8 methyl groups (**1.10 - 1.12** & **3.10 - 3.12)** were synthesized using a two-step process: initial reaction in DMF with K_2_CO_3_ under microwave conditions, followed by Suzuki coupling (Scheme 3). All synthesized compounds were characterized using 1H-NMR, 13C-NMR, LC-MS, and HR-MS, with purity confirmed by LC-MS analysis.

### Efflux Inhibitory Activity of First-generation Quinoline-type EPI Compounds

Direct antimicrobial activities of the quinoline derivatives were measured on *A. baumannii* AYE and Ab5075-UW strains, using MIC tests and growth curve assays. Testing concentrations (Table S4) were identified which had no significant impact on bacterial growth (defined as less than 15% reduction in OD_600_ measured after 20h growth) compared to wild type. Two known EPIs, PAβN and CCCP, were also tested in parallel with the quinoline series using the Hoechst assay.[25] Increased accumulation of the fluorescent dye inside the bacterial cells correlates with reduced efflux activity via one of a number of pumps, through inhibition by the EPIs. AYE and AB5075-UW showed essentially identical accumulation of Hoechst over time, ensuring that studies with efflux pump/regulator mutants in either of these backgrounds are directly relatable to the Hoechst data.

Several of the quinoline compounds induced a rapid increase in fluorescence due to Hoechst accumulation and reached a steady state consistent with activity as EPIs (Figure 2). Compound **12,** containing a quinoline side chain, showed a very different profile of Hoechst accumulation, with the final fluorescence being one of the highest values measured by the end of the incubation. This possibly suggests a very different mechanism of action contributing to a reduction in efflux or inherent fluorescence of the compound due to the presence of the quinoline ring. The efflux inhibitory activity of each compound was calculated by normalising the data at 24 minutes to the fluorescence level of the compound-free (control) cells (Table 1). Four of the compounds tested, compounds **1**, **3**, **4** and **9** showed significantly higher fluorescence accumulation than the control and the others. Generally, for the pair of compounds with the same R_2_ substitution, the one with Br on C- 7 accumulated higher fluorescence in the cells than the other one without Br. Among the four compounds with the best EPI activity, compound **9** is the only compound that does not have the C-7 Br substitution, and its corresponding compounds with C-7 Br, compound **3**, also showed significant inhibitory activity. Interestingly, all four compounds had basic R2 side chain with a tertiary N either part of a heteroaliphatic ring (compounds **1** and **4**) or as dimethyl amine terminal group. Compounds containing naphthalene (compounds **2** and **8**) or quinoline terminal ring (compound **6** and **12**) did not show significant efflux inhibitory activity despite having more hydrophobic side chains.

**Figure 2.**
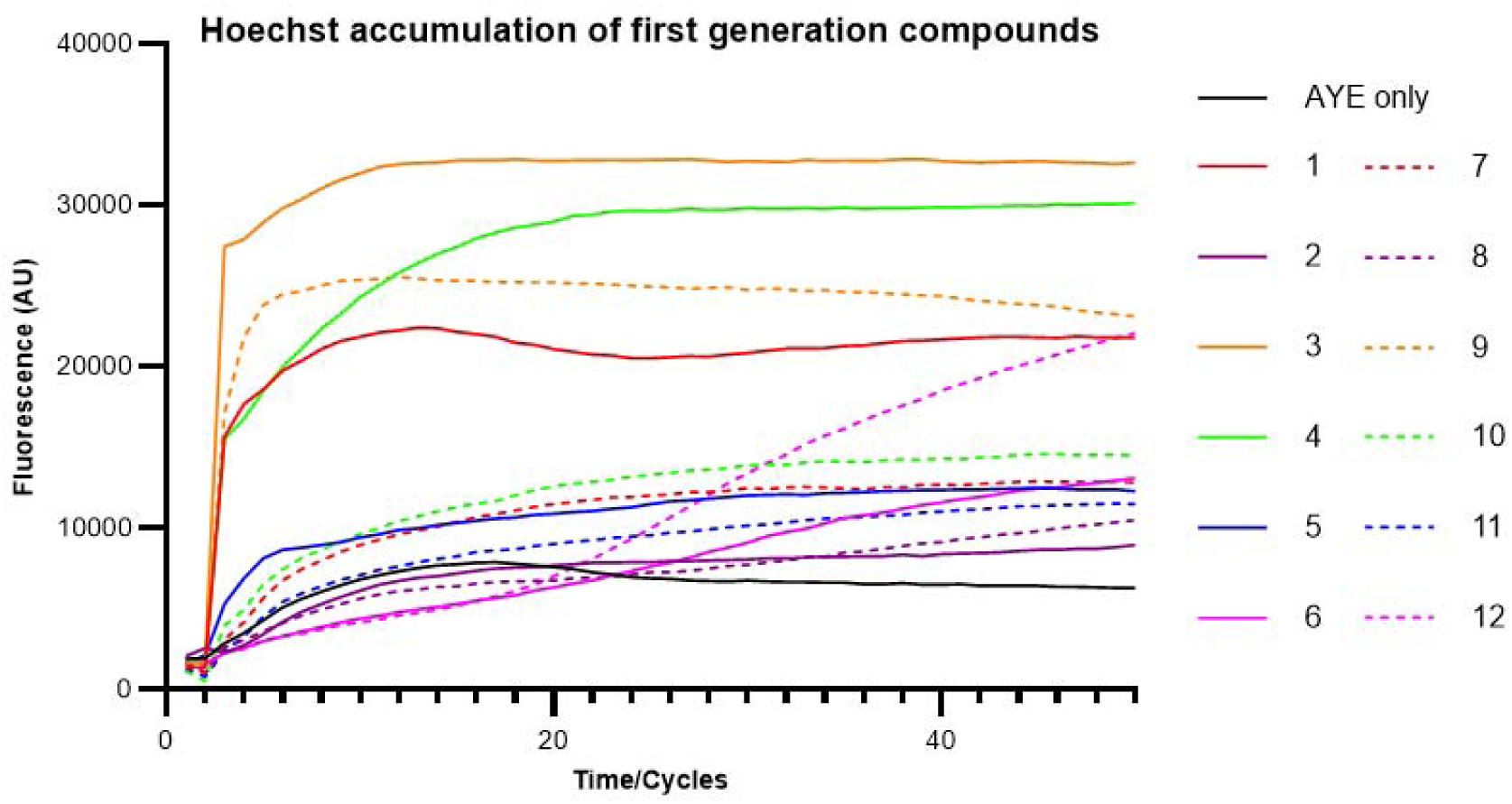
HOECHST DYE accumulation in AYE cells with the addition of synthesised EPI compounds. All results were performed in triplicates and the curve presented is the fluorescence value of three biological repeats after being blanked against cell free PBSM+G with HOECHST DYE; for clarity error bars are not shown on the graph, but the SD is included in the end-point measurements shown in table 1.

**Table 1.** The inhibitory activity of the first-generation compounds against strain Ab5075-UW assessed by changes in Hoechst accumulation, compared to known EPIs.

| Compound ID | Inhibitory activity | P value* | Name | Inhibitory activity | P value |
| --- | --- | --- | --- | --- | --- |
| **1** | 3.12±0.11 | 8.5E-04 | **7** | 1.31±0.32 | 0.26 |
| **2** | 0.91±0.13 | 0.59 | **8** | 0.83±0.14 | 0.27 |
| **3** | 4.57±0.11 | 7.83E-06 | **9** | 3.59±0.28 | 5.0E-04 |
| **4** | 3.53±0.19 | 8.11E-05 | **10** | 1.41±0.22 | 0.07 |
| **5** | 1.36±0.09 | 0.07 | **11** | 1.05±0.10 | 0.75 |
| **6** | 0.64±0.05 | 0.07 | **12** | 0.61±0.04 | 0.03 |
| **PAβN** | 1.17±0.03 | 0.28 | **CCCP** | 2.35±0.13 | 1E-04 |

The inhibitory activity was calculated as the fluorescence level of the cells with EPI compounds divided by that of the EPI-free cells at steady state (24 mins for all compounds except **1.6** and **1.12**). Values over 1, suggest that the compounds promoted accumulation of Hoechst, by efflux inhibition. \*P value: Student t-test was used to calculate the significance, between the cells with and without EPIs. Values in red are significant ( $P \leq 0.05$ ). The results were calculated from 9 measurements from three independent biological replicates.

To investigate whether the EPI candidate compounds targeted AdeABC and/or AdeFGH, the degree of MIC potentiation was tested with chloramphenicol and gentamicin (Table S5). Compounds **1**, **3** and **4** were selected for this study based on their activity in the Hoechst accumulation assay. None of the compounds reduced the MIC of gentamicin in the wildtype strain, and only compound **3** showed a 2-4 fold potentiation of chloramphenicol in the wildtype strain AYE. In the AdeG- overexpressing strain Ab5075-CHL, compounds **1** and **3** potentiated chloramphenicol by 4- and 8- fold, respectively. No gentamicin potentiation was observed for either PaβN or CCCP, consistent with previous data showing that neither are effective inhibitors of AdeABC in *A. baumannii* AYE.[21, 24] Potentiation of further antibiotics, rifampicin, clarithromycin, and colistin, were tested in strain AYE (Table S6). Compounds **1, 3, 4** and CCCP decreased the MIC of colistin by more than 4-fold. There was no evidence of an increase in colistin MIC with either AdeABC or AdeFGH overexpression. It is possible that the observed potentiation of colistin relates to the PMF uncoupling mechanism of CCCP and this may not be linked to efflux directly, with the EPIs also potentially affecting other cellular functions. Compounds **1** and **4** also showed between 4-8-fold potentiation of rifampicin MIC, which was similar to the observed effect of CCCP but much lower potentiation than observed with PaβN (up to 256-fold potentiation; rifampicin MIC 0.125-0.25 µg/mL). Clarithromycin was not potentiated by the three quinoline compounds or CCCP, but the MIC was reduced up to 128-fold by PaβN. This data suggests the presence of at least one additional RND-family efflux pump, which is inhibited by PaβN and capable of effluxing rifampicin and clarithromycin, but other efflux pumps/ PMF-dependent processes may also affect susceptibility to rifampicin.

### Efflux Inhibitory Activity of Second-generation Quinoline-type EPI Compounds

The direct antimicrobial activities of the second-generation compounds were measured, and their optimal testing concentration was determined (Table S4). To allow a systematic evaluation of the efflux pump inhibitory activity between the second-generation compounds and their parent molecules, a single fixed concentration was selected for comparison purposes in the Hoechst assay. A concentration of 25µg/mL was used as this did not affect growth across the range of compounds tested. Compounds **1.5** and **3.5** were not included in the assay as they produced negative fluorescence levels after normalization suggesting that they had strong intrinsic fluorescence (results not shown). Most of the 24 second-generation compounds showed strong efflux inhibitory activity in the Hoechst assay (Figure 3 and Figure S2). At the concentration of 25 µg/mL, compounds **1.6** and **3.6**, which contain a phenyl substitution at the C-7 position, and compounds **1.8** and **3.8**, both containing a quinoline substitution at the C-7 position, showed the best inhibitory activity and an improvement from their parental compounds containing Br at the C-7 position (Table 2). However, removal of the C-4 and C-8 dimethyl groups led to a reduction of inhibitory activity in each case. For the compounds whose optimal testing concentrations are higher than 25 µg/mL, the inhibitory activity was tested again. At concentration of 100 µg/mL, compounds **1.1, 3.1** and **1.7** showed an increase in their inhibitory activity to 1.79, 1.81 and 2.03 (P values are 0.002, 0.002 and 0.001), but compound **3.7** failed to show an improvement.

**Figure 3.**
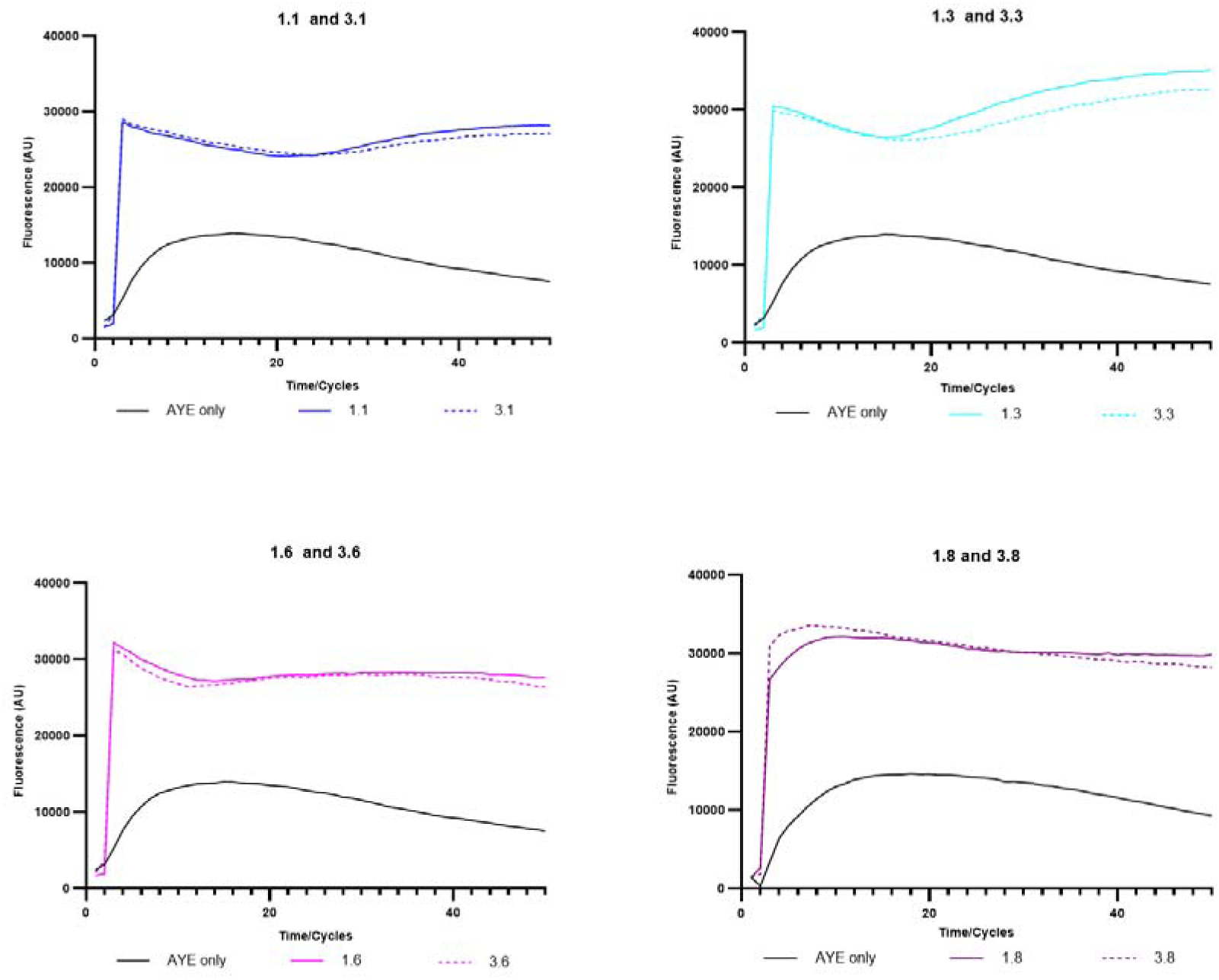
HOECHST DYE accumulation in AYE cells with the addition of some of the most active second-generation quinoline EPI compounds. All results were performed in triplicates and the curve presented is the fluorescence value of three biological repeats after being blanked against cell free PBSM+G with HOECHST DYE; for clarity error bars are not shown on the graph, but the SD is included in the end-point measurements shown in table 2.

**Table 2.** The inhibitory activity of the parental compounds (**1** and **3**) and the second-generation compounds with (25µg/mL) against strain AYE as assessed by Hoechst accumulation.

| *Compound* | *Inhibitory activity* | *P value* | *Compound* | *Inhibitory activity* | *P value* |
| --- | --- | --- | --- | --- | --- |
| **1** | 1.65±0.10 | 8.5E-04 | **3** | 1.86±0.05 | 7.83E-06 |
| **1.1** | 1.11±0.03 | 0.16 | **3.1** | 1.06±0.05 | 0.33 |
| **1.2** | 1.86±0.07 | 0.0007 | **3.2** | 2.07±0.04 | 0.0004 |
| **1.3** | 1.71±0.09 | 0.0007 | **3.3** | 1.93±0.14 | 0.014 |
| **1.4** | 1.93±0.08 | 0.002 | **3.4** | 1.82±0.05 | 6.6E-05 |
| **1.6** | 2.14±0.04 | 0.0001 | **3.6** | 2.11±0.07 | 0.004 |
| **1.7** | 1.60±0.03 | 0.0016 | **3.7** | 1.10±0.06 | 0.18 |
| **1.8** | 2.22±0.07 | 0.0012 | **3.8** | 2.19±0.02 | 0.001 |
| **1.9** | 1.13±0.07 | 0.09 | **3.9** | 1.26±0.03 | 0.009 |
| **1.10** | 1.20±0.05 | 0.03 | **3.10** | 1.06±0.02 | 0.3 |
| **1.11** | 1.24±0.08 | 0.023 | **3.11** | 1.22±0.05 | 0.025 |
| **1.12** | 1.20±0.002 | 0.047 | **3.12** | 1.35±0.02 | 0.035 |

The inhibitory activity was calculated as the fluorescence level of the cells added with EPI compounds divided by that of the EPI-free cells at steady state (after 24min). \*P value: Student t-test was used to calculate the significance, between the cells with and without the EPIs. Values in red are significant ( $P \leq 0.05$ ).

In terms of the structure-activity relationship (SAR), when the compounds were tested at the same concentration, compound **1** accumulated higher fluorescence than the C-7 thiophene-substituted **1.1** and C-7 pyridyl-substituted **1.9**, and similar fluorescence to the C-7 naphthalene-substituted **1.7**. This trend was consistent with compound **3** analogues, as compound **3** showed better efflux inhibitory activity than **3.1**, **3.7**, and **3.9** (Table 2). Interestingly, the C-7 benzothiophene-substituted compounds **1.2** and **3.2** showed greater efflux inhibitory activities than both compounds **1** and **3**. Among other C-7 substitutions, quinoline (**1.8**/**3.8**) and phenyl (**1.6**/**3.6**) exhibited the highest fluorescence accumulation in AYE cells. Compound **1.8** showed better activity than the C-7 naphthalene-substituted **1.7**, suggesting that the nitrogen atom in the quinoline ring of **1.8** played a role in efflux inhibition in the Hoechst assay. However, the C-7 pyridine-substituted compound **1.9** did not show better activity than the C-7 phenyl-substituted **1.6**, indicating that either the ring size, the position of the nitrogen atom within the ring, or both are important for efflux inhibition activity in this assay for quinoline-type compounds. While most second-generation compounds had C-7 aromatic or heteroaromatic rings directly connected to the C-7 position of the quinoline ring, a flexible two-carbon linker was introduced in compounds **1.3** and **3.3**, which contained a phenyl substitution at the end of the linker. Both compounds showed good efflux inhibitory activity comparable to that of the parent compounds **1** and **3**.

After the test of efflux pump inhibitory activities of the second-generation compounds, the antibiotic potentiation activity of all the compounds were assessed by using gentamicin and chloramphenicol at their test concentration (Table 3). Two compounds, the flexible C-7 linker containing **3.3** and the C7-phenhyl substituted **3.11,** significantly reduced the MIC of chloramphenicol against the wild-type Ab5075-UW by 8-32 and 4-8-fold respectively. In the AB5075-CHL strain with overexpressed AdeFGH, a much wider range of compounds gave a greater than 4-fold potentiation. Notably, two pairs of compounds, the C-7 flexible linker containing **1.3** (64-fold), and **3.3** (32-64-fold) and the C7- quinoline substituted **1.8** (16-fold), **3.8** (16-32-fold), that showed highest accumulation in Hoechst assay, showed consistently higher chloramphenicol potentiation than their parent compounds, **1** and **3** (4-fold and 8-fold respectively). C-7 phenyl substituted compound **3.11** showed similar levels of chloramphenicol potentiation (32-64-fold) but this was not replicated by the same modification on the compound **1** scaffold (4-8-fold potentiation for **1.11**). None of the compounds reduced the MIC of gentamicin in either strain AYE or Ab5075-UW, suggesting the structural modifications had selectively improved inhibitory activity against AdeFGH and had no notable effect on inhibition of AdeABC.

**Table 3:**
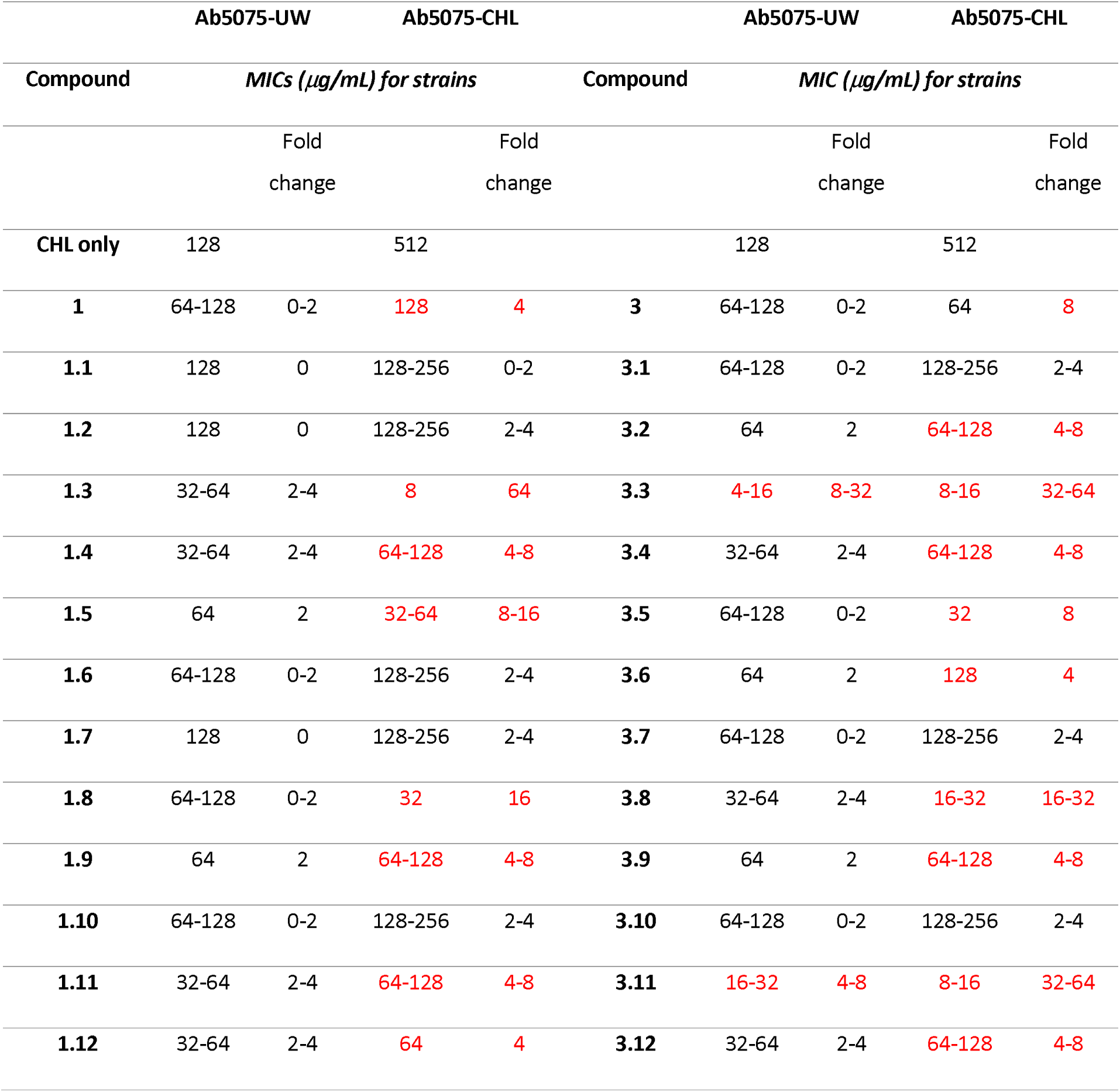
The MICs and fold-reduction for chloramphenicol with the strains Ab5075-UW and Ab50785-CHL-adpated in presence of the second-generation EPIs. The numbers labelled in red colour show the MIC reductions were significant (≥4-fold).

As the interaction of the second-generation compounds with the hydrophobic distal binding pocket of AdeG was considered during the design stage, compounds **1.3** and **3.3**, which showed both strong efflux inhibitory activity in the Hoechst accumulation assay and excellent potentiation of chloramphenicol, were selected for further molecular modelling to study their interactions with the distal binding pocket of AdeG. The flexible C-7 side chain of both compounds effectively interacted with the Phe loop of AdeG, with additional interactions observed with the hydrophobic isoleucine and leucine residues within the binding pocket (Figure 4). The results reinforce the importance of efflux pump inhibitors interacting with the hydrophobic Phe loop within the distal binding pocket of RND-type efflux pumps.

**Figure 4.**
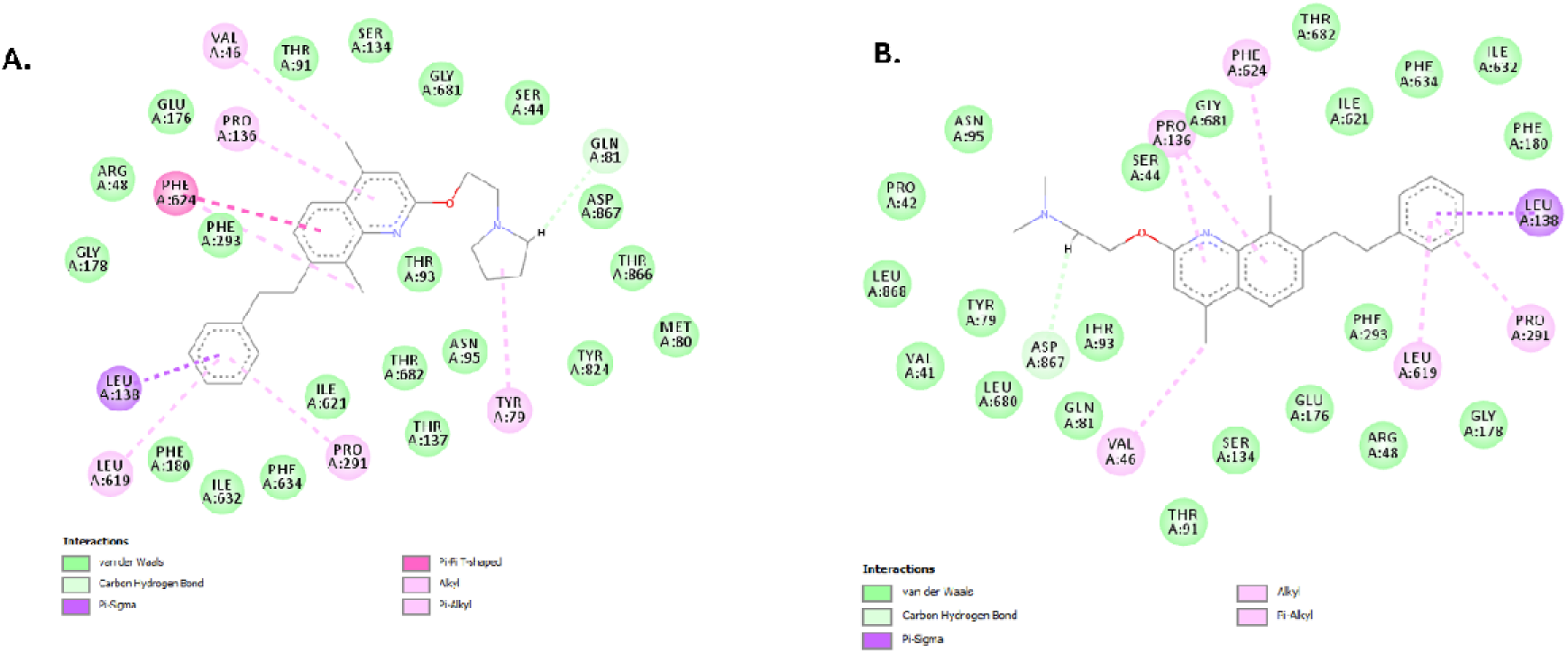
The interaction of 1.3 (A) and 3.3 (B) with the distal binding pocket of AdeG shows that the C-7 substituted group interacts with the Phe loop and hydrophobic residues.

One of the key observations from the study is the discrepancy between the results obtained from the Hoechst dye accumulation assay and the antibiotic potentiation assay. The difference in the results between this assay and the antibiotic potentiation assay suggests that the quinoline compounds may inhibit other efflux pumps besides AdeFGH. This indicates that the reduced efflux of Hoechst dyes observed in the assay may not solely be due to the inhibition of AdeFGH but rather a combination of the inhibition of multiple efflux pumps presents in the bacterial strains used. Additionally, Hoechst assay is also a much more sensitive measurement of efflux than MIC, which is a blunt measurement that is influenced by various factors.

Despite the limitations of the Hoechst dye accumulation assay as a direct measure of AdeFGH inhibition, the data indicates that eight pairs of compounds with specific modifications to the quinoline core scaffold showed selective potentiation of the AdeFGH substrate antibiotic chloramphenicol. Notably, these compounds did not show a similar potentiation effect on the AdeABC substrate antibiotic gentamicin. This selective potentiation underscores the specificity of these quinoline-based efflux pump inhibitors (EPIs) towards AdeFGH, suggesting that the quinoline-based compounds reported in this study inhibit the AdeFGH pump strongly enough to result in significant potentiation of its antibiotic substrate.

Furthermore, structural analysis revealed that some second-generation compounds exhibited strong efflux pump inhibitory activity in both the Hoechst dye accumulation assay and the antibiotic potentiation assay. Specifically, compounds **1.8** and **3.8** demonstrated the best efflux pump inhibitory activity, correlating with their structural features. These compounds possessed a hetero non-aromatic group (either pyrrolidine or dimethyl amine) and a quinoline moiety at the C-2 and C-7 positions of the core quinoline ring, along with C-4 and C-8 methyl groups. This structure activity relationship information suggests that it is feasible to develop selective inhibitors of RND efflux pumps using quinoline as a core scaffold, which could potentiate clinically used antibiotics that are substrates of these efflux pumps.

Finally, the toxicity of the most active compounds was tested using the *Galleria mellonella* non-animal toxicity assay. None of the compounds showed any notable toxicity at 50 mg/Kg dose level suggesting this quinoline scaffold is non-toxic at the doses studied. The ability of these compounds to selectively potentiate the activity of chloramphenicol, coupled with their low toxicity, supports their potential as therapeutic agents. Further optimization and structural refinement of these compounds could lead to the development of clinically useful efflux pump inhibitors that could be co-administered with antibiotics to combat bacterial resistance.

## Conclusions

Quinoline-type compounds were developed in this study and assessed as EPIs for *A. baumannii* AdeABC, AdeFGH and AdeIJK. Data suggested that the the compounds could inhibit efflux from AdeFGH but not the closely releated pump AdeABC, potentially due to specific interactions with Phe residues within the binding interface. The study allowed us to look at both general factors affecting dye accumulation such as efflux inhibition, especially in isogenic lines with mutations in key regulators or efflux pumps, and the function of specific pumps. Although chloramphenicol has been reported as a substrate for AdeIJK, non-RND efflux systems like AbeM and AbeS [26–28] and a novel pemease-based efflux system (Ab5075-UW;[29]), there was a clear MIC phenotype in this study, which could be linked to a mutation in AdeL. Previous studies have defined a similar phenotype associated with gentamicin resistance in AYE and AB5075 [7].

Although the study did not produce an efflux pump inhibitor capable of potentiation activity of clinically useful antibiotics to below the resistance breakpoint, it did provide valuable information on inhibitor specificity of different but structurally related pumps. This may enable future development of EPIs with broader spectrum of activity against RND-family efflux pumps. The variability in the level of potentiation observed for different compounds, suggests that these compounds, in addition to considering them for further med-chem modification to obtain potential development candidates, can be used as probes for efflux pump SAR studies.

## Materials and Methods

**Chemistry:** The solvent and the reagents used for the synthesis were purchased from various commercial sources, including Sigma-Aldrich, Fisher Scientific, and Fluorochem. ^1^H and ^13^C nuclei nuclear magnetic (NMR) analyses were performed on a Spectrospin 400 MHz spectrometer (from Bruker) by using deuterated solvents for the preparation of the samples. The spectra of each compound were analysed using Topspin 3.5pl7 software (Bruker). The chemical shifts were reported relative to Trimethylsilane (TMS) used as standard (0.00 ppm). Signals were identified and described as singlet (s), doublet (d), t (triplet), or m (multiplets). Coupling constants were shown in Hertz (Hz). High resolution mass spectrometry (HRMS) was carried out on an Exactive HCD Orbitrap mass spectrometer (Thermo Scientific). LC-MS analyses were performed on a Waters Alliance 2695 system, eluting in gradient with a flow rate of 0.5 mL/min using a solvent gradient starting with 5% acetonitrile that was increased to 95% acetonitrile over a 7.5 minutes’ time period (ESI). The analyses were performed on a Monolithic C18 50 X 4.60 mm column (made by Phenomenex). UV detection was performed on a Diode Array Detector. Mass spectra were registered in both the ESI+ and ESI- mode. The synthesis and characterization of the quinoline based EPIs reported in this paper can be found in the Supporting Information.

### Bacterial strains and culture conditions

All strains (Table S1) were grown in tryptic soy broth (TSB) (SIGMA) or on tryptic soy agar (TSA) (Biomerieux) at 37°C unless otherwise stated. All the chemicals used in this study were purchased from Sigma unless otherwise stated. All antibiotics stock solutions were dissolved in water to the stock concentration except for chloramphenicol (100% ethanol) and ciprofloxacin (diluted acetic acid). Upon use, they were diluted to the desired concentration in TSB. All quinoline-based compounds tested in this study were dissolved in DMSO at a concentration of 10 mg/mL to make the stock solution, and they were diluted to the desired concentration in TSB before using in the test. CCCP and PAβN stock solutions were made in DMSO and dH_2_O, respectively, and diluted in TSB.

### Minimum inhibitory concentration (MIC) test

A broth-microdilution method was used in accordance with the methodology laid out by the European Committee on Antimicrobial Susceptibility Testing (EUCAST) with modification as described previously.[30] All MICs were performed in TSB media using polystyrene 96-well plates (Corning, Flintshire UK) except for the colistin, where polypropylene plates (Griener Bio-One Ltd, Stonehouse UK) and non-cation adjusted Mueller Hinton media were used. Bacteria were grown in media overnight at 37°C, 200 RPM. They were then diluted to a concentration of 1x10^6^ CFU/mL (OD_600_ = 0.01) and antibiotics added as a 2-fold dilution series, either alone or in combination with potential EPIs. The plate was then incubated at 37 °C statically for 20 hrs after which the absorbance at 600nm was read. After background subtraction, the lowest concentration of antibiotic where OD_600_ value was below 0.1 was considered as the MIC value. All results were carried out at least in triplicates. Bacterial growth in the presence of compounds was monitored by taking an OD_600_ reading every hour for 20 hrs using a FLUOstar Omega plate reader (BMG Labtach GMBH, Ortenberg, Germany).

### Hoechst assay

Hoechst Dye (H33342 bisbenzimide) accumulation assay to measure efflux in *A. baumannii* strains, was carried out as described previously with the following modifications.[25] Compounds to be tested were diluted in DMSO to a concentration 25 times the test concentration. Bacteria were grown in TSB media overnight at 37°C, 200 RPM. 250 µL of the overnight culture was added to 5 mL fresh TSB media, which was then further incubated at 37°C, 200 RPM until the OD_600_ reached 0.4-0.6. Bacterial cells were harvested by centrifugation at 4500 g for 10 mins at room temperature. The supernatant was discarded, and the cell pellet was resuspended PBSM+G (PBS buffer with 20 mM glucose and 1 mM MgSO_4_. The OD_600_ of the cell suspension was measured and adjusted to 0.5. Black bottom 96-well plate (Corning, Flintshire UK) was used in this experiment. In each well, 176 µL of cell suspension, together with 4 µL of compound stock solution, 4 µL of DMSO or 4ul media were added. The plate was incubated for 15min at 37°C in the plate reader before 20 µL of 25μM HOECHST DYE stock solution in dH2O, was injected into each well except for the blank controls to give a final well volume of 200 µL. The fluorescence level (excitation and emission filter at 355 nm and 460 nm) was measured every 2.6 mins for 133 min after the addition of the HOECHST dyes. Hoechst dye has been found to adsorbs on polytetrafluoroethylene-coated material in aqueous solvent, with a concomitant fluorescnece drop later in the Hoechst assay. [31] To mitigate this effect the steady fluorescence level at 24 min after the addition of the Hoechst dye was analysed, rather than the endpoint at 133 min.

### Development of the antibiotic-resistant strains

A gradient plate method was used to select for resistant *A. baumannii* strains to three different antibiotics (gentamicin, chloramphenicol, ceftazidime).[32], known to be substrates for the three efflux pump systems. To adapt each strain to each antibiotic, 20 mL of molten TSA containing no antibiotic was set in a standard petri dish where one side of the plate was elevated 1 cm to allow a slope to form (Figure S2). After the slope had set, 20 mL of TSA containing antibiotic at a concentration of 4-fold the MIC level was added to the plate and allowed to set. Plates were rested overnight at room temperature to ensure diffusion of the antibiotic. 100 mL of bacterial from overnight culture was inoculated on each plate, and the plates were incubated overnight at 37°C. Four colonies nearest to the zone of inhibition from each plate were picked and re-streaked onto a fresh TSA plate with no antibiotic for storage. From the storage plate, MIC of individual colonies were tested. The whole procedure was repeated at a higher concentration across the gradient until a significant increase of MIC (≥ 4-fold) was obtained. Clones of interest were sent for whole genome sequencing to identify mutations arising.

### Whole Genome sequencing and analysis

Bacterial DNA of both wildtype and mutants were purified with a Wizard genomic DNA purification kit (Promega, Wisconsin, US). DNA was then tagged and multiplexed with the Nextera XT DNA kit (Illumina, San Diego, US) and sequenced by Public Health England Genomic Services and Development Unit, (PHE-GSDU) on an Illumina (HiSeq 2500) with paired-end read lengths of 150 bp. A minimum 150 Mb of Q30 quality data were obtained for each isolate. FastQ files were quality trimmed using Trimmomatic.[33] SPAdes 3.1.1 was used to produce draft chromosomal assemblies, and contigs of less than 1 kb were filtered out. [34] FastQ reads from selected isolates were mapped to their respective parental strain pre-exposure chromosomal sequence using BWA 0.7.5. [35]Bam format files were generated using Samtools, [36] and VCF files were constructed using GATK2

Unified Genotyper (version 0.0.7). [37] They were further filtered using the following filtering criteria to identify high-confidence SNPs: mapping quality >30; genotype quality >40; variant ratio >0.9; read depth >10. All the above-described sequencing analyses were performed using PHE Galaxy. [38] BAM files were visualized in Integrative Genomics Viewer (IGV) version 2.3.55.[39] Whole genome alignment and phylogenetic tree generation were performed using progressive alignment in Mauve Version 20150226 build 10. Tree visualisation was performed in FigTree Version 1.4.3.

### Molecular Docking

Since the structures of the AdeB and AdeG have not been validated, the molecular modelling work was based on the models of AcrB and MexB, respectively. Firstly, the amino acid sequence of AdeB (B7I7F7), AcrB (P31224), AdeG (A0A090C131) and MexB (P52002) were downloaded from Uniprot. (Table 7.4) AdeB and AcrB, were 50.36% identical whilst AdeG and MexB were 41.97%. The crystal structures of efflux pump AcrB (1IWG) and MexB (3W9J) were downloaded from the Protein Data Bank (http://www.rcsb.org/) and used as the basis for homology modelling of AdeB and AdeG using Swiss-model. [40] Any missing parts of the predicted AdeB and AdeG structure were amended by using the Biovia Discovery studio visualizer. [41]Finally, the programme Amber was used to minimize energy of the structures. [42, 43] AutoDock SMINA, [44]which fuses the AutoDock Vina scoring function by default, was firstly applied for the blind molecular docking of potential EPI compounds to each structure. This identified the best binding site in the target by exploring all possible binding cavities of the transporter. SMINA was performed with default settings, which samples nine ligand conformations using the Vina docking routine of stochastic sampling. Afterwards, GOLD molecular docking was used to dock of the compounds to the best binding site that located by SMINA of the efflux pump for performing flexible molecular docking.[45] Based on the fitness function scores and ligand binding positions, ten best-docked poses for the compounds were selected. Among the ten poses, the higher fitness function score of poses, generated using the GOLD program that has the more negative GOLD fitness energy value, reveals the best-docked pose for each compound.

### Galleria mellonella survival test

Wax moth larvae (*Galleria mellonella* ) were kept on wood chips at 14°C in the dark until use. For experiments, it was assumed that each *Galleria* had a haemolymph volume of 50 µL and can maximum tolerant 10 μL of liquid injection. Therefore, the stock solution of each compound was prepared at 6 times of the concentration they were going to be tested. For each compound at each test concentration, 10 µL of compound stock solution was injected into 10 *Galleria* via the foremost proleg using a Hamilton syringe. Ten of the control larvae were injected with 10 μL of PSB to control for potential lethal effects from the injection process. After injection, larvae were kept at 37°C inside the petri dishes, and the number of live larvae was recorded every 24 hrs for 5 days. This method was adapted from Want, *et al*. [46]

## ASSOCIATED CONTENT

### Supporting information

Supporting information is available containing: The collection of bacterial strains, gene primers, MIC data, LC-Ms method, experimental details for the synthesis of quinoline EPIS, NMR spectra and HRMS, Molecular formula strings (CSV)

### Notes

The authors declare no competing financial interest.

### Author Contributions

YZ, CKH, MC and KMR carried out experiments, YZ, TAA, MEW and KMR analysed data, CKH, JMS and KMR supervised the experiments, YZ and KMR wrote the manuscript, All authors contributed to the edit and review of the manuscript. All authors agreed to the final version of the manuscript.

## ACKNOWLEDGMENT

This work was supported by an Industrial PhD award for Yiling Zhu (Project ID 537242), a research grant from Medical Research Council (MR/W018594/1) and through UKHSA Grant in Aid Project 109502 and 109505. We also gratefully acknowledge the provision of transposon library strains, as described in Gallagher *et al* 2015 from the Manoil lab[47].

---

**Scheme 1.**
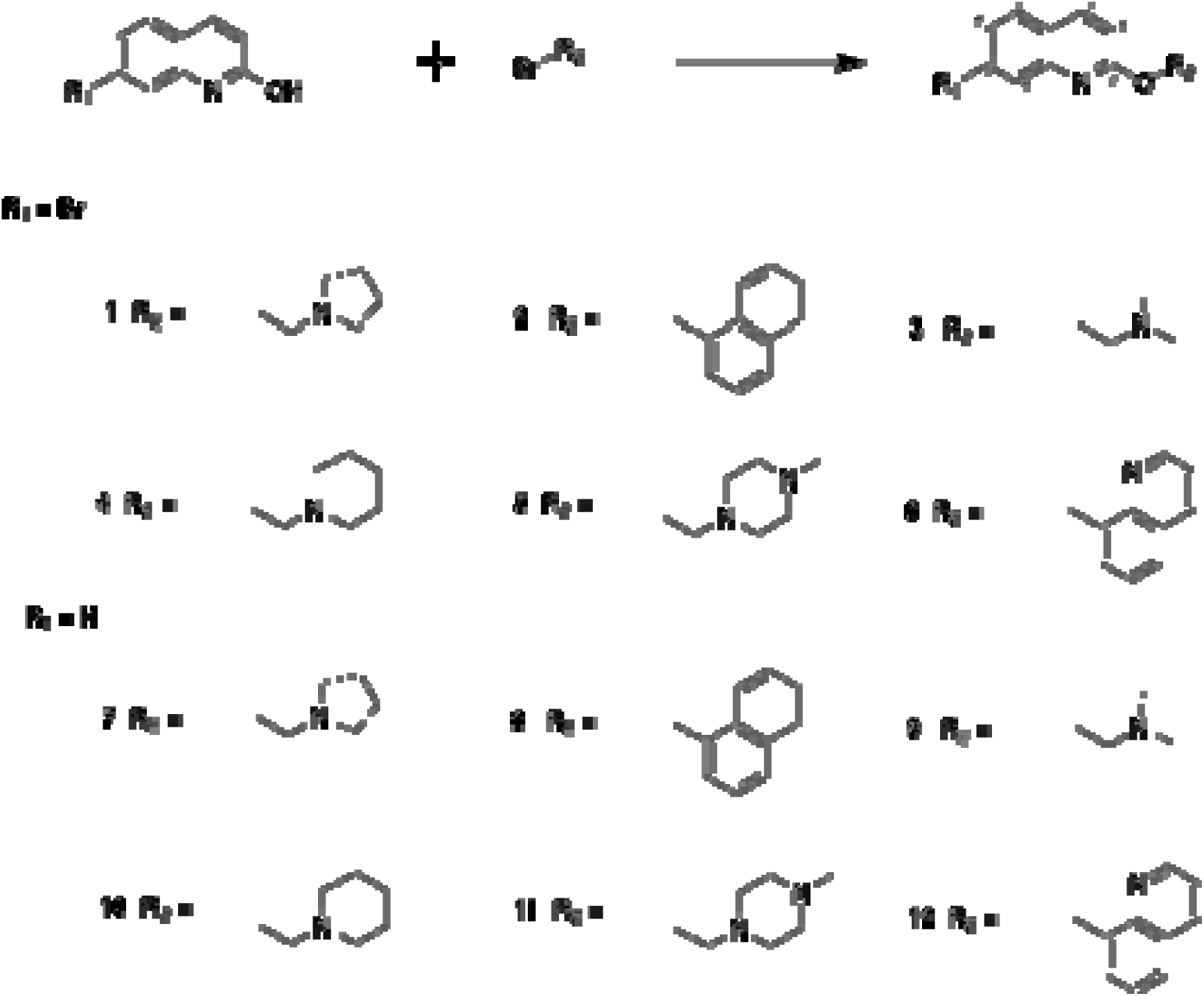
General reaction scheme for synthesis of the first-generation compounds. To synthesize the 12 compounds, three different conditions were used. Condition 1: K_2_CO_3_, Acetone, reflux overnight (for compounds **1**, **2**, **6**); Condition 2: K_2_CO_3_, DMF, reflux overnight (for compounds **3**, **7**); Condition 3; K CO , DMF, microwave 170°C for 30min (for compounds **4**, **5**, **8**, **9**, **10**, **11**, **12**).

**Scheme 2.**
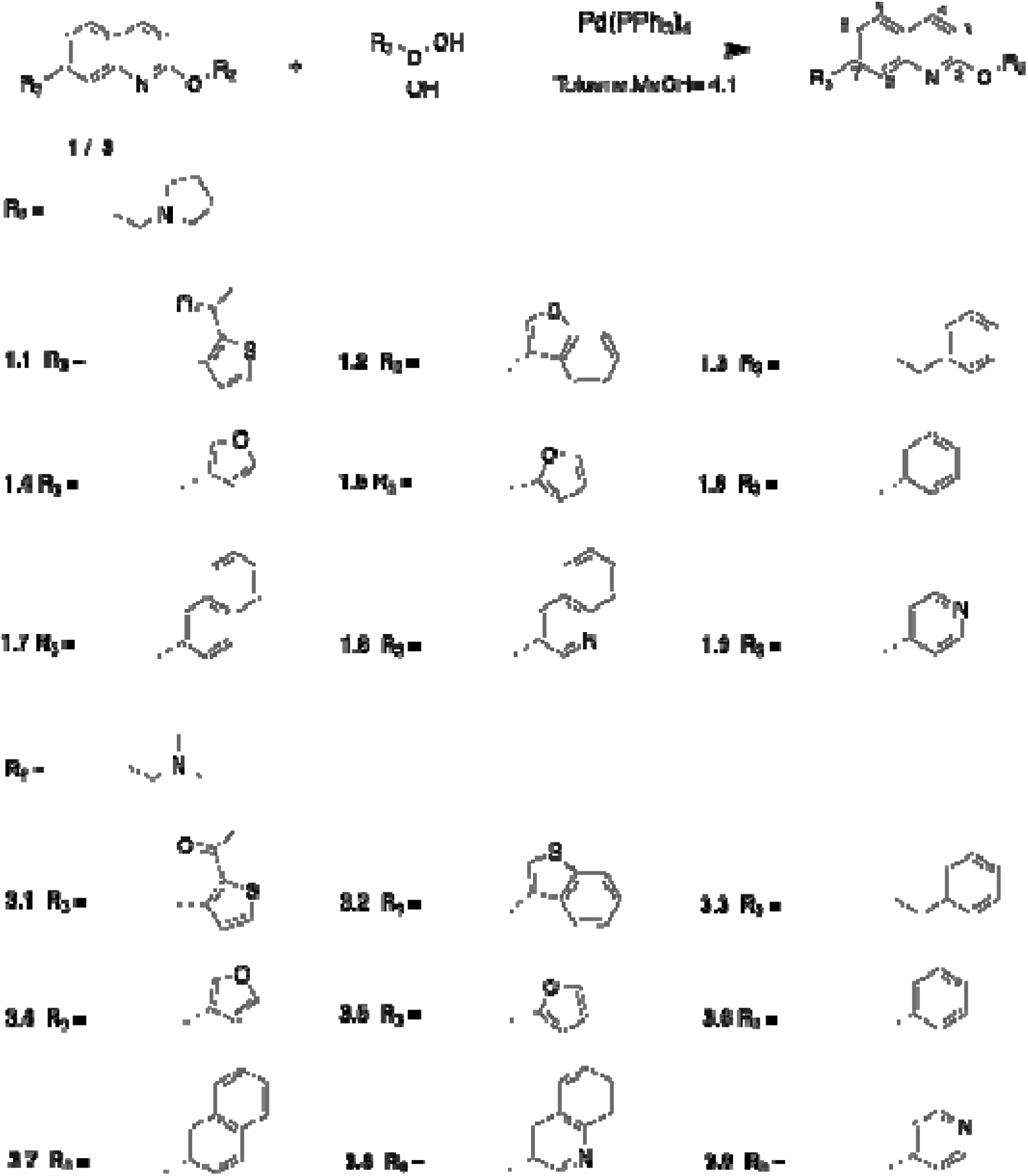
Synthesis of the compounds with C-4 and C-8 methyl groups (1.1 - 1.9, 3.1 - 3.9). Condition: Pd(PPh_3_)_4_ (10% mol of 1 / 3), Toluene/MeOH (4:1), reflux for 6h/overnight.

**Scheme 3.**
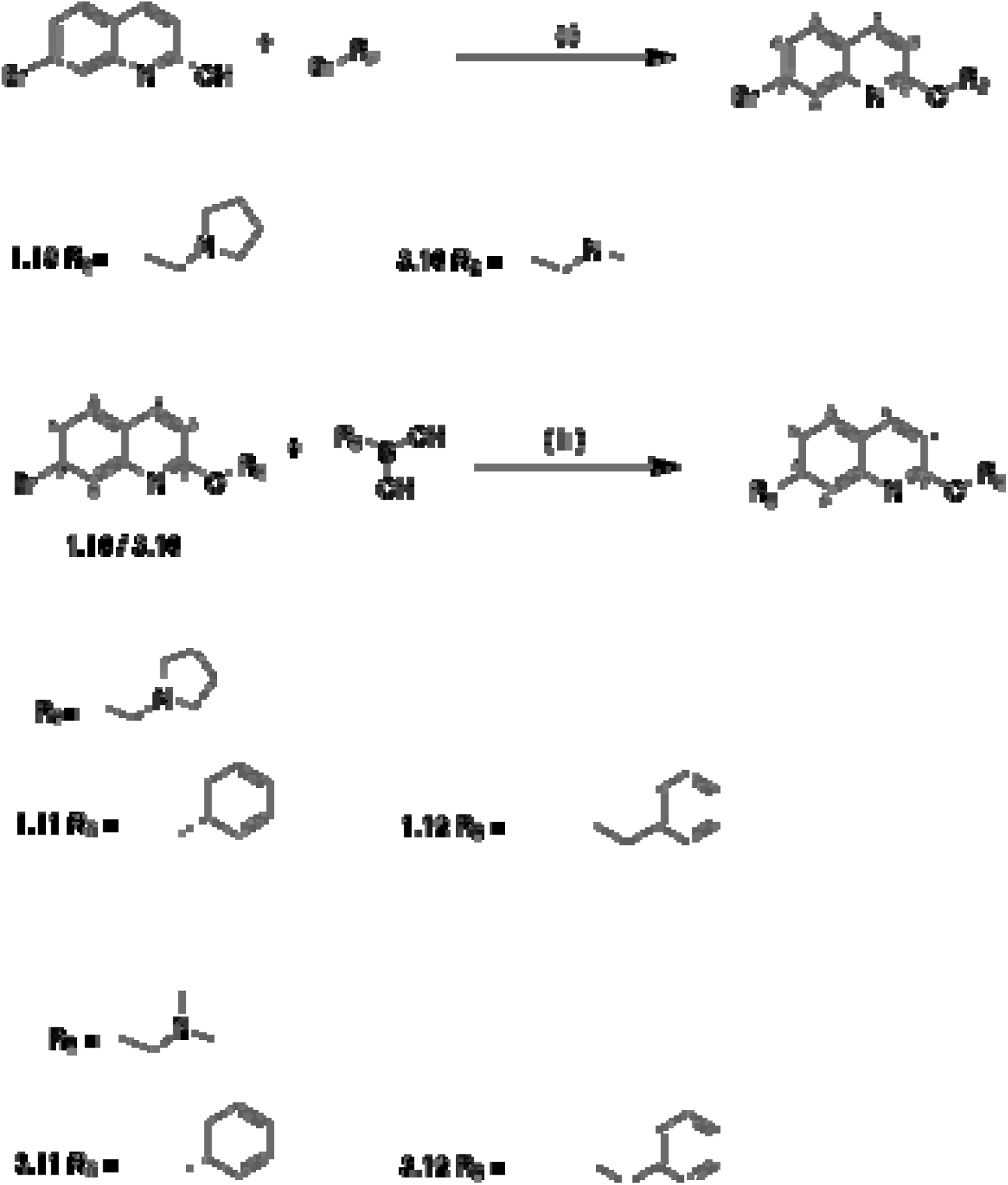
Synthesis of compound without C-4 and C-8 methyl groups (1.10 - 1.12, 3.10 - 3.12). Reagent and conditions: (I) DMF, K_2_CO_3,_ Microwave (60 min). (ii) Pd(PPh_3_)_4_ (10% mol of 1.10 / 3.10), Toluene: MeOH= 4:1 (5ml), reflux overnight.

## Notes

### Competing Interest Statement

The authors have declared no competing interest.

