## Supplementary Information for "C7-substituted Quinolines as Potent Inhibitors of AdeG Efflux Pumps in *Acinetobacter baumannii*"

***Corresponding authors:**

**Supporting Information**

**Supplementary Methods**

**Q-PCR method and primers for checking overexpression of pump genes**

Real-time PCR (RT-PCR) was used to measure the expression of *adeB, adeR, adeS, adeG, adeL, adeJ,* and *adeN* in the Ab5075 transposon mutants, chloramphenicol adapted and Ab5075-UW W/T strains. The method was adapted from *Wand et al*.[32] The primers used for each gene were listed in Table s2. The primer efficiency was firstly checked. In the test, 50 µL of PCR reaction, containing 1µL of template DNA; 0.5 µL of each primer, 25 µL of GoTaq® reaction buffer and 23 µL dH_2_O, was set up. After obtaining the PCR products, they were diluted to 10^-2^, 10^-4^, 10^-5^, 10^-6^, 10^-7^, 10^-8^, 10^-9^, 10^-10^, 10^-11^. For each set of primers, 7 dilution products from 10^-5^ to 10^-11^ were used in the efficiency check test. Seven reactions of 20 µL mixture, each containing 10 µL of SYBR green master mix, 0.2 µL of each 10mM primer stock, 6 µL of PCR product dilution and 3.6 µL dH_2_O, were set up. For each set of primers, 1 no-template control was involved, in which, 6 µL of diluted PCR product was replaced with 6 µL dH_2_O. Each reaction mixture was loaded into a 96-multiwell PCR plate (SIGMA) before amplifying using the StepOnePlus real-time PCR system. The primer efficiency for each set of primers was calculated from by the programme included in the system. (Tabel S7)

After the primer efficiency being checked, the RT-PCR of each gene was carried out. Bacterial overnight cultures were diluted in TSB to an OD_600_ of 0.1, and further incubated till reaching the mid-log phase (OD_600_ = 0.5) and cells were then harvested by using the RNA protect bacteria reagent (Qiagen). Afterwards, RNA of each bacterial strain was extracted by using the RNeasy mini kit (Qiagen), including on-column DNase treatment according to the manufacturer’s instructions. Additionally, 5μg RNA was treated with a DNA-free kit (Ambion), of which 0.2 μg RNA was reverse transcribed by using the SuperScript III first-strand synthesis system for RT-PCR (Invitrogen) according to the manufacturer’s instructions. qPCR was then repeated for at least three times on each sample using a StepOnePlus real-time PCR system and Fast SYBR green master mix (Life Technologies). After the RT-PCR was finished, data were analysed with the Expression Suite Software version 1.0.3 (Life Technologies) by using the *recN, proC* and *fabD* as endogenous controls and taking primer efficiency into account.

**Statistical analysis.**

For comparison of EPI inhibitory activity between experiment added with certain EPI and the EPI-free control, or between different two different EPIs, student's unpaired *T*-test was utilized. A *P* value of <0·05 was considered significant. All statistical analyses were performed on three or more independent experiments using Microsoft Excel.

**Tables**

Table S1. The collection of bacterial strains that were used in this study.

| ***Species*** | ***Strain*** | ***Genotype/mutation*** | ***Strain name used in this study*** | ***Source / Reference*** |
| --- | --- | --- | --- | --- |
| *A. baumannii* | NCTC13421 | AYE | AYE | NCTC |
| *A. baumannii* | AYEΔ*adeB* | ∆*adeB* mutant | AYEΔ*adeB* | Richmond et al 2016 [7] |
| *A. baumannii* | AYEΔ*adeRS* | ∆*adeRS* mutant | AYEΔ*adeRS* | Richmond et al 2016 [7] |
| *A. bumannii* | Ab5075-UW | Ab-5075 W/T | Ab5075-UW | Gallagher et al 2015 [47] |
| *A. baumannii* | Ab-5075 Cam | Ab-5075 chloramphenicol adapted mutant | Ab5075-CHL | This study |
| *A. baumannii* | ABUW_1975 | *adeB* transposon mutant | Ab5075Δ*adeB* | Gallagher et al 2015; [47] |
| *A. baumannii* | ABUW_1973 | *adeR* transposon mutant | Ab5075Δ*adeR* | Gallagher et al 2015; [47] |
| *A. baumannii* | ABUW_1972 | *adeS* transposon mutant | Ab5075Δ*adeS* | Gallagher et al 2015; [47] |
| *A. baumannii* | ABUW_1335 | *adeG* transposon mutant | Ab5075Δ*adeG* | Gallagher et al 2015; [47] |
| *A. baumannii* | ABUW_1338 | *adeL* transposon mutant | Ab5075Δ*adeL* | Gallagher et al 2015; [47] |
| *A. baumannii* | ABUW_1336 | *adeJ* transposon mutant | Ab5075Δ*adeJ* | Gallagher et al 2015; [47] |
| *A. baumannii* | ABUW_1731 | *adeN* transposon mutant | Ab5075Δ*adeN* | Gallagher et al 2015; [47] |

Table S2. The MICs of different antibiotics on AYE, Ab5075 and their mutants show effects of AdeABC and its regulator AdeRS on efficacy of gentamicin and ciprofloxacin.

| **Antibiotic** | **MIC(μg/mL) for strain** | | | | | | | | | | |  |
| --- | --- | --- | --- | --- | --- | --- | --- | --- | --- | --- | --- | --- |
|  | **AYE** | **AYE**  **Δ*adeB*** | **AYE**  **Δ*adeRS*** | **Ab5075-UW** | **Ab5075**  **Δ*adeB*** | **Ab5075**  **Δ*adeR*** | **Ab5075**  **Δ*adeS*** | **Ab5075**  **Δ*adeG*** | **Ab5075**  **Δ*adeL*** | **Ab5075**  **Δ*adeJ*** | **Ab5075**  **Δ*adeN*** | **Ab5075-CHL-adapted** |
| **GENT** | ≥1024 | 32-64 | 32-128 | ≥1024 | 32-64 | 32-64 | 128-256 | ≥1024 | ≥1024 | ≥1024 | >1024 | 512 |
| **RIF** | 32 | 16 | 16-32 | 1-4 | 2-16 | 2 | 1 | 2 | 1 | 0.5-2 | 2 | N/A |
| **CLR** | 32-64 | 8-32 | 32 | 16-64 | 8-32 | 8-64 | 16 | 16 | 32 | 16-32 | 16 |  |
| **CHL** | 256-512 | 128-256 | 256-512 | 64-128 | 64 | 128 | 128 | 128 | 128 | 128 | 128-256 | 512 |
| **IME** | 1-2 | 1 | 1 | 16-32 | 16 | 16 | 16 | 32 | 16 | 16 | 32 | N/A |
| **MER** | 1-2 | 1-2 | 1 | 32 | 32 | 32 | 32 | 64 | 32 | 32 | 32 |  |
| **CEFO** | >1024 | 1024 | >1024 | >1024 | 1024 | >1024 | >1024 | >1024 | >1024 | >1024 | >1024 |  |
| **CIP** | 128 | 32 | 32 | 64 | 16 | 32 | 32 | 64 | 64 | 32 | 128 |  |

GENT = gentamicin; RIF=rifampicin; CLR=clarithromycin; CHL= chloramphenicol; IME= imipenem; MER= Meropenem; CEFO= cefotaxime; CIP= ciprofloxacin. Numbers shown in red have a reduction in MIC of at least 4-fold for the mutants compared to the corresponding wildtype strain. N/A not tested.

Table S3. The ΔG and ChemScore for compounds 1 and 3 in the best poses against AdeB and AdeG

| **Compounds** | **AdeB** | | | **AdeG** | |
| --- | --- | --- | --- | --- | --- |
|  | **Chem Score** | **∆G kcal/mol** | | **Chem Score** | **∆G kcal/mol** |
| **1** | 29.28 | | -32.95 | 28.76 | -37.42 |
| **3** | 29.45 | | -30.73 | 30.05 | -30.34 |

Table S4. The test concentration selected for all synthesised EPIs on AYE and Ab5075-UW based on less than 10% reduction in growth compared to control.

| ***Compound Name*** | ***Conc. tested (μg/mL)*** | ***Compound Name*** | ***Conc. tested (μg/mL)*** |
| --- | --- | --- | --- |
| **1** | 25 | **7** | 25 |
| **2** | 25 | **8** | 50 |
| **3** | 50 | **9** | 50 |
| **4** | 25 | **10** | 25 |
| **5** | 12.5 (25 for Ab5075-UW) | **11** | 50 |
| **6** | 100 | **12** | 100 |
| **1.1** | 100 | **3.1** | 100 |
| **1.2** | 100 | **3.2** | 6.25 |
| **1.3** | 25 | **3.3** | 25 |
| **1.4** | 25 | **3.4** | 25 |
| **1.5** | 25 | **3.5** | 25 |
| **1.6** | 12.5 | **3.6** | 12.5 |
| **1.7** | 100 | **3.7** | 100 |
| **1.8** | 25 | **3.8** | 25 |
| **1.9** | 100 | **3.9** | 100 |
| **1.10** | 50 | **3.10** | 50 |
| **1.11** | 50 | **3.11** | 50 |
| **1.12** | 50 | **3.12** | 50 |
| **PAβN** | 25 | **CCCP** | 10 |

Table S5. The MICs of chloramphenicol but not gentamicin is reduced in the presence of compound **1** and **3** treatment in a strain overexpressing AdeFGH.

| **Antibiotic** | **EPI** | **AYE** | **Ab5075-UW** | **Ab5075-CHL-adapted** | **Ab5075*ΔadeG*** | **Ab5075*ΔadeL*** |
| --- | --- | --- | --- | --- | --- | --- |
|  |  | ***MICs(μg/mL) for strains*** | | | | |
| **GENT** | No EPI | >1024 | ≥1024 | 512 | 512 | 512 |
|  | **1** | ≥1024 | 512 | N/A | | |
|  | **3** | ≥1024 | 512 |  |  |  |
|  | **4** | ≥1024 | 512 |  |  |  |
|  | **PAβN** | ≥1024 | 512 | 512 | 512 | 512 |
|  | **CCCP** | 1024 | 512 | 512 | 512 | 512 |
| **CHL** | No EPI | 256-512 | 128 | 512 | 128 | 128 |
|  | **1** | 256 | 64-128 | 128 | 128 | 128 |
|  | **3** | 128 | 64-128 | 64 | 128 | 128 |
|  | **4** | 256 | 64-128 | 256 | 128 | 128 |
|  | **PAβN** | 128 | 64-128 | 128-256 | 64 | 64 |
|  | CCCP | 256 | 128 | 256 | 128 | 128 |

All EPIs compounds were tested at conc. of 25μg/mL The number in red colour show a reduction in MIC of at least 4-fold, compared to the untreated control, due to the addition of EPI. . *N/A not tested.

Table S6. The MICs of rifampicin, clarithromycin and colistin for strain AYE in the presence of PAβN and CCCP and compared to three selected EPI compounds **1**,**3** and **4**.

| ***Antibiotics*** | ***EPIs*** | ***MIC (μg/mL) for AYE*** |
| --- | --- | --- |
| **RIF** | No EPI | 32 |
|  | PAβN | 0.125-0.25^*^ |
|  | CCCP | 4 |
|  | **1** | 4-8 |
|  | **3** | 16 |
|  | **4** | 8 |
| **CLR** | No EPI | 32-64 |
|  | PAβN | 0.5^*^ |
|  | CCCP | 16 |
|  | **1** | 32-64 |
|  | **3** | 16 |
|  | **4** | 16 |
| **CST** | No EPI | 0.5 |
|  | PAβN | 0.5 |
|  | CCCP | ≤0.125^*^ |
|  | **1** | ≤0.125^*^ |
|  | **3** | ≤0.125^*^ |

The numbers in red colour show the MIC of reduced for at least 4-fold due to the addition of EPI.

Table S7. The primers and prime efficiency for each gene tested in RT-PCR

| **Gene name** | **S primer** | **AS Primer** |
| --- | --- | --- |
| ***adeB*** | GGATTATGGCGACTGAAGGA | AATACTGCCGCCAATACCAG |
| ***adeR*** | TGCACTAGAGCGAACCGTAG | CTATATCCCACGCCACGCAC |
| ***adeS*** | GAATTCACTCCGCCGAAATGT | AACTCATGTGCGATAGCTGC |
| ***adeG*** | AATCGCGACCGCTTGTTTATT | ACCAGATGGCGCGGTTAC |
| ***adeL*** | GGACAGCCCGTATTTTAGCG | CGTCCAATCGATACAGGCACA |
| ***adeJ*** | CATCGGCTGAAACAGTTGAA | GCCTGACCATTACCAGCACT |
| ***adeN*** | ATGCTGTCTCTCTTGACGACAT | ATCGCAGATTGCAGTAAATAAGCC |

1 cm

1 cm

20mL of TSA agar

Concentration of antibiotic=0μg/mL

Concentration of antibiotic= 4* MIC

20ml of TSA agar with antibiotic at concentration of 4*MIC

Figure S1. The method of making antibiotic gradient plate.

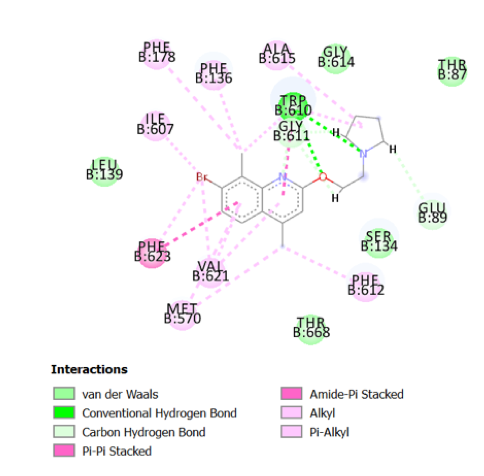

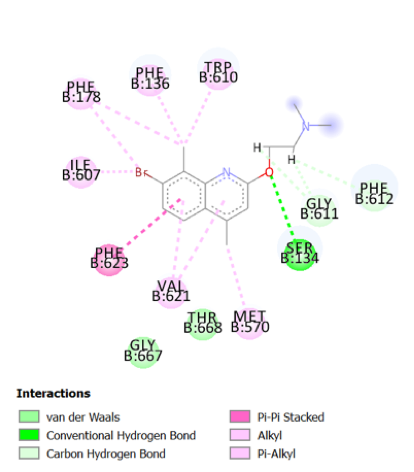

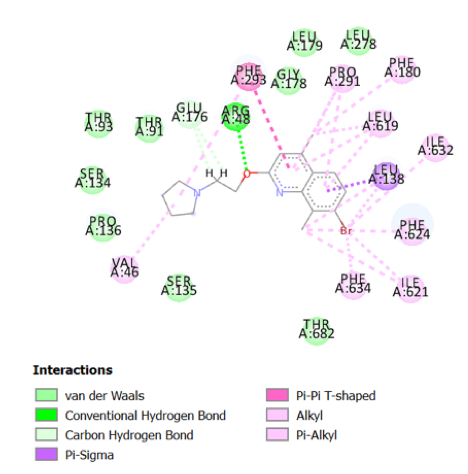

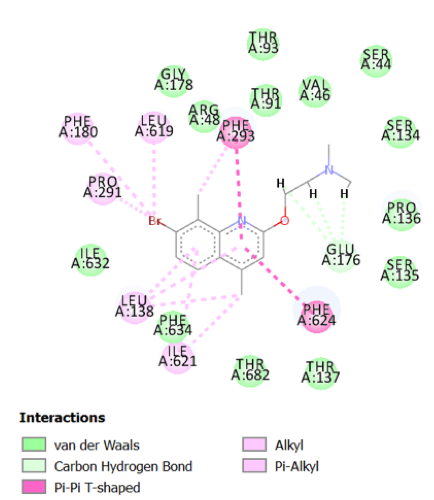

a

b

c

d

Figure S2. The interaction between compound **1** or **3** and key residues in the efflux complexes. (a)**1** and AdeB complexes; (b) **3** and AdeB complexes; (c) **1** and AdeG complexes; (d) **3** and AdeG complexes.

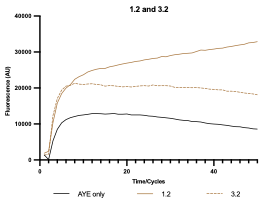

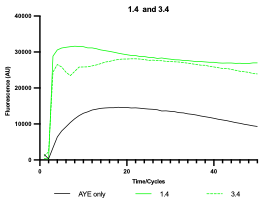

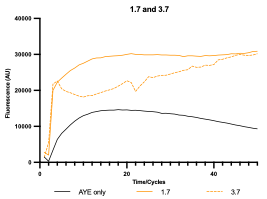

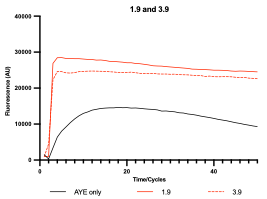

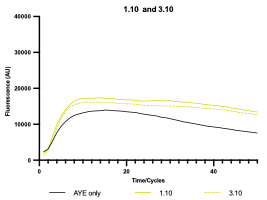

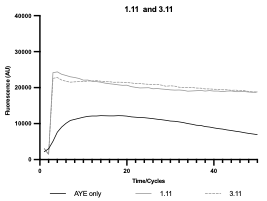

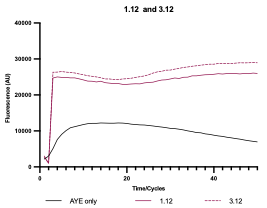

Figure S3. HOECHST DYE accumulation in AYE cells with the addition of 14 synthesised second-generation EPI compounds. The EPI compounds were added to the cell suspension at the maximum concentration which did not adversely affect growth and incubated for 15 min before the addition of HOECHST dyes. Fluorescence levels were then recorded every 2 min 33 sec for 50 cycles. Cells with compounds with the same R_3_ group but different R_2_ are shown in the same colour, with Br substitutions shown as solid lines (R_2_=quinoline), compared to dotted lines for EPIs with no substitution (R_2_=amine). All results were performed in triplicates and the curve presented is the fluorescence value of three biological repeats after being blanked against cell free PBSM+G with HOECHST DYE; for clarity error bars are not shown on the graph, but the SD is included in the end-point measurements shown in table 2.

**Chemistry experimental section**

### Purity determination of synthesized final compounds

The level of purity of the compounds for biological testing has been evaluated through LC-MS analysis, using two different gradient methods, reported hereafter. LC-MS analyses were performed on a Waters Alliance 2695 system (from Waters), with elution in gradient. HPLC grade solvents were used as mobile phase while a Monolithic C18 50 X 4.60 mm column (from Phenomenex) was used as stationary phase. UV detection was performed using a Waters 2996 photo array detector (from Waters). Injection volume has been set to 10 µL. The compounds have been dissolved in a mixture of H2O/ACN (50/50, v/v) or DMSO/ACN (50/50, v/v) accordingly to the solubility. The area of the peak corresponding to the compound has been automatically determined by the software included in the LC-MS system. The eventual presence of solvent UV trace has been subtracted to the total in order to determine the percentage of purity. All compounds showed at least 95% purity in both methods.

LC-MS methods:

Method A: flow 0.5 mL/min

A) water + 0.1 % formic acid

B) acetonitrile + 0.1% formic acid

| Time (min) | 0 | 3 | 3.5 | 4.5 | 5 |
| --- | --- | --- | --- | --- | --- |
| A (%) | 95 | 10 | 5 | 5 | 95 |
| B (%) | 5 | 90 | 95 | 95 | 5 |

Method B: flow 1 mL/min

A) water + 0.1 % formic acid

B) acetonitrile + 0.1% formic acid

| Time (min) | 0 | 2 | 5 | 6 | 7.5 | 9 | 10 |
| --- | --- | --- | --- | --- | --- | --- | --- |
| A (%) | 95 | 95 | 50 | 50 | 5 | 95 | 95 |
| B (%) | 5 | 5 | 50 | 50 | 95 | 5 | 5 |

**7-bromo-4,8-dimethyl-2-(2-(pyrrolidin-1-yl)ethoxy)quinoline (1)**

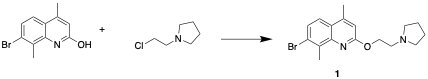

7-bromo-4,8-dimethylquinolin-2-ol (100 mg, 39.7 mmol) was mixed with 1-(2-chloroethyl)pyrrolidine (63.6 mg, 47.6 mmol) and K_2_CO_3_ (164.6 mg, 119.1 mmol), followed by addition of 20 ml acetone. The mixture was heated to reflux overnight or until TLC (DCM/ MeOH 9:1) showed complete consumption of the starting material. Upon cooling, solvent was evaporated. The product was extracted with EtOAc, dried over MgSO_4_ and concentrated by rotary evaporation. Purification by flash column chromatograph was used to afford the title compound (108 mg, 30.9 mmol, yield=78^1^H-NMR (400MHZ, CD_3_OD*,* ppm) δ: 1.9-2.1 (m,4H), 2.56 (s,3H), 2.71 (s,3H), , 3,62 (t, *J*=6.0Hz, 6.0Hz, 2H), 4.72(t, *J*=6Hz,6Hz, 2H), 6.85 (s, 1H), 7.42 (d, *J*=8.8Hz, 1H), 7.65 (d, *J*=9.2Hz,1H). ^13^C-NMR (101MHz, Chloroform-*d,* ppm) δ:161.2, 147.0, 146.0, 135.4, 127.4, 125.7, 124.1, 122.1, 112.9, 64.4, 57.9, 54.9, 54.7, 54.2, 23.5, 22.7, 18.9, 17.8.HR-MS, m/z calc. for C_17_H_21_BrN_2_O (M)^+^ 349.1722 found 349.3711 ([M]+H)^+^.

**7-bromo-4,8-dimethyl-2-(naphthalen-1-ylmethoxy)quinoline (2).**

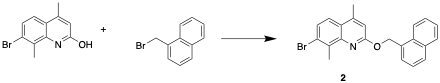

Compound **2** was obtained with 7-bromo-4,8-dimethylquinolin-2-ol (100mg, 39.7mmol) and 1-(bromomethyl)naphthalene (105.2 mg, 47.6 mmol) under the same condition of compound **1**. (56 mg, 14.3 mmol, yield=36%). ^1^H-NMR (400MHZ, DMSO-*d_6_,* ppm) δ: 2.59 (s, 3H), 2.80 (s, 3H), 5.99 (s, 2H), 6.98 (s, 1H), 7.52 (t, *J*=7.6Hz, 8Hz, 1H), 7.58 (t, *J*=7.2Hz, 6Hz, 2H), 7.63 (d, *J*=9.2Hz, 1H), 7.75 (d, *J*=8.4Hz, 2H), 7.77 (s, 1H), 7.97 (d *J*=9.6Hz, 1H), 8.13 (d, *J*=2.4Hz, 1H). ^13^C-NMR (101MHz, DMSO-*d_6_,* ppm) δ: 160.8, 148.3, 145.2, 134.1, 133.3, 132.6, 131.3, 128.7, 128.5, 127.5, 127.4, 126.5, 125.9, 125.34, 125.29, 124.0, 123.8, 123.2, 112.7, 65.2, 18.3, 17.6. HR-MS, m/z calc. for C_22_H_18_BrNO (M)^+^ 392.0876 found 392.9897 ([M]+H)^+^.

**2-((7-bromo-4,8-dimethylquinolin-2-yl)oxy)-*N*,*N*-dimethylethan-1-amine (3).**

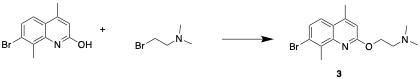

7-bromo-4,8-dimethylquinolin-2-ol (100 mg, 39.7 mmol) and K_2_CO_3_ (164.6 mg, 119.1 mmol) was mixed in 10 mL DMF, and the solution was heated up at 100^o^C for 2 hrs. 2-bromo-*N*,*N*-dimethylethan-1-amine (72.4 mg, 47.6 mmol) was then added into the solution. The mixture was heated to reflux overnight. The reaction was cooled to room temperature and extracted with EtOAc and water. The organic layer was separated, dried over MgSO_4_ and evaporated. The residue was purified by flash column chromatography to obtain the product as white solid (62 mg, 19.2 mmol, yield=48%). ^1^H-NMR (400MHZ, CD_3_OD*,* ppm) δ:2.25 (s, 6H), 2.43 (s, 3H), 2.60 (s, 3H), 2.71 (t, *J*= 5.6Hz, 5.6Hz, 2H), 4.45 (t, *J*=6Hz, 5,6Hz, 2H), 6.65 (s, 1H), 7.34 (d, *J=8*.8Hz, 1H), 7.44 (d, *J*=8.8Hz, 1H). ^13^C-NMR (101MHz, CD_3_OD*,* ppm) δ:162.4, 149.0, 147.0, 136.0, 128.57, 128.60, 125.3, 123.6, 113.7, 63.9, 58.9, 45.86, 18.9, 18.0. HR-MS, m/z calc. for C_15_H_19_BrN_2_O (M)^+^ 322.9765 found 323.7734 ([M]+H)^+^

**7-bromo-4,8-dimethyl-2-(2-(piperidin-1-yl)ethoxy)quinoline (4).**

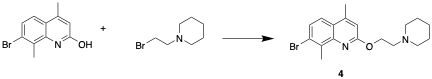

7-bromo-4,8-dimethylquinolin-2-ol (100 mg, 39.7 mmol), 1-(2-bromoethyl)piperidine (91.4 mg, 47.6 mmol) and K_2_CO_3_ (164.6 mg, 119.1 mmol) were mixed in 5 ml DMF in a microwave-proof glass tube. The mixture was heated for 30 mins at 170^o^C in a microwave synthesis. Upon cooling, the reaction was extracted with EtOAc and water. MgSO_4_ was used to dry the separated organic layer. The compound was obtained by flash column chromatography (64 mg, 17.6 mmol, yield=44.3%). ^1^H-NMR (400MHZ, CD_3_OD*,* ppm) δ:1.49-1.50 (m, 2H), 1.62-1.68 (m, 4H), 2.51 (s, 3H), 2.57 (t*, J*=3.2Hz, 5.2Hz, 4H), 2.68 (s, 3H), 2.86 (t*, J*=5.6Hz, 5.6Hz, 3H), 4.57 (t*, J*=5.6Hz, 5.6Hz, 3H), 6.72 (s, 1H), 7.42 (d*, J*=8.4Hz, 1H), 7.52 (d*, J*=8.4Hz, 1H). ^13^C-NMR (101MHz, DMSO-*d_6_,* ppm) δ: 206.5, 160.87, 148.0, 145.2, 134.0, 127.2, 125.2, 123.8, 123.1, 112.7, 54.2, 40.1, 30.7, 25.3, 18.3, 17.4. HR-MS, m/z calc. for C_18_H_23_BrN_2_O (M)^+^ 363.1275 found 364.0189 ([M]+H)^+^

**7-bromo-4,8-dimethyl-2-(2-(4-methylpiperazin-1-yl)ethoxy)quinoline (5).**

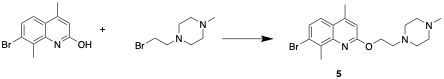

7-bromo-4,8-dimethylquinolin-2-ol (100 mg, 39.7 mmol), 1-(2-bromoethyl)-4-methylpiperazine (98.6 mg, 47.6 mmol) and K_2_CO_3_ (164.6 mg, 119.1 mmol) were mixed in 5 ml DMF in a microwave-proof glass tube. The mixture was heated for 1h at 170^o^C in a microwave synthesis. Upon cooling, the reaction was extracted with EtOAc and water. The organic layer was collected, dried by MgSO_4,_ and concentrated. The product was obtained by flash column chromatography (80 mg, 21.1 mmol, yield=53.1%). ^1^H-NMR (400MHZ, Chloroform-*d,* ppm) δ:2.25 (s, 3H), 2.48 (t, *J*=9.2Hz, 5.6Hz, 4H), 2.51 (s, 3H), 2.60 (t, *J*=2.4Hz, 2.1Hz), 2.72 (s, 3H), 2.81 (t, *J*=6Hz, 5.6Hz), 4.55 (t, *J*=6Hz, 6Hz), 6.69 (s, 1H), 7.44 (d, *J*=8.8Hz, 1H), 7.48(d, *J*=9.2Hz, 1H). ^13^C-NMR (101MHZ, Chloroform-*d,* ppm) δ: 161.1, 147.1, 145.9, 135.4, 127.5, 125.8, 124.1, 122.1, 112.9, 63.1, 57.0, 55.0, 53.4, 50.8, 46.0, 18.9, 17.8. HR-MS, m/z calc. for C_18_H_24_BrN_3_O (M)^+^ 378.1437 found 378.1711 ([M]+H)^+^

**7-bromo-4,8-dimethyl-2-(quinolin-8-ylmethoxy)quinoline (6).**

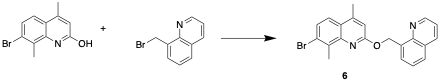

The compound was obtained with 7-bromo-4,8-dimethylquinolin-2-ol (100 mg, 39.7 mmol) and 8-(bromomethyl)quinoline (105.7 mg, 47.6 mmol) under the same condition of compound **1** (62 mg, 15.9 mmol, yield=40%). ^1^H-NMR (400MHZ, Chloroform-*d,* ppm) δ:2.6 (s, 3H), 2.8 (s, 3H), 6.3 (s, 2H), 6.89 (s, 1H), 7.45 (d, *J*=4Hz, 1H), 7.47 (d, *J*=4Hz, 1H), 7.53 (t, *J*=4Hz, 8Hz, 1H), 7.54 (d, *J*=8Hz, 1H), 7.78 (d, *J*=8Hz, 1H), 7.95 (d, *J*=8Hz, 1H), 8.18 (d, *J*=8Hz, 1H), 9.12 (d, *J*=8Hz, 1H). ^13^C-NMR (101MHZ, Chloroform-*d,* ppm) δ:161.2, 149.8, 147.1, 136.3, 135.8, 135.6, 128.6, 128.2, 127.6, 127.5, 126.3, 125.7, 124.2, 122.1, 121.2, 113.0, 63.7, 18.9, 17.9. HR-MS, m/z calc. for C_21_H_17_BrN_2_O (M)^+^ 393.1251 found 394.1243 ([M]+H)^+^

**4,8-dimethyl-2-(2-(pyrrolidin-1-yl)ethoxy)quinoline (7).**

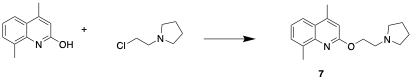

4,8-dimethylquinolin-2-ol (100 mg, 57.7 mmol) and K_2_CO_3_ (239.2 mg, 173,1 mmol) was dissolved in 20 mL DMF, and the solution was heated up at 100^o^C for 2h. 1-(2-chloroethyl)pyrrolidine (92.5 mg, 69.24 mmol) was then added into the solution. The mixture was heated to reflux overnight. Upon finishing, the reaction was cooled to room temperature, and extracted with EtOAc and water. The organic layer was separated, dried over MgSO_4_ and evaporated. The residue was purified by flash column chromatography to obtain the product (30 mg, 11.1 mmol, yield=19.2%). ^1^H-NMR (400MHZ, Chloroform-*d,* ppm) δ: 1.82-1.84 (m, 4H), 2.60 (s, 3H), 2.70 (s, 3H), 2.76 (t, *J*=3.2Hz, 5.2Hz, 4H), 3.25 (t, *J*=6Hz, 6Hz, 2H), 4.70 (t, *J*=6Hz, 6Hz, 2H), 6.81 (s, 1H), 7.29 (t, *J*=5.6Hz, 2Hz, 1H), 7.49 (d, *J*=6.8Hz, 1H), 7.73 (s, *J*=8Hz, 1H). ^13^C-NMR (101MHz, DMSO-*d_6_,* ppm) δ: 162.3, 155.6, 146.0, 136.2, 129.7, 123.6, 122.8, 122.1, 113.5, 66.8, 56.9, 56.52, 56.46, 23.56, 23.62, 19.9, 16.9.HR-MS, m/z calc. for C_17_H_22_N_2_O (M)^+^ 271.2135 found 272.1733 ([M]+H)^+^

**4,8-dimethyl-2-(naphthalen-1-ylmethoxy)quinoline (8).**

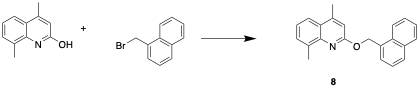

4,8-dimethylquinolin-2-ol (100 mg, 57.7 mmol), 1-(bromomethyl)naphthalene (191.4 mg, 86.6 mmol) and K_2_CO_3_ (160 mg, 115.4 mmol) were mixed in 5 μL DMF in a microwave-proof glass tube. The mixture was heated for 1h at 170^o^C in a microwave ml synthesis. Upon cooling, the reaction was extracted with EtOAc and water. The organic layer was collected, dried by MgSO_4,_ and concentrated. The product was obtained by flash column chromatography (38 mg, 12.1 mmol, yield=21%). ^1^H-NMR (400MHZ, Chloroform-*d,* ppm) δ: 2.48 (s, 3H), 2.69 (s, 3H), 5.90 (s, 2H), 6.69 (s,1H), 7.20 (t, *J*=8Hz, 7.6Hz, 1H), 7.36(t, *J*=8Hz, 7.2Hz, 1H), 7.38 (d, *J*=6.8Hz, 1H), 7.43 (t, *J*=8Hz, 8Hz, 2H), 7.62 (d, *J*=7.2Hz, 2H), 7.74 (d, *J*=8.4Hz, 1H), 7.78 (d, *J*=7.6Hz, 1H), 8.11 (d, *J*=8.4Hz, 1H). ^13^C-NMR (101MHz, Chloroform-*d,* ppm) δ: 160.6, 147.2, 145.3, 135.8, 133.8, 133.3, 132.1, 129.6, 128.9, 128.7, 127.6, 126.4, 125.8, 125.4, 125.3, 124.2, 121.5, 112.8, 65.7, 19.1, 18.4. HR-MS, m/z calc. for C_22_H_19_NO (M)^+^ 314.1333 found 313.1121 ([M]+H)^+^

**2-((4,8-dimethylquinolin-2-yl)oxy)-*N*,*N*-dimethylethan-1-amine (9).**

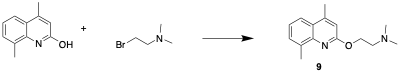

4,8-dimethylquinolin-2-ol (100 mg, 57.7 mmol) and 2-bromo-*N*, *N*-dimethylethan-1-amine (131.7 mg, 86.6 mmol) was used to afford the product under the same condition of**.8**. (20 mg, 8.3 mmol, yield=14.3%). ^1^H-NMR (400MHZ, Chloroform-*d,* ppm) δ: 2.32 (s, 6H), 2.52 (s, 3H), 2.61 (s, 3H), 2.76 (t, *J*=6Hz, 5.6Hz, 2H), 4.55 (t, *J*=5.6Hz, 6Hz, 2H), 6.72 (s, 1H), 7.21 (d, *J*=8Hz, 1H), 7.40 (d, *J*=7.2Hz, 1H), 7.66 (d, *J*=8.4H, 1H). ^13^C-NMR (101MHz, Chloroform-*d,* ppm) δ: 160.6, 147.0, 145.2, 135.7, 129.5, 125.2, 123.3, 121.4, 112.8, 62.8, 58.1, 45.8, 19.0, 18.2. HR-MS, m/z calc. for C_15_H_20_N_2_O (M)^+^ 245.2766 found 246.2153 ([M]+H)^+^

**4,8-dimethyl-2-(2-(piperidin-1-yl)ethoxy)quinoline (10).**

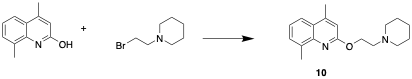

4,8-dimethylquinolin-2-ol (100 mg, 57.7 mmol) and 1-(2-bromoethyl)piperidine (166.3 mg, 86.6 mmol) was used to afford the product follow the same condition of **8** (28 mg, 9.8 mmol, yield=17% ^1^H-NMR (400MHZ, Chloroform-*d,* ppm) δ: 1.33-1.36 (m, 2H), 1.51-1.56 (m, 4H), 2.45 (s, 3H), 2.50 (t, *J*=16Hz, 16Hz, 4H) 2.54 (s, 3H), 2.77 (t, *J*=6Hz, 6Hz, 2H), 2.54 (t, , *J*=6Hz, 6Hz, 2H), 6.61 (s, 1H), 7.12 (t, *J*=5.6Hz, 5.6Hz, 1H), 7.32 (d, , *J*=7.2Hz, 1H), 7.57 (d, , *J*=8Hz, 1H). ^13^C-NMR (101MHz, Chloroform-*d,* ppm) δ:160.4, 147.0, 145.2, 135.7, 129.5, 125.1, 123.3, 121.4, 112.6, 62.6, 57.6, 54.9, 36.5, 25.6, 23.9, 19.0, 18.2. HR-MS, m/z calc. for C_18_H_24_N_2_O (M)^+^ 285.1933 found 2854.9756 ([M]+H)^+^

**4,8-dimethyl-2-(2-(4-methylpiperazin-1-yl)ethoxy)quinoline (11).**

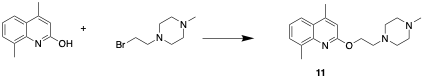

4,8-dimethylquinolin-2-ol (100 mg, 57.7 mmol) and 1-(2-bromoethyl)-4-methylpiperazine (179.4 mg, 86.6 mmol) was used to afford the product follow the same condition of **8** (38 mg, 12.7 mmol, yield=22%). ^1^H-NMR (400MHZ, Chloroform-*d,* ppm) δ: 2.18 (s, 3H), 2.45 (t, *J*=9.2Hz, 6Hz,4H), 2.53 (t, *J*=9.2Hz, 5.6Hz,4H), 2.74 (t, *J*=6Hz, 6Hz,2H), 4.48 (t, *J*=6Hz, 6Hz,2H), 6.61 (s. 1H), 7.13 (t, *J*=8.4Hz, 6.8Hz,1H), 7.31 (d, *J*=5.6Hz, 1H), 7.56 (d, *J*=5.6Hz, 1H). ^13^C-NMR (101MHz, DMSO-*d_6_,* ppm) δ: 167.3, 153.1, 147.3, 137.6, 127.8, 126.7, 126.6, 126.1, 112.9, 63.6, 54.9, 54.9, 53.8, 53.0, 46.1, 17.6, 18.7. HR-MS, m/z calc. for C_18_H_25_N_3_O (M)^+^ 300.2978 found 301.2694([M]+H)^+^.

**4,8-dimethyl-2-(quinolin-8-ylmethoxy) quinoline (12).**

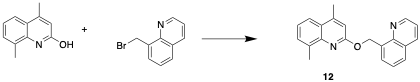

4,8-dimethylquinolin-2-ol (100 mg, 57.7 mmol) and 8-(bromomethyl)quinoline (192.3 mg, 86.6 mmol) was used to afford the product follow the same condition of **8**. (46 mg, 14.6 mmol, yield=25.6%). ^1^H-NMR (400MHZ, Chloroform-*d,* ppm) δ: 2.51 (s, 3H), 2.58 (s, 3H), 6.24 (s, 2H), 6.79 (s, 1H), 7.18 (t, *J*=7.2Hz, 8.8Hz, 1H), 7.33 (d, *J*= 4.4Hz, 1H), 7.51 (t, *J*=6.8Hz, 6.4Hz,1H), 7.43 (t, *J*=7.2Hz, 8Hz,1H), 7.62 (d, *J*=8Hz, 1H), 7.65 (d, *J*=9.2Hz, 1H), 8.04(d, *J*=7.2Hz, 1H), 8.92 (d, *J*= 4.4Hz, 1H). ^13^C-NMR (101MHz, DMSO-*d_6_,* ppm) δ: 167.3, 153.1, 153.2, 148.3, 147.3, 137.6, 130.9, 129.6, 127.9, 127.8, 127.8, 126.7, 126.6, 126.6, 126.5, 126.0, 122.1, 112.9, 69.5, 18.7, 17.6. HR-MS, m/z calc. for C_21_H_17_N_2_O (M)^+^ 315.1439 found 316.1422 ([M]+H)^+^.

**1-(3-(4,8-dimethyl-2-(2-(pyrrolidin-1-yl)ethoxy)quinolin-7-yl)thiophen-2-yl)ethan-1-one (1.1).**

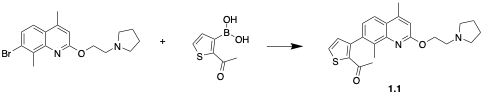

Compound **1** (100 mg, 28.6 mmol), and 2-acetylthiophen-3-yl) boronic acid (72.9 mg, 42.9 mmol) were used to afford the compound **1.1** (25 mg, 6.3 mmol, yield=22%). ^1^H-NMR (400MHZ, DMSO-*d_6_,* ppm) δ:1.86-1.90 (m, 4H), 2.27 (s, 3H), 2.31 (s, 3H), 3.8 (s,3H), 2.85 (t, *J*=8 Hz, 8.4Hz, 2H), 2.97-3.01 (m, 4H), 4.20 (t, *J*=6.4 Hz, 8.4Hz, 2H), 6.91 (s, 1H), 7.40 (d, *J*=6.4 Hz, 1H), 7.51 (d, *J*=8.4 Hz, 1H), 7.65 (d, *J*=6.4 Hz, 1H), 8.32 (d, *J*=8.4 Hz, 1H). ^13^C-NMR (101MHZ, DMSO-*d_6_,* ppm) δ: 191.4, 167.3, 147.3, 147.2, 145.8, 132.7, 129.2, 128.8, 127.6, 127.4, 126.7, 127.4, 126.7, 123.9, 112.9, 62.6, 55.0, 54.7, 54.6, 27.4, 23.4, 23.4, 18.7. HR-MS, m/z calc. for C_23_H_26_N_2_O_2_S(M)^+^ 396.1779 found 396.1812 ([M]+H)^+^

**7-(benzo[*b*]thiophen-3-yl)-4,8-dimethyl-2-(2-(pyrrolidin-1-yl)ethoxy)quinoline (1.2).**

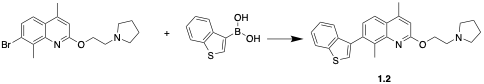

Compound **1** and benzo[*b*]thiophen-3-ylboronic acid (76.4 mg, 42.9 mmol) were used to obtain the compound **1.2** (31 mg, 7.7 mmol, yield=27%). ^1^H-NMR (400MHZ, Chloroform-*d,* ppm) δ: 1.90-1.95 (m,4H), 2.56 (s, 3H), 2.69 (s,3H), 2.95 (t, *J*=6.8Hz, 8Hz, 4H), 3.23 (t, *J*=5.6Hz, 5.2Hz, 2H), 4.80 (t, *J*=5.6Hz, 5.2Hz, 2H), 6.87 (s,1H), 7.34 (t, *J*=6Hz, 6Hz, 1H),7.37 (s, 1H), 7.39 (t, *J*=6Hz, 6Hz, 1H), 7.47 (d, *J*=8.4Hz, 1H), 7.82 (d, *J*=8.4Hz, 1H), 7.96 (d, *J*=7.6Hz, 1H). ^13^C-NMR (101MHz, DMSO-*d_6_,* ppm) δ: 164.3, 153.1, 147.3, 147.1, 138.1, 137.4, 135.0, 130.7, 130.2, 127.8, 126.7, 126.0, 125.4, 125.3, 122.3, 121.7, 112.9, 62.6, 54.9, 54.7, 54.8, 23.4, 23.4, 18.9, 18.7. HR-MS, m/z calc. for C_25_H_26_N_2_OS (M)^+^ 403.1928 found 404.1863 ([M]+H)^+^.

**4,8-dimethyl-7-phenethyl-2-(2-(pyrrolidin-1-yl)ethoxy)quinoline (1.3).**

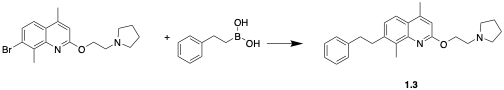

Compound **1** and phenethylboronic acid (64.3 mg, 42.9 mmol) were used to afford the compound **1.3** (42 mg, 11.2 mmol, yield=39.2%). ^1^H-NMR (400MHZ, Chloroform-*d,* ppm) δ: 1.97-2.01 (m, 4H), 2.67 (s, 3H), 2.76 (s, 3H), 2.95-3.02 (m, 4H), 3.10 (t, *J*=4.8Hz, 8Hz, 2H), 3.25 (t, *J*=5.6Hz, 6Hz, 2H), 5.12 (t, *J*=4.8Hz, 4.8Hz, 2H), 5.32 (t, *J*=4.8Hz, 4.8Hz, 2H), 6.82 (s, 1 H) 7.57 (d, *J*=4.2Hz, 1H), 7.68 (t, *J*=4.8Hz, 6Hz, 1H), 7.71 (t, *J*=4.8Hz, 6Hz, 2H), 8.01 (d, *J*=8Hz, 2H), 8.23 (d, *J*=9.6Hz, 1H). ^13^C-NMR (101MHz, DMSO-*d_6_,* ppm) δ: 162.4, 146.3, 145.3, 139.0, 136.5, 129.0, 128.7, 128.7, 128.3, 127.9, 127.8, 127.4, 126.9, 126.7, 112.9, 62.6, 54.0, 54.8, 54.7, 35.8, 35.7, 23.4, 23.3, 18.6. HR-MS, m/z calc. for C_25_H_30_N_2_O (M)^+^ 375.3723 found 376.2720 ([M]+H)^+^.

**7-(furan-3-yl)-4,8-dimethyl-2-(2-(pyrrolidin-1-yl)ethoxy)quinoline (1.4).**

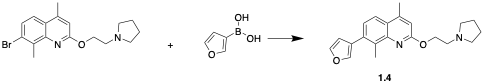

Compound **1** and furan-3-ylboronic acid (48 mg, 42.9 mmol) were used to afford the compound **1.4** (56 mg, 16.6 mmol, yield=58%). ^1^H-NMR (400MHZ, DMSO-*d_6_,* ppm) δ: 1.85-1.89 (m, 4H), 2.25 (s, 3H), 2.45 (s, 3H), 2.85 (t, *J*=8.4Hz, 5.6Hz, 2H), 2.97-3.00 (m, 4H), 4.14 (t, *J*=8Hz, 5.6Hz, 2H) 6.45 (d, *J*=4Hz, 1H), 6.80 (s, 1H), 7.3 (d, *J*=8.4Hz, 1H), 7.69 (d, *J*=4.8Hz, 1H), 8.12(s, 1H), 8.28 (d, *J*=8.4Hz, 1H). ^13^C-NMR (101MHZ, DMSO-*d_6_,* ppm) δ: 167.1, 147.3, 147.2, 143.1, 141.1, 132.7, 127.8, 127.4, 125.7, 126.6, 123.9, 112.9, 109.8, 62.6, 54.9, 54.7, 54.6, 23.4, 23.3, 18.7. HR-MS, m/z calc. for C_21_H_24_N_2_O_2_ (M)^+^ 337.1904 found 338.1927 ([M]+H)^+^.

**7-(furan-2-yl)-4,8-dimethyl-2-(2-(pyrrolidin-1-yl)ethoxy)quinoline (1.5).**

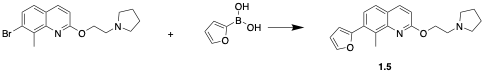

Compound **1** and furan-2-ylboronic acid (48 mg, 42.9 mmol) were used to afford the compound **1.5** (46 mg, 13.7 mmol, yield=47.8). ^1^H-NMR (400MHZ, DMSO-*d_6_,* ppm) δ: 1.86-1.90 (m, 4H), 2.29 (s, 3H), 2.44(s, 3H), 2.85 (t, *J*=8.4Hz, 8.4Hz, 2H), 2.97-3.01 (m, 4H), 4.15 (t, *J*=6Hz, 6.4Hz, 2H), 6.56 (t, *J*=4.0Hz, 4.8Hz, 1H), 6.81 (s, 1H), 6.85 (d, *J*=5.6Hz, 1H), 7.82 (d, *J*=6Hz, 2H), 8.17 (d, *J*=8.4Hz, 1H), 8.40 (d, *J*=5.6Hz, 2H). ^13^C-NMR (101MHZ, DMSO-*d_6_,* ppm) δ: 167.3, 154.1, 147.3, 147.4, 142.3, 128.2, 127.8, 127.4, 126.7, 126.6, 112.9, 111.4, 104.8, 62.6, 55.0, 54.7, 54.7, 23.4, 23.4, 18.7. HR-MS, m/z calc. for C_21_H_24_N_2_O_2_ (M)^+^ 348.2070 found 348.2102 ([M]+H)^+^.

**4,8-dimethyl-7-phenyl-2-(2-(pyrrolidin-1-yl)ethoxy)quinoline (1.6).**

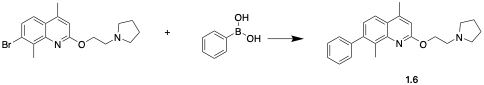

Compound **1** and phenylboronic acid (52.3 mg, 42.9 mmol) were used to afford the title compound (52 mg, 15 mmol, yield=52.5%). ^1^H-NMR (400MHZ, Chloroform-*d,* ppm) δ: 1.88-1.92 (m, 4H), 2.65 (s, 3H), 2.67 (s, 3H), 2.81 (t, *J*=8Hz, 8.4Hz, 4H), 3.10 (t, *J*=4.8Hz, 8Hz, 2H), 4.75 (t, *J*=4.8Hz, 4.8Hz, 2H), 6.84 (s, 1H), 7.34 (d, *J*=4.2Hz, 1H), 7.39 (t, *J*=4.8Hz, 6Hz, 1H), 7.43 (t, *J*=4.8Hz, 6Hz,

2H), 7.48 (d, *J*=8Hz, 2H), 7.79 (d, *J*=9.6Hz, 1H). ^13^C-NMR (101MHz, DMSO-*d_6_,* ppm) δ: 167.3, 153.1, 147.3, 139.5, 137.6, 130.7, 130.1, 128.0, 128.8, 128.7, 128.6, 128.6, 126.7, 126.1, 112.9, 62.7, 55.1, 55.0, 55.0, 23.4, 23.3, 19.1, 18.3. HR-MS, m/z calc. for C_23_H_26_N_2_O (M)^+^ 347.2170 found 347.2988 ([M]+H)^+^.

**4,8-dimethyl-7-(naphthalen-2-yl)-2-(2-(pyrrolidin-1-yl)ethoxy)quinoline (1.7).**

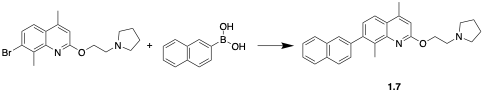

Compound **1** and naphthalen-2-ylboronic acid (73.8 mg, 42.9 mmol) were used to afford the compound **1.7** (70 mg, 17.7 mmol, yield=61.7%). ^1^H-NMR (400MHZ, DMSO-*d_6_,* ppm) δ: 1.88-1.92 (m, 4H), 2.26 (s, 3H), 2.52 (s, 3H), 2.87 (t, *J*=8.4Hz, 8Hz, 2H), 3.01-3.05 (m, 4H), 4.16 (t, *J*=8Hz, 5.6Hz, 2H), 6.65 (s, 1H), 7.55 (s, 1H), 7.57 (s, 1H), 7.70 (d, *J*=1.2Hz,1H), 7.72 (d, *J*=1.2Hz,1H), 7.87 (d, *J*=8.4Hz, 2H), 7.92 (t, *J*=8Hz, 5.6Hz, 2H), 8.31 (d, *J*=5.6Hz, 1H), 8.51 (t, *J*=6Hz, 5.6Hz, 1H). ^13^C-NMR (101MHZ, DMSO-*d_6_,* ppm) δ: 167.1, 147.3, 147.2,140.3, 139.2, 133.7, 133.6, 128.5, 128.4, 127.7, 127.3, 126.7, 126.6, 126.6, 126.5, 126.5, 125.7, 125.6, 124.5, 112.9, 62.6, 55.0, 54.7, 54.7, 23.4, 23.3, 18.7. HR-MS, m/z calc. for C_27_H_28_N_2_O (M)^+^ 397.2852 found 398.1021 ([M]+H)^+^.

**4',8'-dimethyl-2'-(2-(pyrrolidin-1-yl) ethoxy)-3,7'-biquinoline. (1.8)**

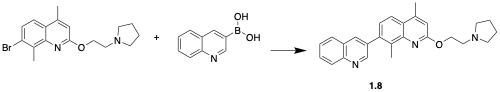

Compound **1** and quinolin-3-ylboronic acid (74.2 mg, 42.9 mmol) were used to afford the compound **1.8** (65 mg, 16.4 mmol, yield=57.2%). ^1^H-NMR (400MHZ, DMSO-*d_6_,* ppm) δ: 1.86-1.90 (m, 4H), 2.23 (s, 3H), 2.52 (s, 3H), 2.87 (t, *J*=6.4Hz, 8Hz, 2H), 2.98-3.02 (m, 4H), 4.19(t, *J*=8Hz, 8.4Hz, 2H), 6.83 (s, 1H), 7.66 (t, *J*=8.4Hz, 8Hz 2H), 7.93 (d, *J*=8Hz,2H), 8.19 (d, *J*=8.4Hz, 1H), 8.33 (d, *J*=8Hz, 1H), 8.7 (s, 1H), 9.4 (d, *J*= 5.6Hz, 1H). ^13^C-NMR (101MHZ, DMSO-*d_6_,* ppm) δ: 167.3, 139.0, 148.1, 147.3, 147.2, 139.2, 133.9, 129.1, 129.0, 128.9, 128.1, 127.4, 127.2, 126.7, 126.6, 126.5, 125.7, 112.9, 62.6, 55.1, 54.7, 54.7, 23.4, 23.4, 18.7. HR-MS, m/z calc. for C_26_H_27_N_3_O (M)^+^ 398.2227 found 399.2258 ([M]+H)^+^.

**4,8-dimethyl-7-(pyridin-4-yl)-2-(2-(pyrrolidin-1-yl)ethoxy)quinoline (1.9)**

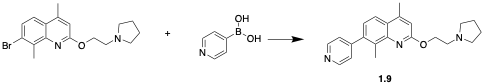

Compound **1** and pyridin-4-ylboronic acid (52.7 mg, 42.9 mmol) were used to afford compound **1.9** (60 mg, 17.2 mmol, yield=60.1%). ^1^H-NMR (400MHZ, DMSO-*d_6_,* ppm) δ: 1.71-1.74 (m, 4H), 2.50-2.51 (m, 4H), 2.58 (s, 3H), 2.65 (s,3H), 2.98 (t, *J*=4.8 Hz, 6.4Hz, 2H), 4.59 (t, *J*=6Hz, 5.6Hz, 2H), 6.96 (s, 1H), 7.11 (d, *J*=4.4Hz, 1H), 7.33 (d, *J*=8.4Hz, 1H), 7.46 (d, *J*=6Hz, 2H), 7.91 (d, *J*=8.4Hz, 1H), 8.68 (d, *J*=5.6Hz, 2H). ^13^C-NMR (101MHZ, DMSO-*d_6_,* ppm) δ: 161.1, 150.1, 149.6, 148.2, 145.1, 139.6, 132.2, 125.2, 124.9, 124.8, 122.3, 113.2, 64.0, 54.4, 54.3, 49.6, 23.5, 18.8, 15.4. HR-MS, m/z calc. for C_22_H_25_N_3_O (M)^+^ 348.2070 found 348.2102 ([M]+H)^+^.

**1-(3-(2-(2-(dimethylamino)ethoxy)-4,8-dimethylquinolin-7-yl)thiophen-2-yl)ethan-1-one (3.1).**

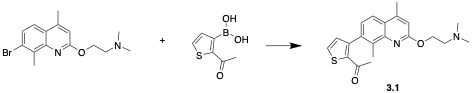

Compounf **3** and 2-acetylthiophen-3-yl)boronic acid (78.7 mg, 42.9 mmol) were used to afford the compound **3.1** (48 mg, 13 mmol, yield=42.1%). ^1^H-NMR (400MHZ, DMSO-*d_6_,* ppm) δ: 2.27 (s,3H), 2.30 (s, 6H), 2.35 (s, 3H), 2.42 (s, 3H), 2.82 (t, *J*=5.6Hz, 6Hz, 2H), 4.14 (t, *J*=5.6Hz, 5.6Hz, 2H), 6.88 (s, 1H), 7.62 (d, *J*=4.8Hz, 1H), 7.52(d, *J*=4.8Hz, 1H), 7.60 (d, *J*=5.6Hz, 1H), 8.34 (d, *J*=8.4Hz, 1H). ^13^C-NMR (101MHZ, DMSO-*d_6_,* ppm) δ: 191.4, 167.3, 147.3, 147.2, 145.8, 132.7, 129.2, 128.8, 127.6, 127.4, 126.7, 126.6, 123.9, 112.9, 62.6, 58.2, 45.6, 45.5, 27.4, 18.7. HR-MS, m/z calc. for C_21_H_24_N_2_O_2_S(M)^+^ 369.1631 found 370.1662 ([M]+H)^+^.

**2-((7-(benzo[*b*]thiophen-3-yl)-4,8-dimethylquinolin-2-yl)oxy)-*N*,*N*-dimethylethan-1-amine (3.2).**

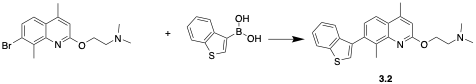

Compound **3** and benzo[*b*]thiophen-3-ylboronic acid (82.5 mg, 46.3 mmol) were used to obtain the title compound (25 mg, 6.6 mmol, yield=21.4%). ^1^H-NMR (400MHZ, DMSO-*d_6_,* ppm) δ:2.23 (s, 3H), 2.30 (s, 6H), 2.36(s, 3H), 2.84 (t, *J*=5.6Hz, 6Hz, 2H), 4.37 (t, *J*=5.6Hz, 6Hz, 2H), 6.83 (s, 1H), 7.29 (t, *J*=6Hz, 6Hz, 2H), 7.34( d, *J*=8.4Hz,1H), 7.89 (d, *J*=8.4Hz, 1H), 8.10 (d, *J*=6.0Hz, 1H), 8.14(d, *J*=8.4Hz, 1H), 8.43 (s, 1H). ^13^C-NMR (101MHZ, DMSO-*d_6_,* ppm) δ:167.0, 147.1, 146.9, 138.1, 137.1, 132.5, 129.9, 127.4, 126.7, 127.6, 125.4, 125.3, 123.7, 122.3, 121.7, 112.9, 62.6, 58.2, 45.6, 45.6, 18.7. HR-MS, m/z calc. for C_23_H_24_N_2_OS (M)^+^ 377.1682 found 378.1708 ([M]+H)^+^.

**2-((4,8-dimethyl-7-phenethylquinolin-2-yl)oxy)-*N*,*N*-dimethylethan-1-amine (3.3).**

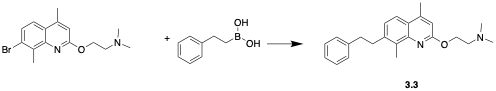

Compound **3** and phenethylboronic acid (69.4 mg, 46.3 mmol) were used to afford the compound **3.3** (28 mg, 8.5mol, yield=25.9%). ^1^H-NMR (400MHZ, CD_3_OD*,* ppm) δ: 2.23 (s, 6H), 2.41 (s, 3H), 2.52 (s, 3H), 2.95 (t, *J*=6Hz, 5.6Hz, 2H), 3.02 (t, *J*=6Hz, 5.6Hz, 2H), 3.15 (t, *J*=6Hz, 5.6Hz, 2H), 4.82 (t, *J*=5.6Hz, 5.6Hz, 2H), 6.56 (s, 1 H), 7.34 (d, *J*=4.2Hz, 1H), 7.53 (t, *J*=4.8Hz, 6Hz, 1H), 7.72 (t, *J*=4.8Hz, 6Hz, 2H), 8.01 (d, *J*=8Hz, 2H), 8.23 (d, *J*=9.6Hz, 1H). ^13^C-NMR (101MHZ, DMSO-*d_6_,* ppm) δ: 167.3, 147.3, 147.2, 143.0, 136.5, 128.9, 128.7, 128.7, 127.5, 128.4, 128.0, 127.4, 126.7, 126.6, 112.9, 62.3, 58.2, 45.6, 45.6, 36.9, 36.8, 18.7. HR-MS, m/z calc. for C_19_H_22_N_2_O_2_ (M)^+^ 311.2301 found 312.1778 ([M]+H)^+^.

**2-((7-(furan-3-yl)-4,8-dimethylquinolin-2-yl)oxy)-*N*,*N*-dimethylethan-1-amine (3.4).**

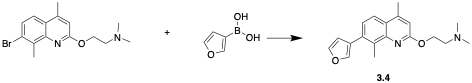

Compound **3** and furan-3-ylboronic acid (51.8 mg, 46.3 mmol) were used to afford the compound **3.4** (42 mg, 13.5 mmol, yield=43.7%). ^1^H-NMR (400MHZ, DMSO-*d_6_,* ppm) δ: 2.29 (s, 6H), 2.62, (s, 3H), 2.72 (s, 3H), 2.77 (t, *J*=5.6Hz, 5.6Hz, 2H), 4.57 (t, *J*=6Hz, 6.4Hz, 2H), 6.88 (s, 1H), 7.45 (d, *J*=8.4Hz, 1H), 7.82 (d, *J*=2Hz, 1H), 7.83 (s, 1H), 7.99 (d, *J*=2Hz, 1H),8.14 (d, *J*=2Hz, 1H). ^13^C-NMR (101MHZ, DMSO-*d_6_,* ppm) δ:161.3, 153.1, 147.3, 143.1, 141.1, 135.0, 130.7, 127.9, 127.8, 126.7, 126.0, 112.9, 109.8, 62.6, 58.2, 45.6, 45.7, 18.9, 18.7. HR-MS, m/z calc. for C_19_H_22_N_2_O_2_ (M)^+^ 311.1749 found 312.1781 ([M]+H)^+^.

**2-((7-(furan-2-yl)-4,8-dimethylquinolin-2-yl)oxy)-*N*,*N*-dimethylethan-1-amine (3.5).**

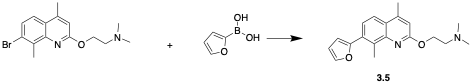

Compound **3** and furan-2-ylboronic acid (51.8 mg, 46.3 mmol) were used to afford the compound **3.5** (30 mg, 9.7 mmol, yield=31.4%).^1^H-NMR (400MHZ, DMSO-*d_6_,* ppm) δ: 2.28 (s, 3H), 2.31 (s, 6H), 2.42(s, 3H), 2.80 (t, *J*=8.2Hz, 6.4Hz, 2H), 4.15 (t, *J*=6Hz, 6.4Hz, 2H), 6.56 (t, *J*=4.0Hz, 4.8Hz, 1H),6.85 (s, 1H), 6.89 (d, *J*=5.6Hz, 1H), 7.84 (d, *J*=6Hz, 2H), 8.20 (d, *J*=8.4Hz, 1H), 8.41(d, *J*=5.6Hz, 2H). ^13^C-NMR (101MHZ, DMSO-*d_6_,* ppm) δ: 161.3, 154.1, 147.3, 147.4, 142.3, 128.2, 127.8, 127.4, 126.7, 126.6, 112.9, 111.4, 104.8, 62.6, 58.2, 45.6, 45.6, 18.7. HR-MS, m/z calc. for C_19_H_22_N_2_O_2_ (M)^+^ 311.2703 found 311.8955 ([M]+H)^+^.

**2-((4,8-dimethyl-7-phenylquinolin-2-yl)oxy)-*N*,*N*-dimethylethan-1-amine (3.6).**

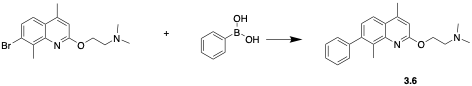

Compound **3** and phenylboronic acid (56.4mg, 42.9mmol) were used to afford the compound **3.6** (38 mg, 11.8 mmol, yield=38.2%). ^1^H-NMR (400MHZ, CD_3_OD*,* ppm) δ: 2.44 (s, 6H), 2.60 (s, 3H), 2.65 (s, 3H), 2.94 (t, *J*=6Hz, 5.6Hz, 2H), 4.66 (t, *J*=5.6Hz, 5.6Hz, 2H), 6.83 (s, 1H), 7.29 (d, *J*=4.2Hz, 1H), 7.37 (t, *J*=4.8Hz, 6Hz, 1H), 7.39 (t, *J*=4.8Hz, 6Hz, 2H), 7.45 (d, *J*=8Hz, 2H), 7.82 (d, *J*=8.4Hz, 1H). ^13^C-NMR (101MHZ, DMSO-*d_6_,* ppm) δ: 167.3, 153.1, 147.3, 139.5, 137.6, 130.7, 130.0, 128.9, 128.9, 128.8, 128.6, 128.5, 126.7, 112.9, 62.6, 58.2, 45.6, 45.5, 18.9, 18.7. HR-MS, m/z calc. for C_19_H_22_N_2_O_2_ (M)^+^ 311.1955 found 311.8577 ([M]+H)^+^.

**2-((4,8-dimethyl-7-(naphthalen-2-yl)quinolin-2-yl)oxy)-*N*,*N*-dimethylethan-1-amine (3.7).**

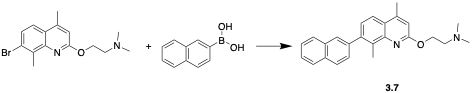

Compound **3** and naphthalen-2-ylboronic acid (79.6 mg, 46.3 mmol) were used to afford the compound **3.7** (56 mg, 15.1 mmol, yield=48.9%). ^1^H-NMR (400MHZ, DMSO-*d_6_,* ppm) δ: 2.24 (s, 6H), 2.3 (s, 3H), 2.59 (s, 3H), 2.83 (t, *J*=6Hz, 5.6Hz, 2H), 4.12 (t, *J*=5.6Hz, 6Hz, 2H), 6.88 (s, 1H), 7.56 (d, *J*=8.4Hz, 6Hz , 2H), 7.71 (d, *J*=8.8Hz, 1H), 7.69 (d, *J*=8Hz, 1H), 7.78 (d, *J*=8.4Hz, 1H), 7.92 (t, *J*=5.6Hz, 6Hz, 2H), 8.30 (d, *J*=8.8Hz, 1H), 8.5(s, 1H). ^13^C-NMR (101MHZ, DMSO-*d_6_,* ppm) δ:163.3, 147.3, 147.2, 140.2, 139.2, 133.7, 133.6, 128.5, 127.4, 127.7, 127.4, 126.7, 126.6, 126.5, 126.5, 126.7, 126.6, 124.5, 112.9, 62.6, 58.2, 45.6, 45.6, 18.7. HR-MS, m/z calc. for C_25_H_26_N_2_O (M)^+^ 371.2733 found 372.1021 ([M]+H)^+^.

**2-((4',8'-dimethyl-[3,7'-biquinolin]-2'-yl) oxy)-*N*,*N*-dimethylethan-1-amine (3.8).**

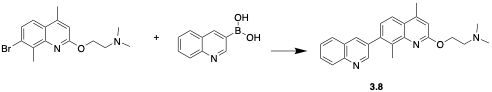

The title compounds were obtained with **3** and quinolin-3-ylboronic acid (80.1mg, 46.3mmol) followed by the general protocol. (53mg, 14.3mmol, yield=46.3%). ^1^H-NMR (400MHZ, DMSO-*d_6_,* ppm) δ:2.23 (s, 6H), 2.58 (s, 3H), 2.59 (s, 3H), 2.70 (t, *J*=5.6Hz, 6Hz, 2H), 4.53 (t, *J*=5.6Hz, 6Hz, 2H), 6.87 (s, 1H), 7.39 (d, *J*=8.4Hz, 1H), 7.63 (t, *J*=7.2Hz, 7.6Hz, 1H), 7.78 (t, *J*=6.8Hz, 8Hz, 1H), 7.84 (d, *J*=8.8Hz, 1H), 8.01 (d, *J*=8Hz, 1H), 8.09 (d, *J*=8.4Hz, 1H), 8.37 (s, 1H), 8.96 (d, *J*=2Hz, 1H). ^13^C-NMR (101MHZ, DMSO-*d_6_,* ppm) δ:161.0, 151.7, 147.9, 147.0, 145.2, 138.5, 135.9, 134.9, 132.8, 130.1, 129.2, 128.7, 127.8, 127.4, 126.0, 124.6, 122.1, 113.0, 63.2, 57.8, 45.8, 18.7, 15.5. HR-MS, m/z calc. for C_24_H_25_N_3_O (M)^+^ 371.2 found 372.2 ([M]+H)^+^.

**2-((4,8-dimethyl-7-(pyridin-4-yl)quinolin-2-yl)oxy)-*N*,*N*-dimethylethan-1-amine (3.9).**

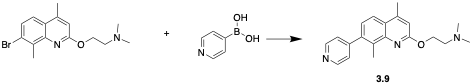

Compound **3** and pyridin-4-ylboronic acid (56.9 mg, 46.3 mmol) were used to afford the product **3.9** (54 mg, 16.8 mmol, yield=54.3%). ^1^H-NMR (400MHZ, DMSO-*d_6_,* ppm) δ: 2.23 (s, 3H), 2.29 (s, 6H), 2.45(s, 3H), 2.82 (t, *J*=6.4Hz, 4Hz, 2H), 4.18 (t, *J*=6Hz, 6.4Hz, 2H), 6.92 (s, 1H), 7.24 (d, *J*=8.4Hz, 1H), 7.93 (d, *J*=6Hz, 2H), 8.36 (d, *J*=8.4Hz, 1H),8.44 (d, *J*=5.6Hz, 2H). ^13^C-NMR (101MHZ, DMSO-*d_6_,* ppm) δ: 167.3, 150.0, 149.8, 147.3, 147.2, 147.0, 139.2, 127.4, 126.7, 126.6, 125.7, 123.1, 123.1, 112.9, 62.6, 58.2, 45.6, 46.6, 18.7. HR-MS, m/z calc. for C_20_ H_23_N_2_OS (M)^+^ 372.2070 found 373.2102 ([M]+H)^+^.

**7-bromo-2-(2-(pyrrolidin-1-yl)ethoxy)quinoline (1.10).**

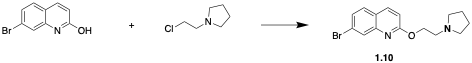

7-bromoquinolin-2-ol (200 mg, 89.2 mmol), 1-(2-chloroethyl)pyrrolidine (99.3 mg, 107 mmol) and K_2_CO_3_ (246.6 mg, 178.4 mmol) were mixed in 15mL DMF in a microwave-proof glass tube. The mixture was heated for 2 hrs at 170^o^C in a microwave synthesis. Upon cooling, the reaction was extracted with EtOAc and H_2_O. MgSO_4_ was used to dry the separated organic layer. The title compound was purified by flash column chromatography. ^1^H-NMR (400MHZ, CD_3_OD*,* ppm) δ: 1.96-1.87 (m, 4H), 2.75 (t, *J*= 6.8Hz, 6.4Hz, 3H), 3.01 (t, *J*= 5.6Hz, 6Hz, 2H), 4.52 (t, *J*= 5.6Hz, 6Hz, 2H), 7.09 (d, *J*= 4.8Hz, 1H), 7.51 (d, *J*= 8Hz, 1H), 7.89 (d, *J*=8Hz, 1H), 8.00 (d, *J*= 4Hz, 1H), 8.95 (s, 1H). ^13^C-NMR (101MHZ, DMSO-*d_6_,* ppm) δ: 163.5, 147.3, 130.2, 128.3, 127.6, 126.9, 126.7, 120.0, 118.2, 62.6, 55.0, 54.7, 54.7, 23.4, 23.4. HR-MS, m/z calc. for C_15_H_17_BrN_2_O (M)^+^ 321.0723 found 321.5149 ([M]+H)^+^.

**7-phenyl-2-(2-(pyrrolidin-1-yl)ethoxy)quinoline (1.11).**

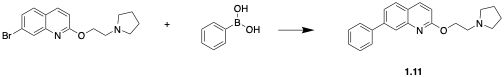

**1.10** (100 mg, 31.1 mmol), phenylboronic acid (60 mg, 37.4 mmol), CsCO_3_(121.6 mg, 37.32 mmol), and Pd(PPh_3_)_4_ (10%mol, 3.11 mmol) was mixed in the round bottom flask, followed by addition of 5mL mixture of toluene and MeOH (4:1). The reaction was heated to reflux for overnight. Upon cooling, EtOAc and H_2_O were used for extraction. The organic layer was dried through MgSO_4_ and evaporated. The final compound was obtained by purification using flash column chromatography (68 mg, 21.4 mmol, yield=68.8%).^1^H-NMR (400MHZ, CD_3_OD*,* ppm) δ: 1.81-1.85 (m, 4H), 2.85 (t, *J=* 8.4HZ, 8.4Hz, 2H), 3.01-3.05 (m, 4H), 4.55 (t, *J=* 8.4HZ, 8.4Hz, 2H), 6.78 (d, *J=* 8.4Hz, 1H), 7.44 (t, *J*= 1.2Hz, 2.4Hz, 1H), 7.50 (t, *J*= 4.8Hz, 2.4Hz, 2H), 7.71 (d, *J*= 6.4Hz, 1H), 7.91 (d, *J*= 6.4Hz, 2H), 8.07 (d, *J*= 8.4Hz, 1H), 8.32 (d, *J*= 5.6Hz, 1H),8.66 (d, *J*= 8.4Hz, 1H). ^13^C-NMR (101MHZ, DMSO-*d_6_,* ppm) δ: 167.3, 147.3, 147.2, 139.2, 139.1, 129.0, 128.9, 128.9, 127.3, 127.1, 127.0, 126.7, 126.6, 125.7, 112.9, 62.6, 55.0, 54.7, 54.7, 23.4, 23.4, 18.7. HR-MS, m/z calc. for C_21_H_22_BN_2_O (M)^+^ 319.1744 found 319.3558 ([M]+H)^+^.

***N*,*N*-dimethyl-2-((7-phenylquinolin-2-yl)oxy)ethan-1-amine (1.12).**

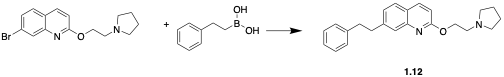

**1.10** (100mg, 31.1mmol), phenethylboronic acid (56 mg, 37.3 mmol), CsCO_3_(121.6 mg, 37.32 mmol), and Pd(PPh_3_)_4_ (10%mol, 3.11mmol) was mixed in the round bottom flask, followed by addition of 5mL mixture of toluene and MeOH (4:1). The reaction was heated to reflux for overnight. Upon cooling, EtOAc and H_2_O were used for extraction. The organic layer was dried through MgSO_4_ and evaporated. The final compound **1.12** was obtained by purification using flash column chromatography (61 mg, 17.6 mmol, yield=56.6%).^1^H NMR (400 MHz, CD_3_OD-*d4,* ppm) δ: 1.97-2.01 (m, 4H), 2.95-3.02 (m, 4H), 3.10 (t, *J*=4.8Hz, 8Hz, 2H), 3.25 (t, *J*=5.6Hz, 6Hz, 2H), 3.82 (t, *J*=4.8Hz, 4.8Hz, 2H), 5.32 (t, *J*=4.8Hz, 4.8Hz, 2H), 6.78 (d, *J=* 8.4Hz, 1H), 7.44 (t, *J*= 1.2Hz, 2.4Hz, 1H), 7.50 (t, *J*= 4.8Hz, 2.4Hz, 2H), 7.71 (d, *J*= 6.4Hz, 1H), 7.91 (d, *J*= 6.4Hz, 2H), 8.07 (d, *J*= 8.4Hz, 1H), 8.32 (d, *J*= 5.6Hz, 1H),8.66 (d, *J*= 8.4Hz, 1H). ^13^C-NMR (101MHZ, DMSO-*d_6_,* ppm) δ: 167.1, 147.1, 146.9, 143.0, 136.5, 128.9, 128.7, 128.7, 128.5, 128.5, 128.0, 127.1, 126.7, 126.6, 112.9, 62.5, 55.0, 54.6, 54.6, 36.9, 36.8, 23.4, 23.4, 18.7. HR-MS, m/z calc. for C_23_H_26_N_2_O (M)^+^ 347.2423 found 348.2337 ([M]+H)^+^.

**2-((7-bromoquinolin-2-yl)oxy)-*N*,*N*-dimethylethan-1-amine (3.10)**

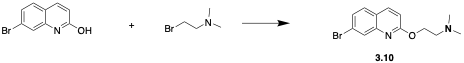

7-bromoquinolin-2-ol (200 mg, 89.2 mmol), 2-bromo-*N*,*N*-dimethylethan-1-amine (203.5 mg, 133.8 mmol)and K_2_CO_3_ (246.6 mg, 178.4 mmol) were mixed in 15 mL DMF in a microwave-proof glass tube. The mixture was heated for 2 hrs at 170^o^C in a microwave synthesis. Upon cooling, the reaction was extracted with EtOAc and H_2_O. MgSO_4_ was used to dry the separated organic layer. The title compound was purified by flash column chromatography. (102 mg, 34.5 mmol, yield=38.7%). ^1^H-NMR (400MHZ, CD_3_OD*,* ppm) δ: 2.41 (S, 6H), 2.88 (t, *J*= 6.4Hz, 6.4Hz, 2H), 4.50 (t, *J*= 6.4Hz, 6.4Hz, 2H), 7.09 (d, *J*= 4.8Hz, 1H), 7.51 (d, *J*= 12.8Hz, 1H), 7.89 (d, *J*=4.8Hz, 1H), 8.01 (d, *J*= 1.6Hz, 1H), 8.96 (s, 1H). ^13^C-NMR (101MHZ, DMSO-*d_6_,* ppm) δ: 163.5, 147.3, 130.2, 128.3, 127.6, 126.9, 126.7, 120.0, 118.2, 62.6, 58.2, 45.6, 45.7. HR-MS, m/z calc. for C_13_H_15_BrN_2_O (M)^+^ 294.9772 found 295.2791 ([M]+H)^+^.

***N*,*N*-dimethyl-2-((7-phenylquinolin-2-yl)oxy)ethan-1-amine. (3.11)**

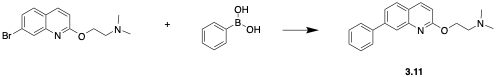

**3.10** (100 mg, 33.9 mmol), phenylboronic acid (65.3 mg, 40.7 mmol), CsCO_3_ (165.7 mg, 50.85 mmol), and Pd(PPh_3_)_4_ (10%mol, 3.39 mmol) was mixed in the round bottom flask, followed by addition of 5 mL mixture of toluene and MeOH (4:1). The reaction was heated to reflux for overnight. Upon cooling, EtOAc and H_2_O were used for extraction. The organic layer was dried through MgSO_4_ and evaporated. The compound **3.11** was obtained by purification using flash column chromatography (49 mg, 16.8 mmol, yield=49.6%).^1^H-NMR (400MHZ, CD_3_OD*,* ppm) δ: 2.45 (s, 6H), 2.93 (t, *J*= 6.4Hz, 65.6Hz, 2H), 4.50 (t, *J*= 6.4Hz, 6.4Hz, 2H), 7.20 (d, *J*= 6.4Hz, 1H), 7.44 (t, *J*= 1.2Hz, 2.4Hz, 1H), 7.49 (t, *J*= 4.8Hz, 2.4Hz, 2H), 7.72 (d, *J*= 6.4Hz, 1H), 7.88 (d, *J*= 6.4Hz, 2H), 8.06 (d, *J*= 8.4Hz, 1H), 8.36 (d, *J*= 5.6Hz, 1H),8.66 (d, *J*= 8.4Hz, 1H). ^13^C-NMR (101MHZ, DMSO-*d_6_,* ppm) δ: 167.3, 147.3, 127.2, 139.2, 129.1, 129.0, 128.9, 128.9, 127.4, 127.1, 126.7, 126.6, 125.7, 112.9, 62.6, 58.2, 45.6, 45.6, 18.7. HR-MS, m/z calc. for C_19_H_20_N_2_O (M)^+^ 293.2133 found 294.1002 ([M]+H)^+^.

***N*,*N*-dimethyl-2-((7-phenethylquinolin-2-yl)oxy)ethan-1-amine (3.12).**

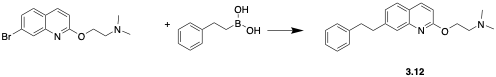

**3.10** (100 mg, 33.9 mmol), phenethylboronic acid (61.1 mg, 40.7 mmol), CsCO_3_(165.7 mg, 50.85 mmol), and Pd(PPh_3_)_4_ (10%mol, 3.39 mmol) was mixed in the round bottom flask, followed by addition of 5mLmixture of toluene and MeOH (4:1). The reaction was heated to reflux for overnight. Upon cooling, EtOAc and water were used for extraction. The organic layer was dried through MgSO_4_ and evaporated. The final compound was purified using flash column chromatography (47 mg, 14.7 mmol, yield=43.4%).1H NMR (400 MHz, CD_3_OD, ppm) δ: 2.43 (s, 6 H) 3.10 (t, *J*=4.8Hz, 8Hz, 2H), 3.25 (t, *J*=5.6Hz, 6Hz, 2H), 3.82 (t, *J*=4.8Hz, 4.8Hz, 2H), 5.32 (t, *J*=4.8Hz, 4.8Hz, 2H), 6.78 (d, *J=* 8.4Hz, 1H), 7.44 (t, *J*= 1.2Hz, 2.4Hz, 1H), 7.50 (t, *J*= 4.8Hz, 2.4Hz, 2H), 7.71 (d, *J*= 6.4Hz, 1H), 7.91 (d, *J*= 6.4Hz, 2H), 8.07 (d, *J*= 8.4Hz, 1H), 8.32 (d, *J*= 5.6Hz, 1H),8.66 (d, *J*= 8.4Hz, 1H). ^13^C-NMR (101 MHZ, DMSO-*d_6_,* ppm) δ: 166.2, 147.1, 146.9, 143.0, 136.5, 128.9, 128.7, 128.5, 128.5, 128.0, 127.3, 126.7, 126.6, 112.9, 62.6, 58.2, 45.6, 45.6, 36.9, 36.8, 18.7. HR-MS, m/z calc. for C_21_H_24_N_2_O (M)^+^ 321.1776 found 322.1135 ([M]+H)

**^1^H and ^13^C NMR spectra of Quinoline based EPI Compounds**

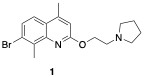

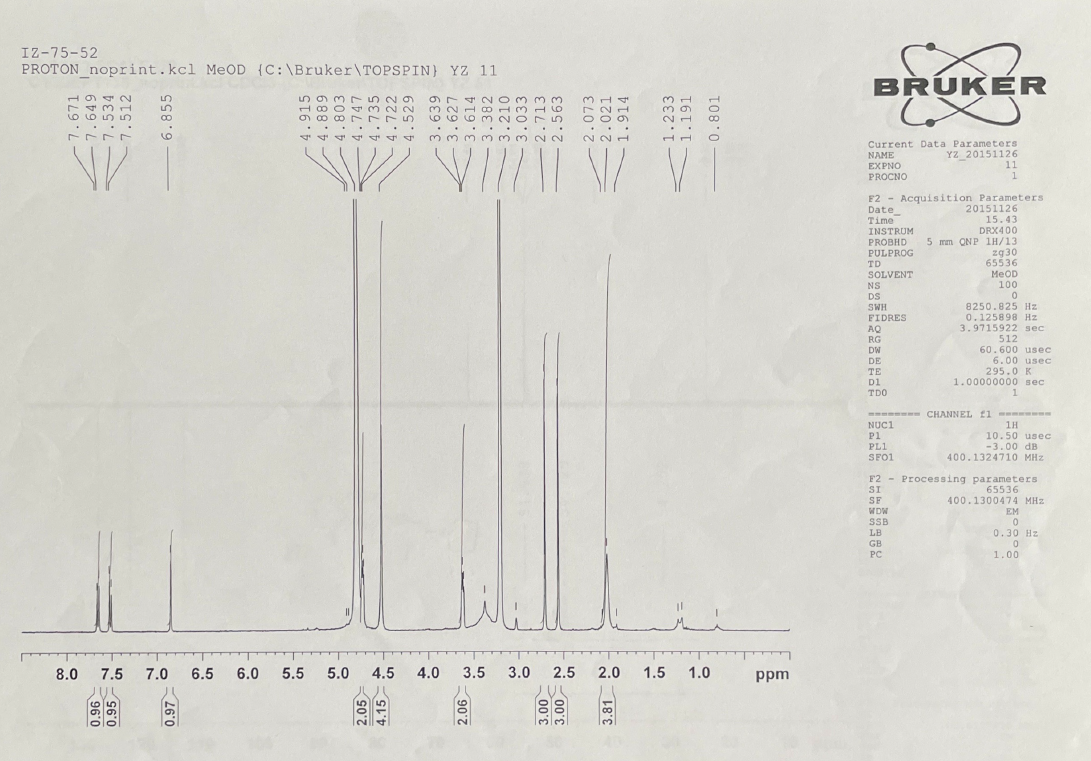

**^1^H-NMR of compound 1**
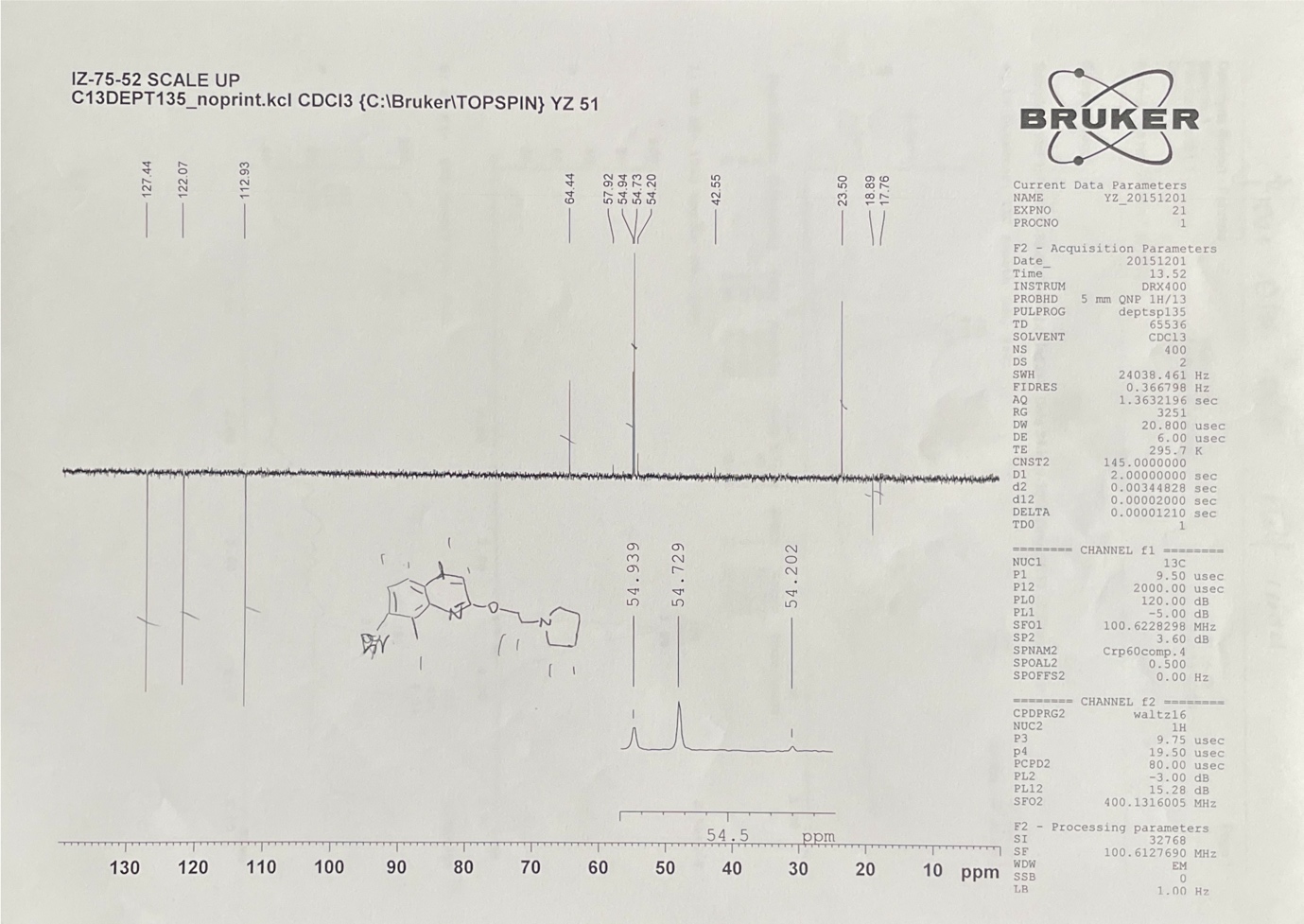

**^13^C-NMR of compound 1**

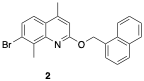

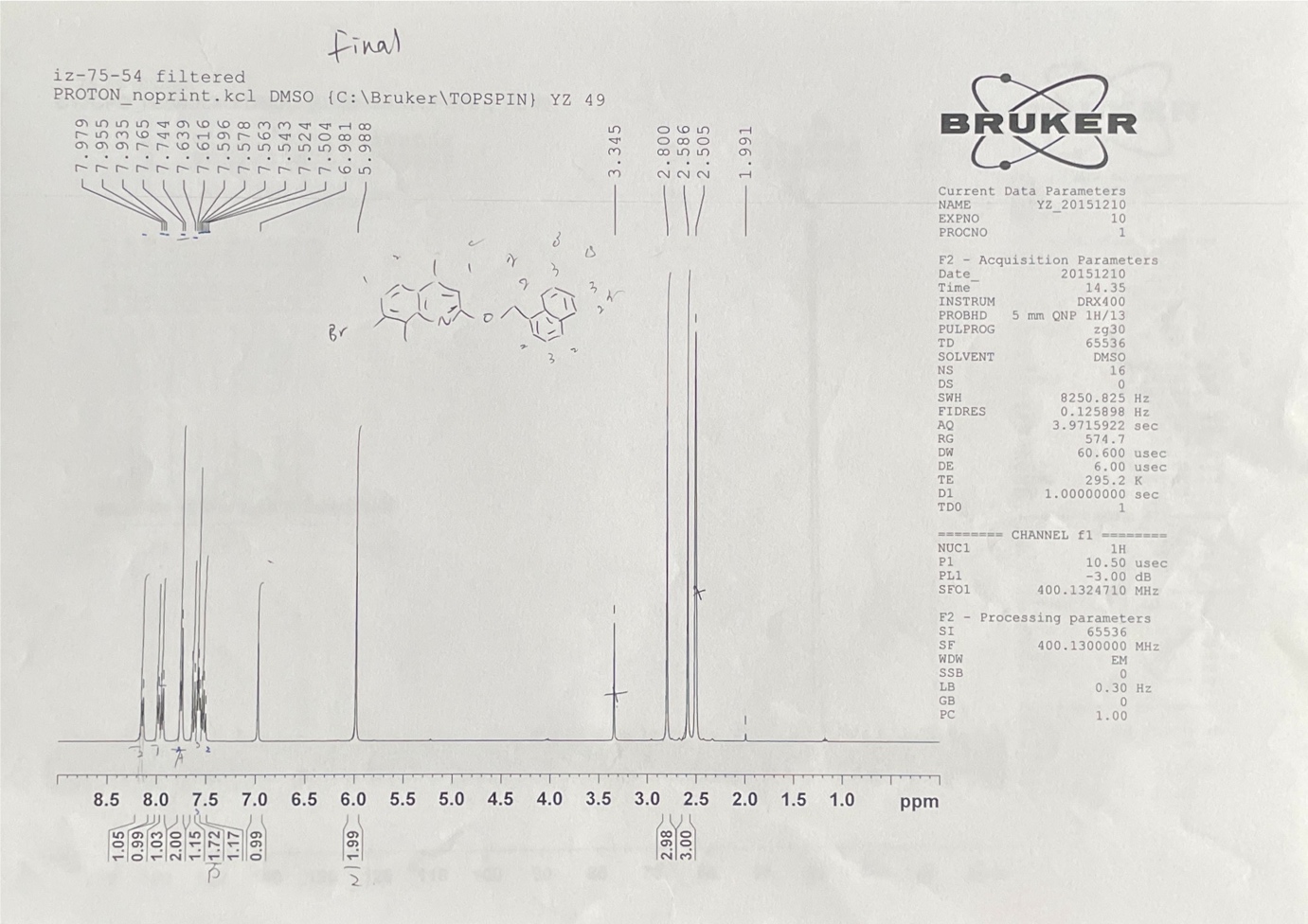

**^1^H-NMR of compound 2**

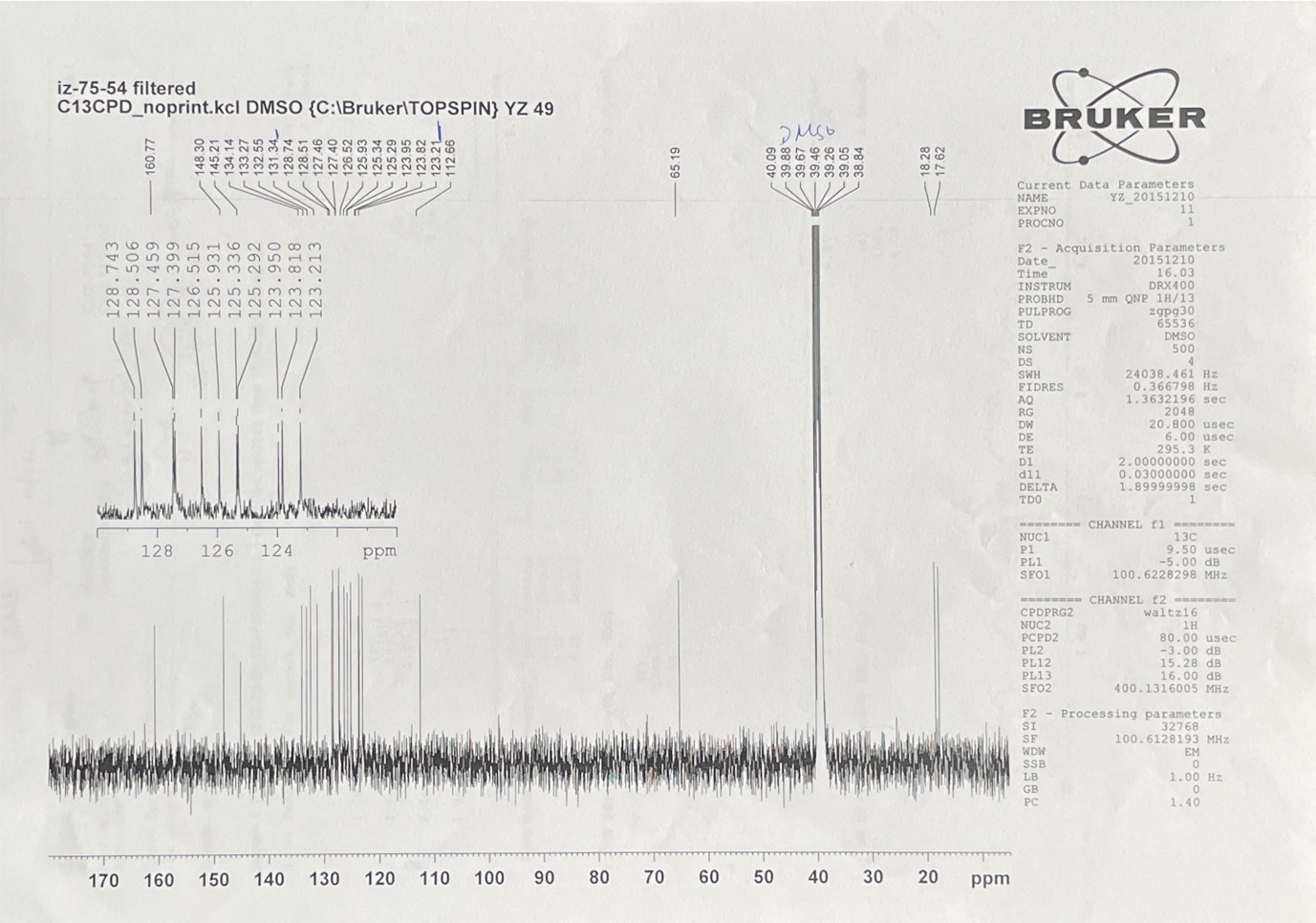

**^13^C-NMR of compound 2**

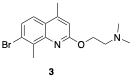

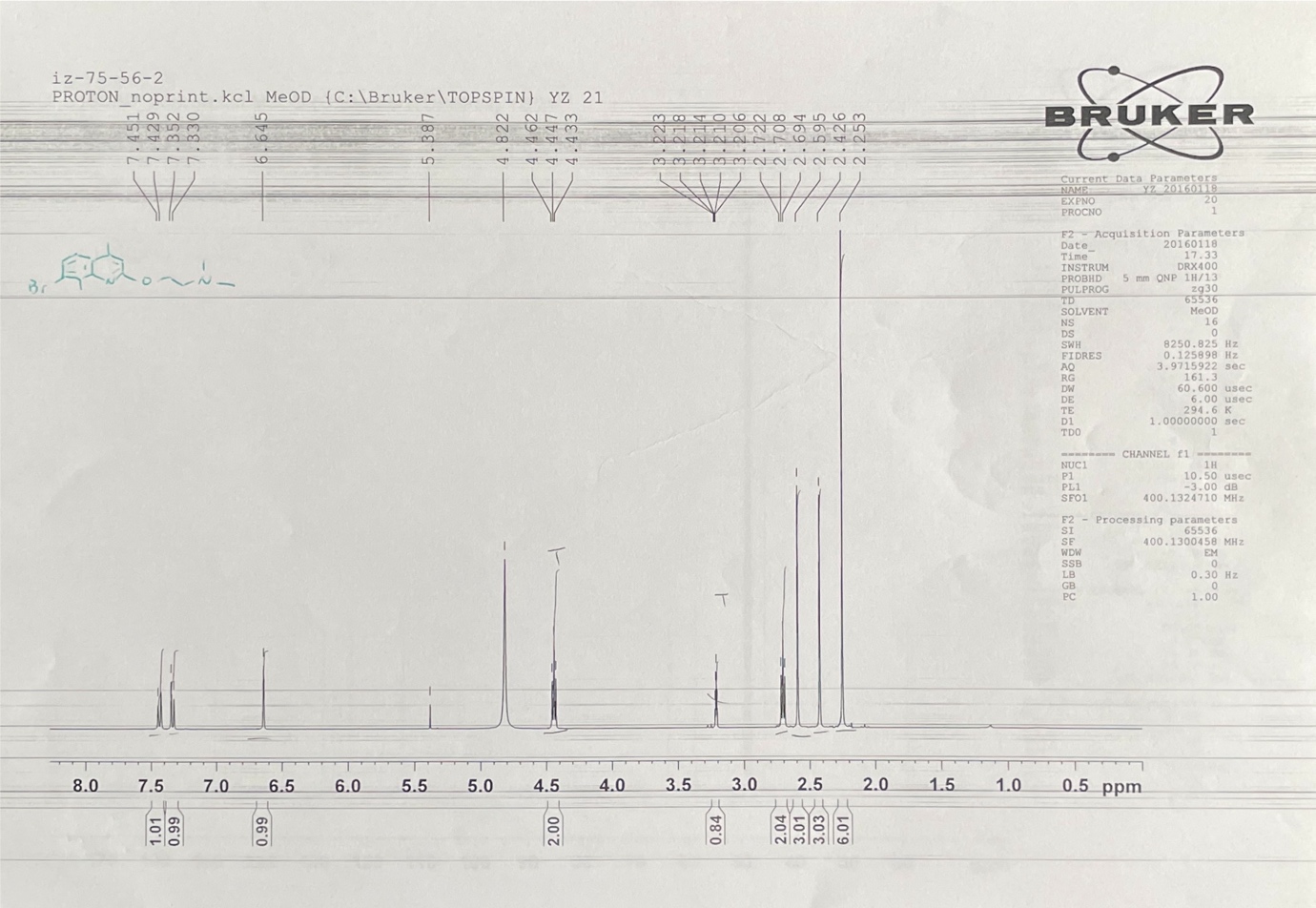

**^1^H-NMR of compound 3**

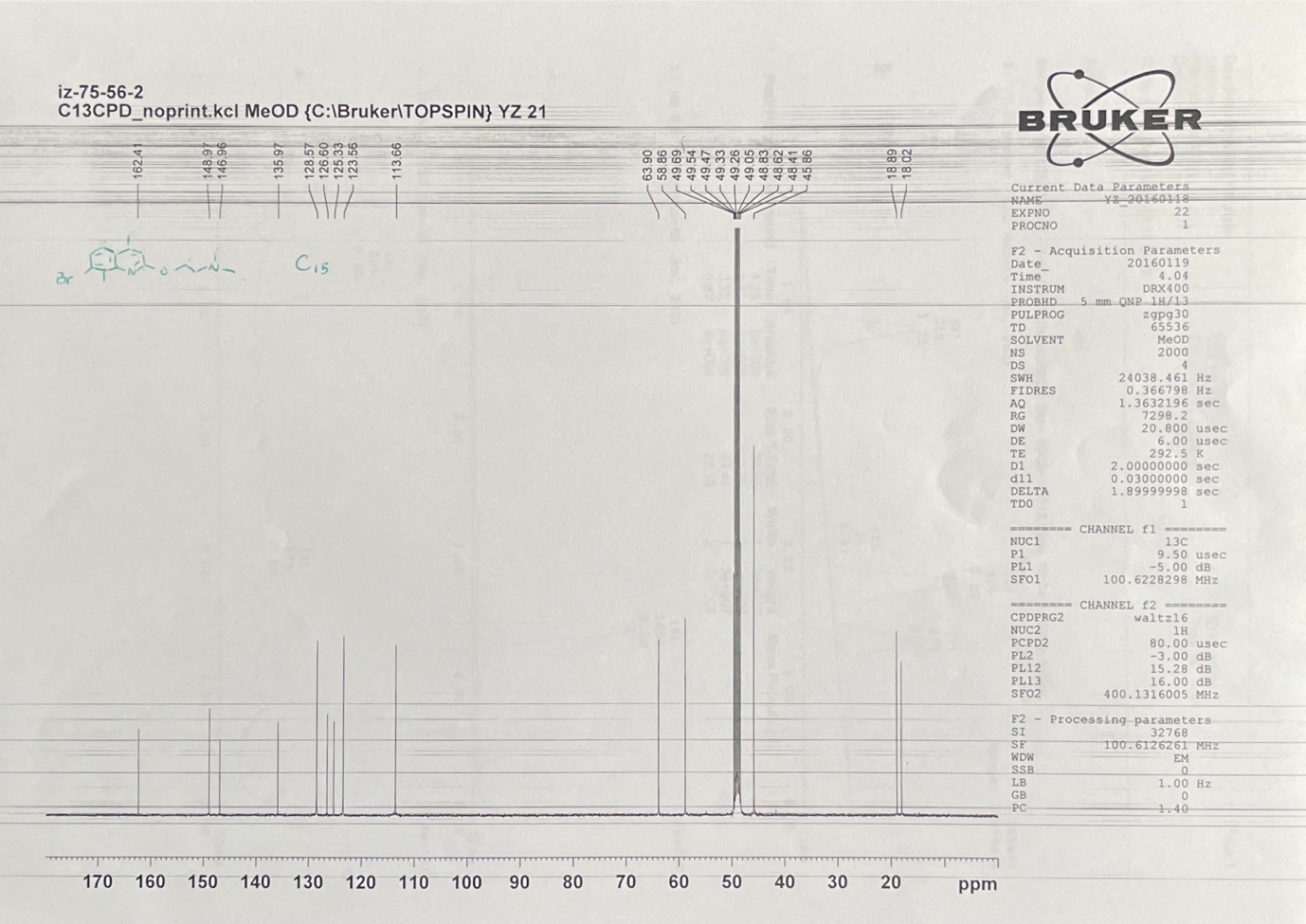

**^13^C-NMR of compound 3**

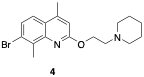

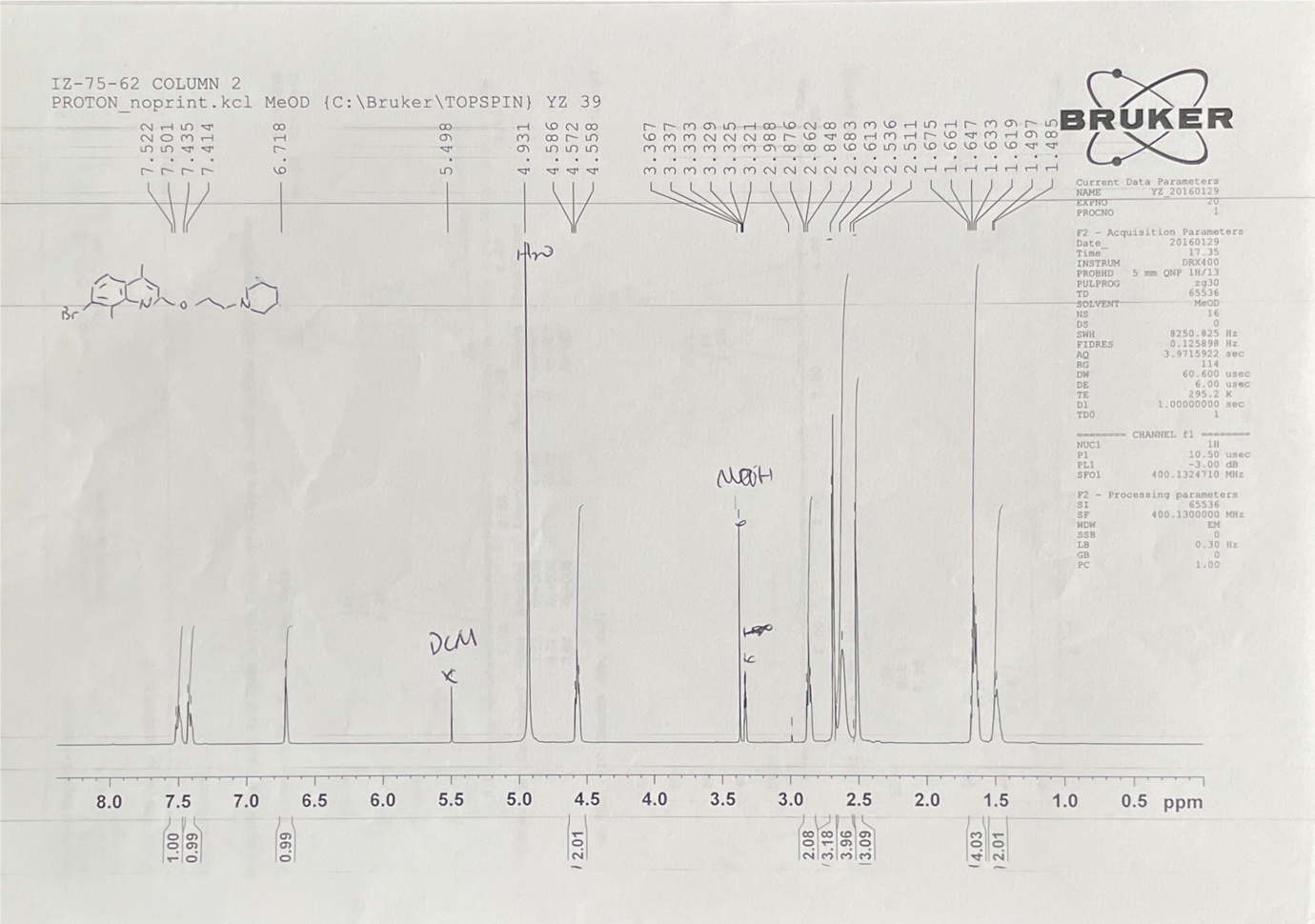

**^1^H-NMR of compound 4**

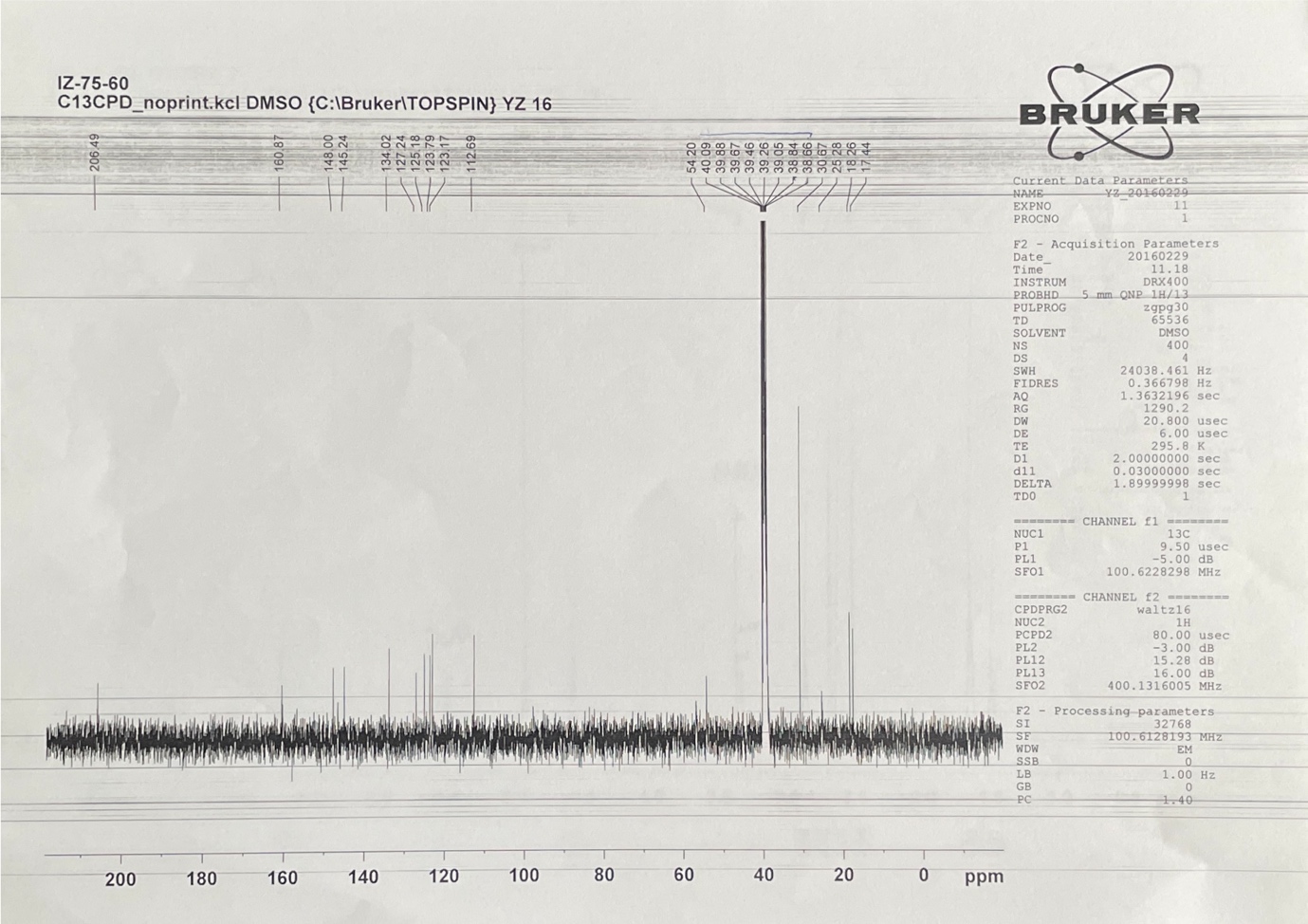

**^13^C-NMR of compound 4**

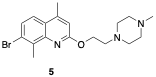

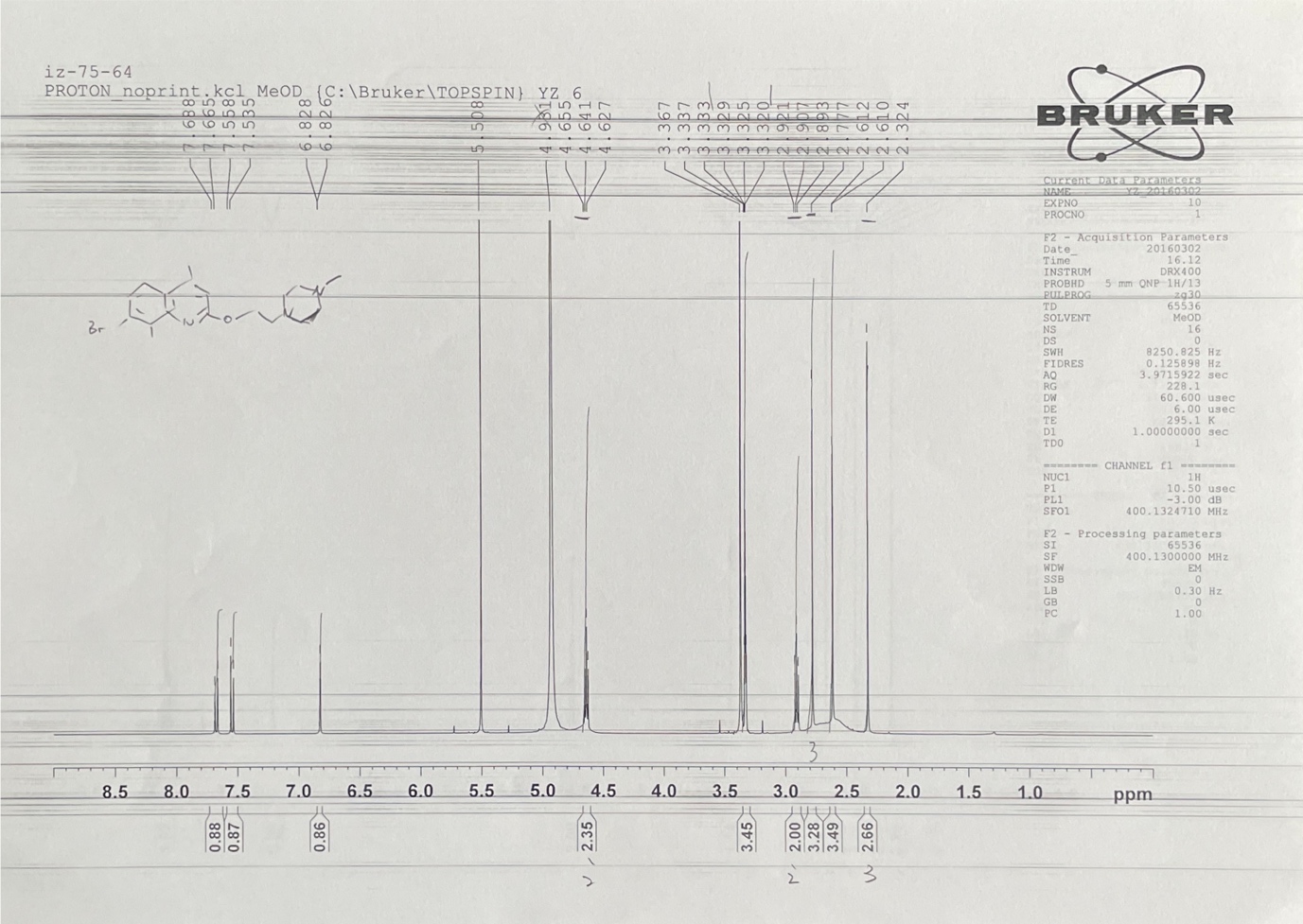

**^1^H-NMR of compound 5**

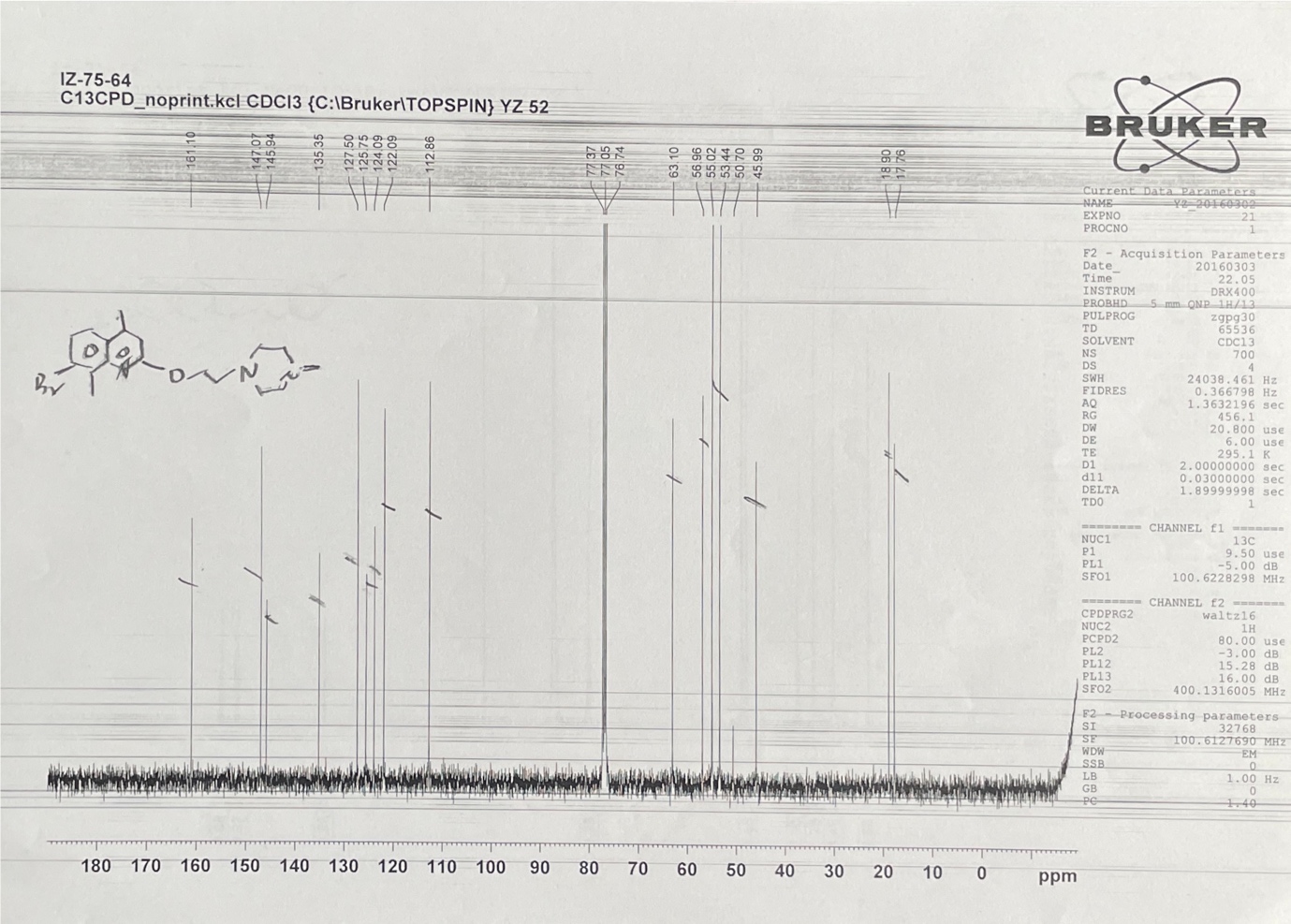

**^13^C-NMR of compound 5**

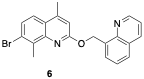

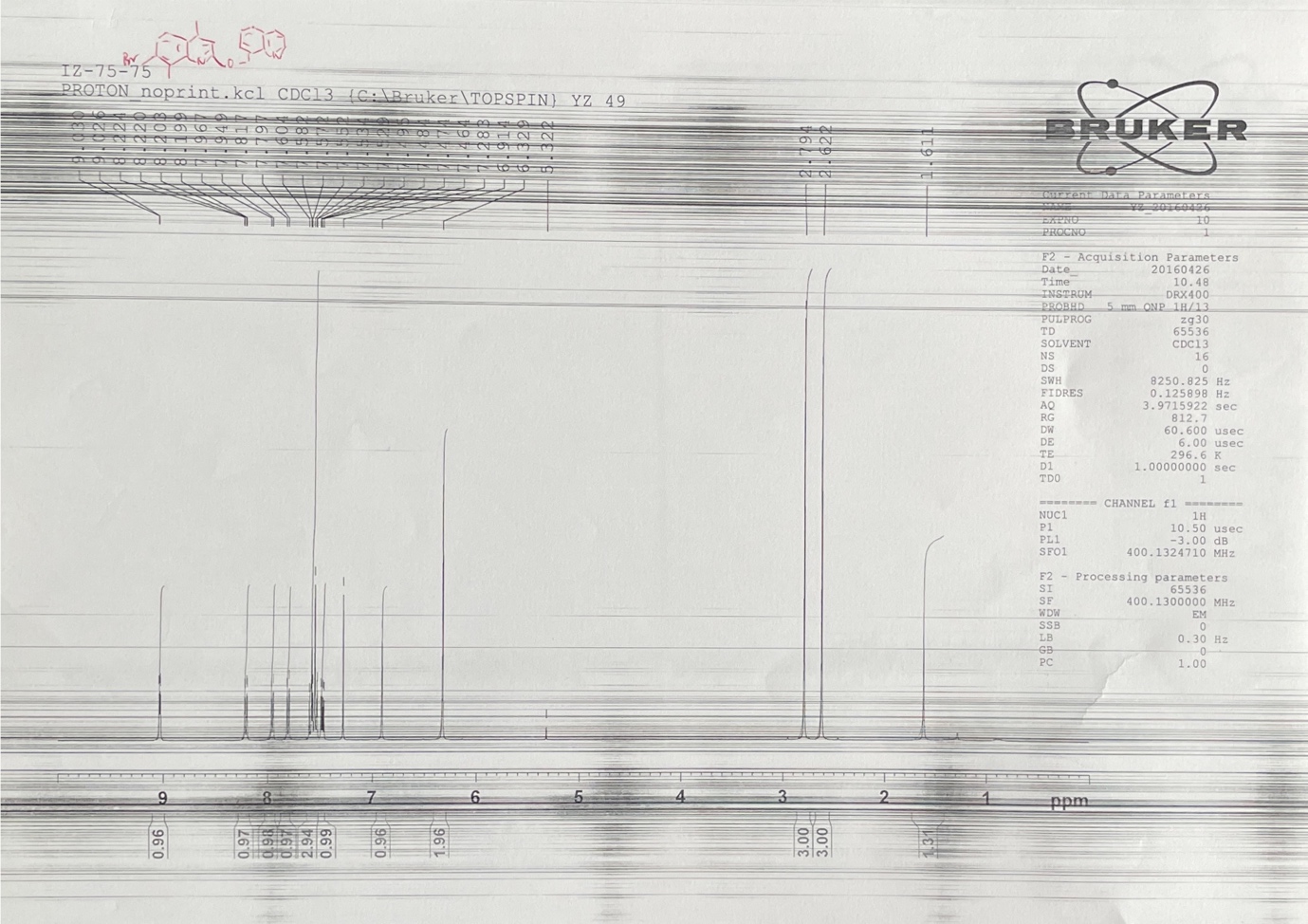

**^1^H-NMR of compound 6**

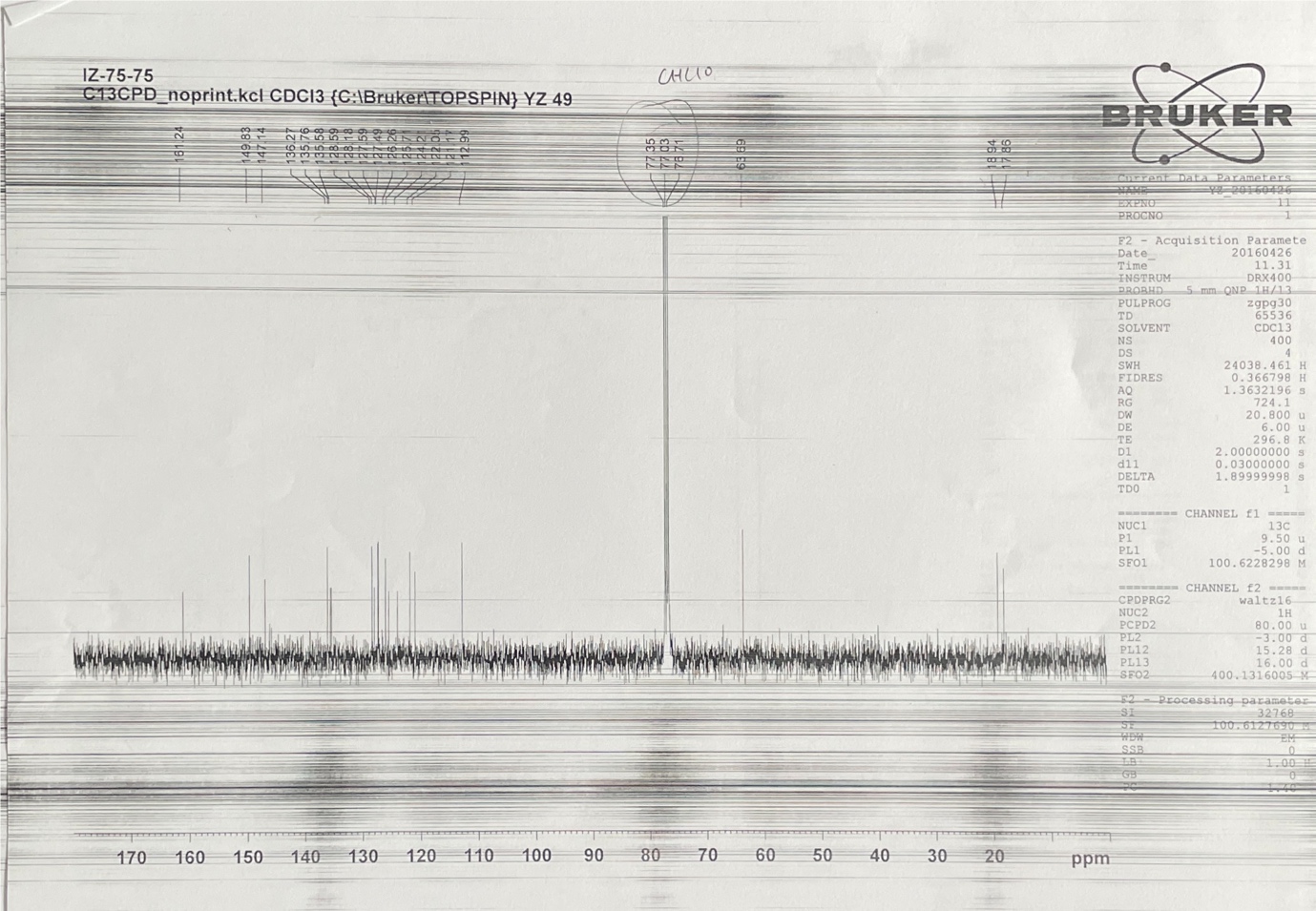

**^13^C-NMR of compound 6**

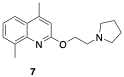

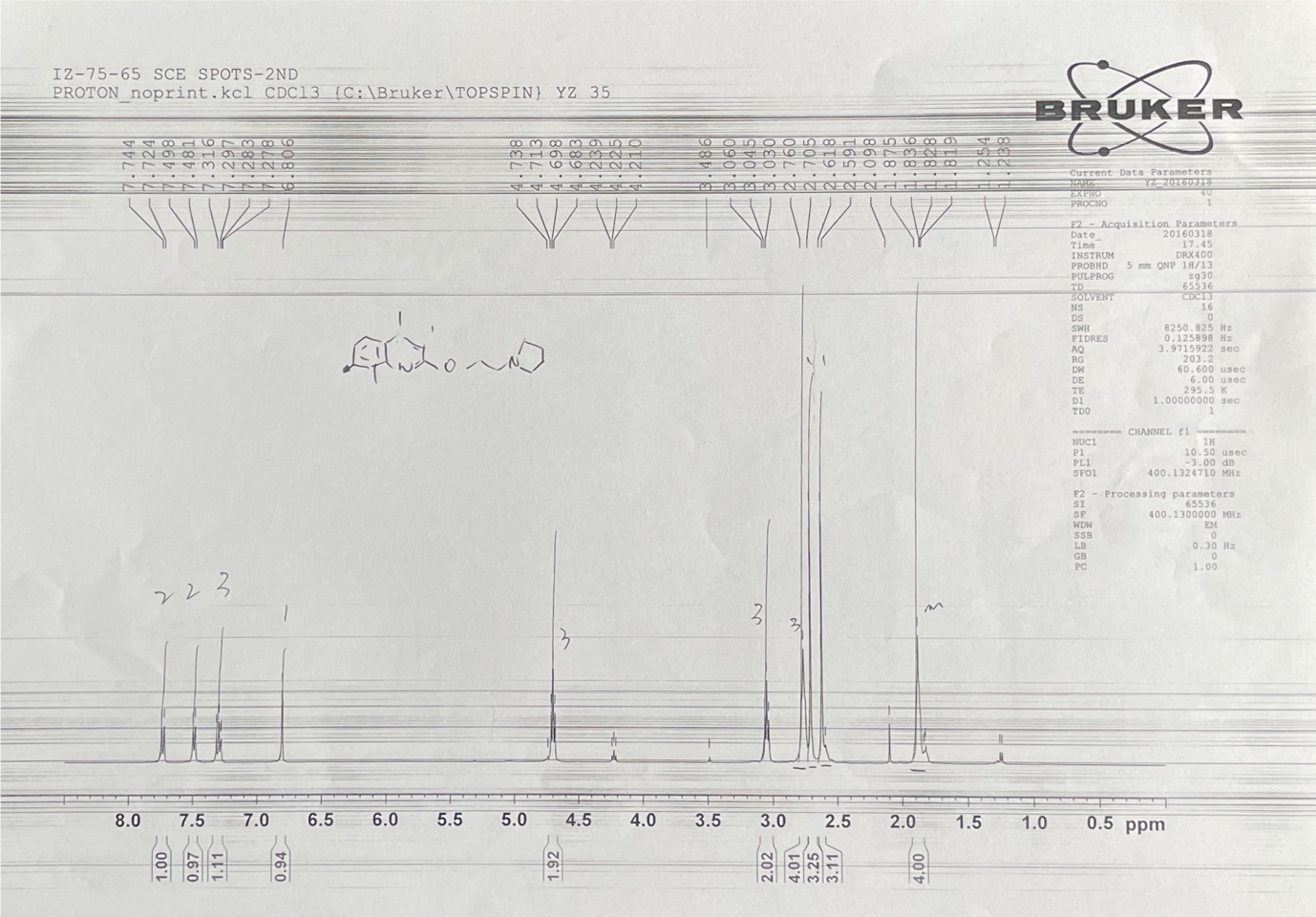

**^1^H-NMR of compound 7**

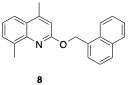

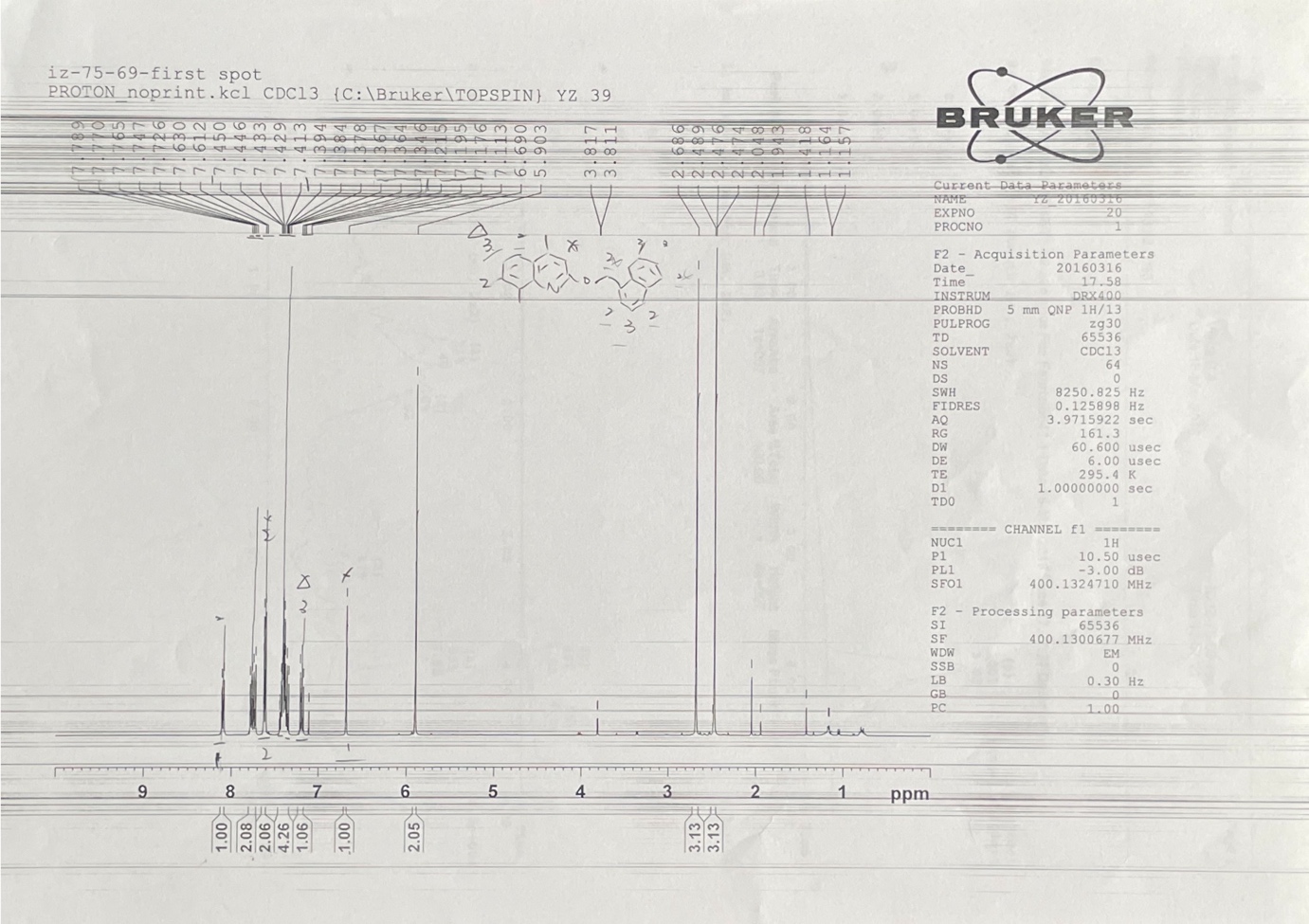

**^1^H-NMR of compound 8**

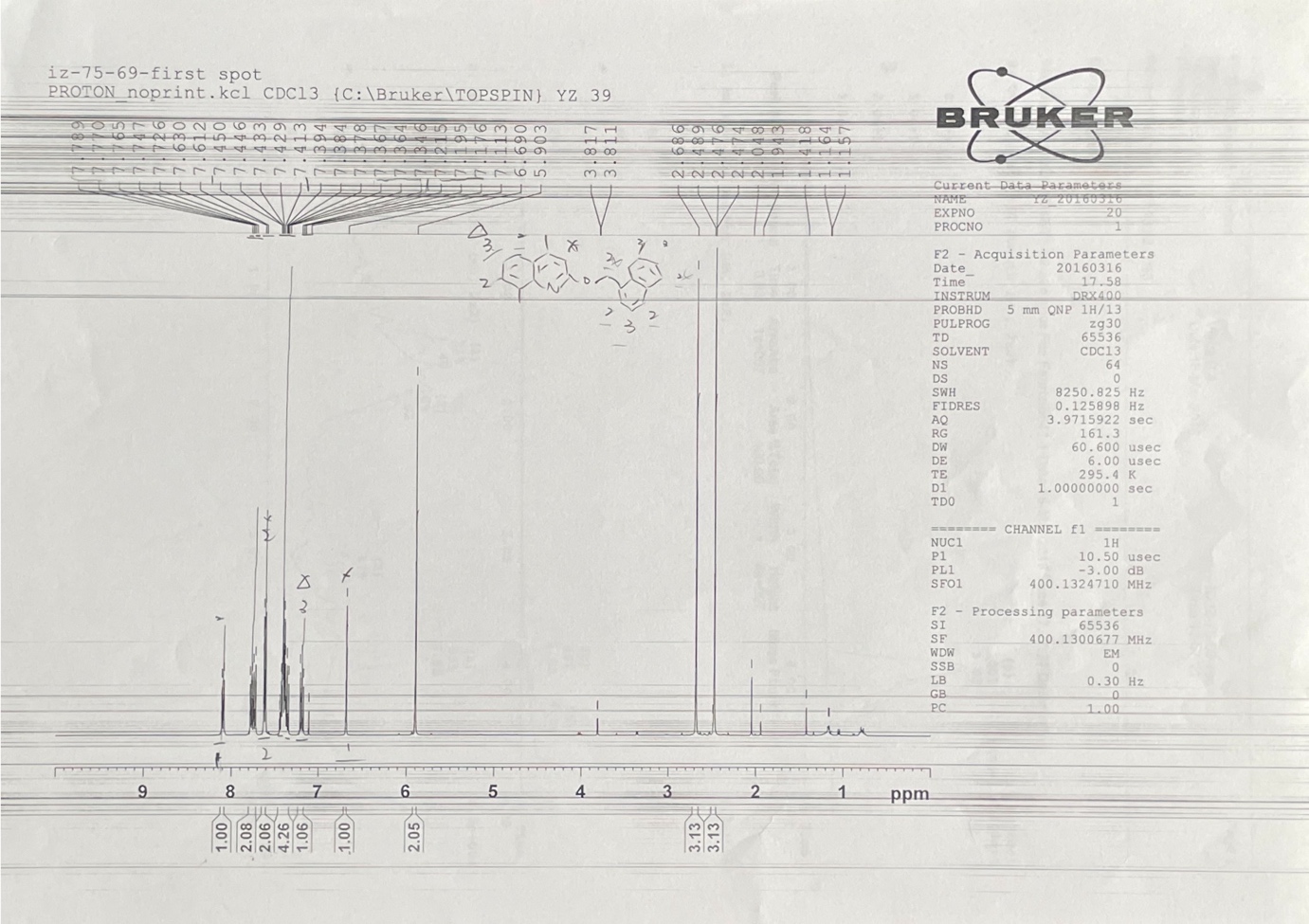

**^13^C-NMR of compound 8**

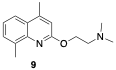

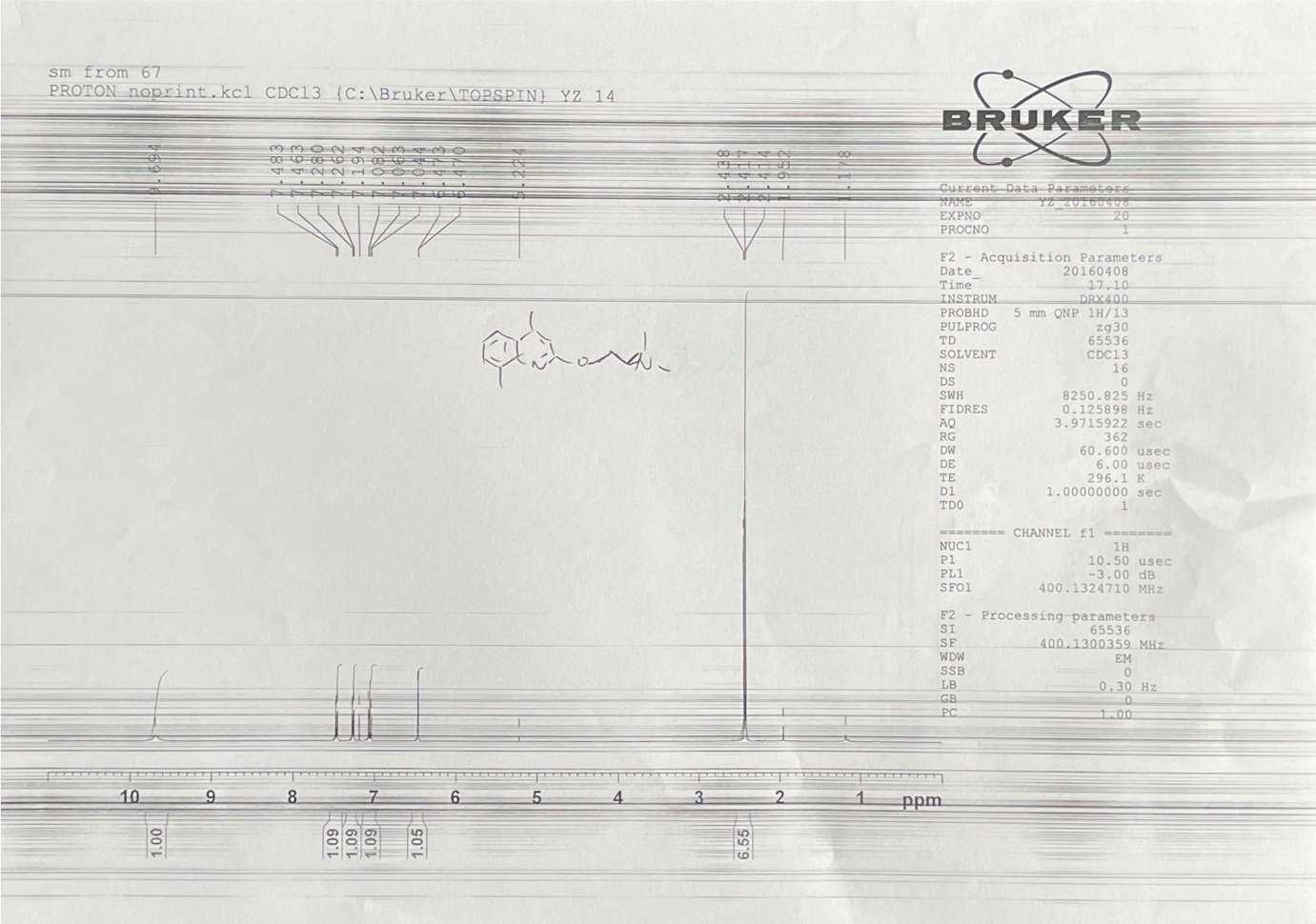

**^1^H-NMR of compound 9**

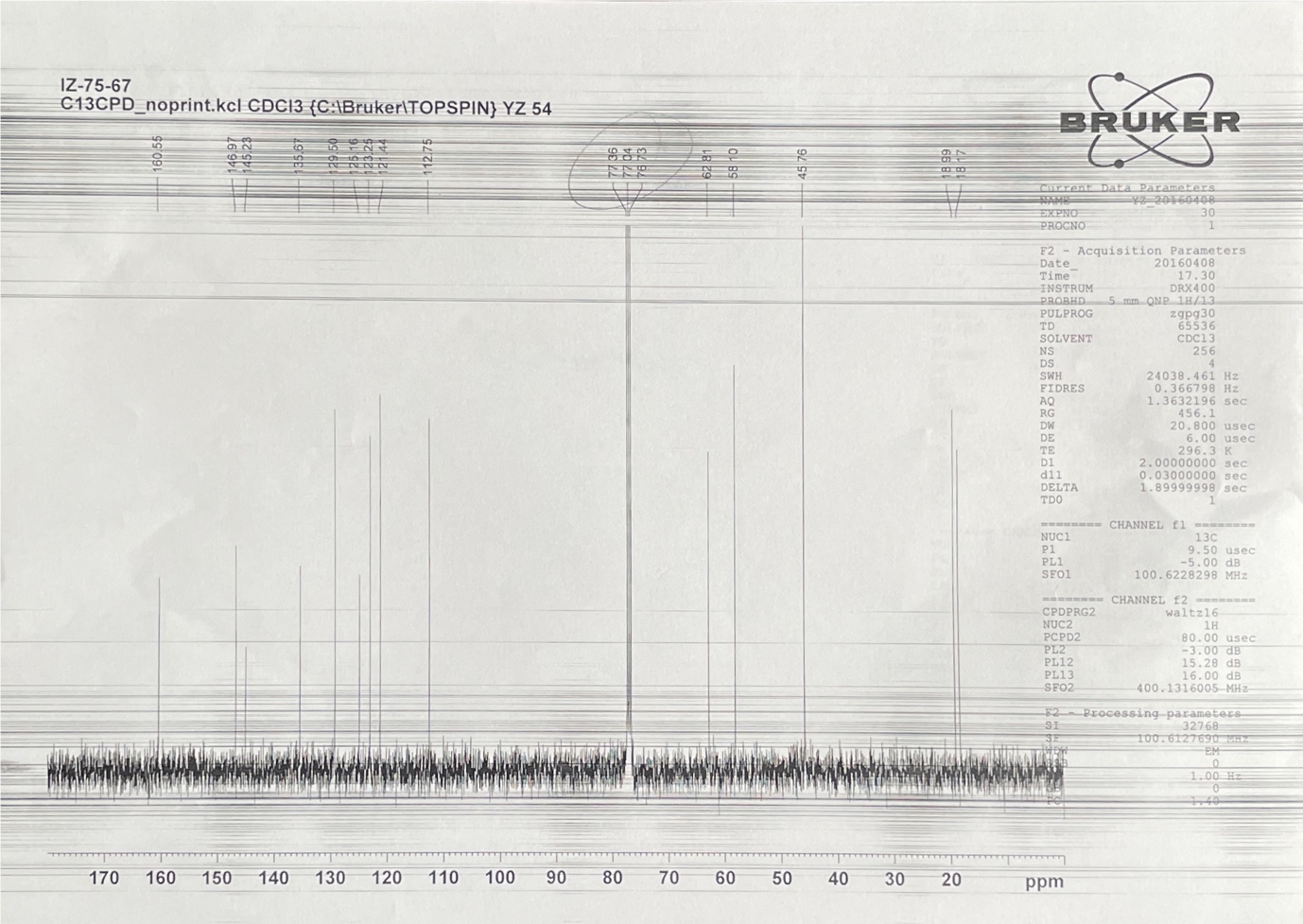

**^13^C-NMR of compound 9**

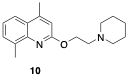

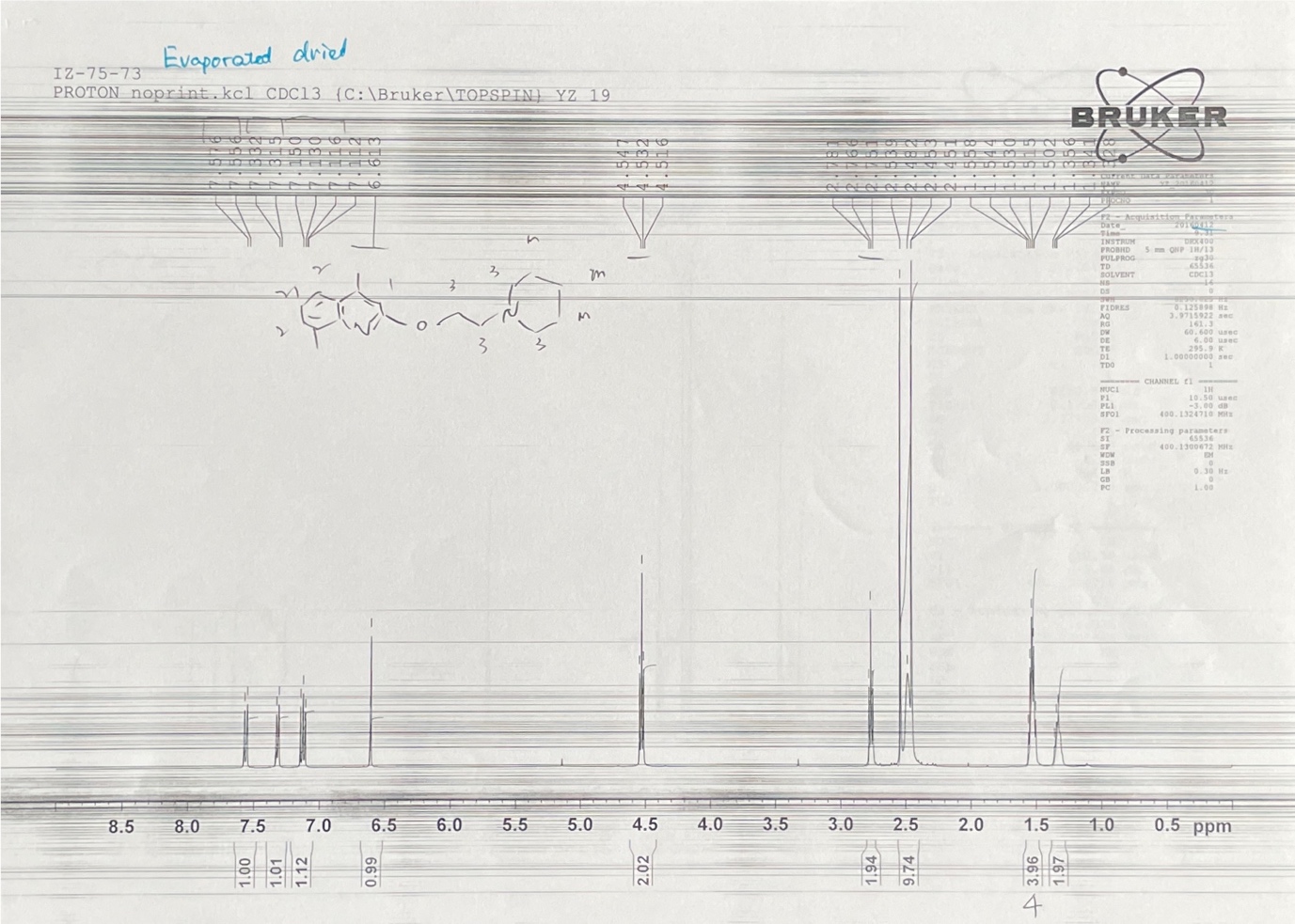

**^1^H-NMR of compound 10**

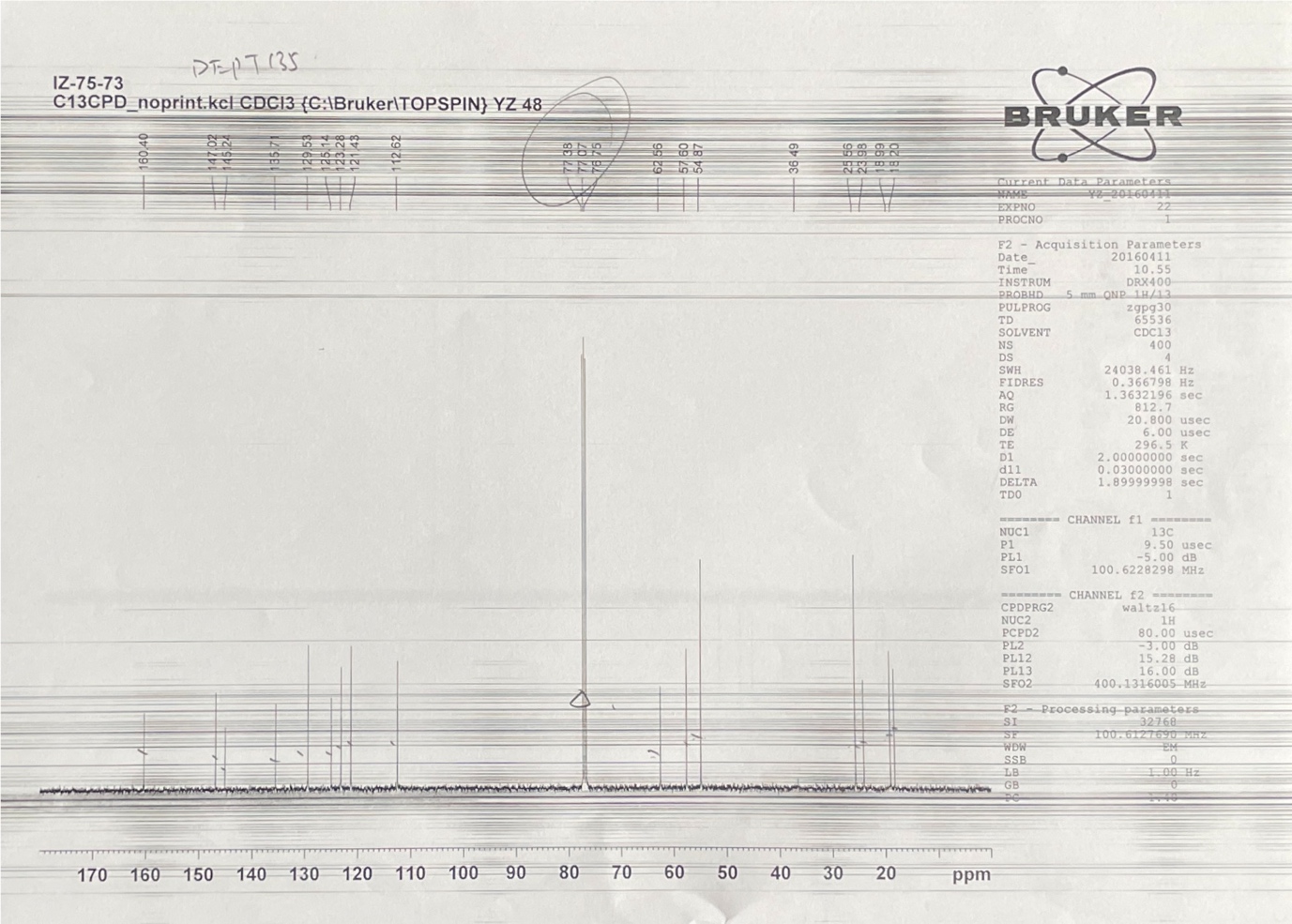

**^13^C-NMR of compound 10**

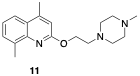

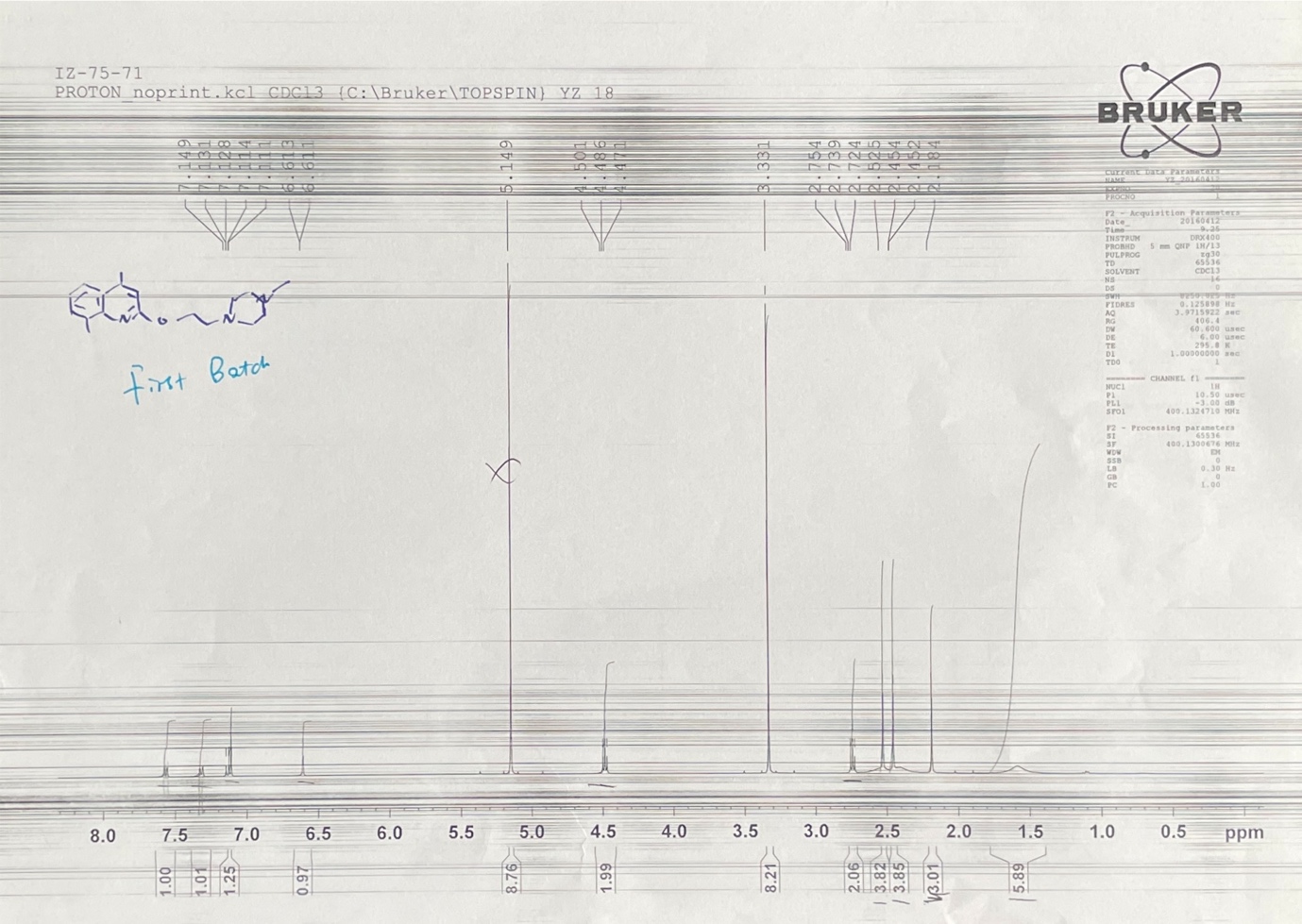

**^1^H-NMR of compound 11**

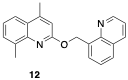

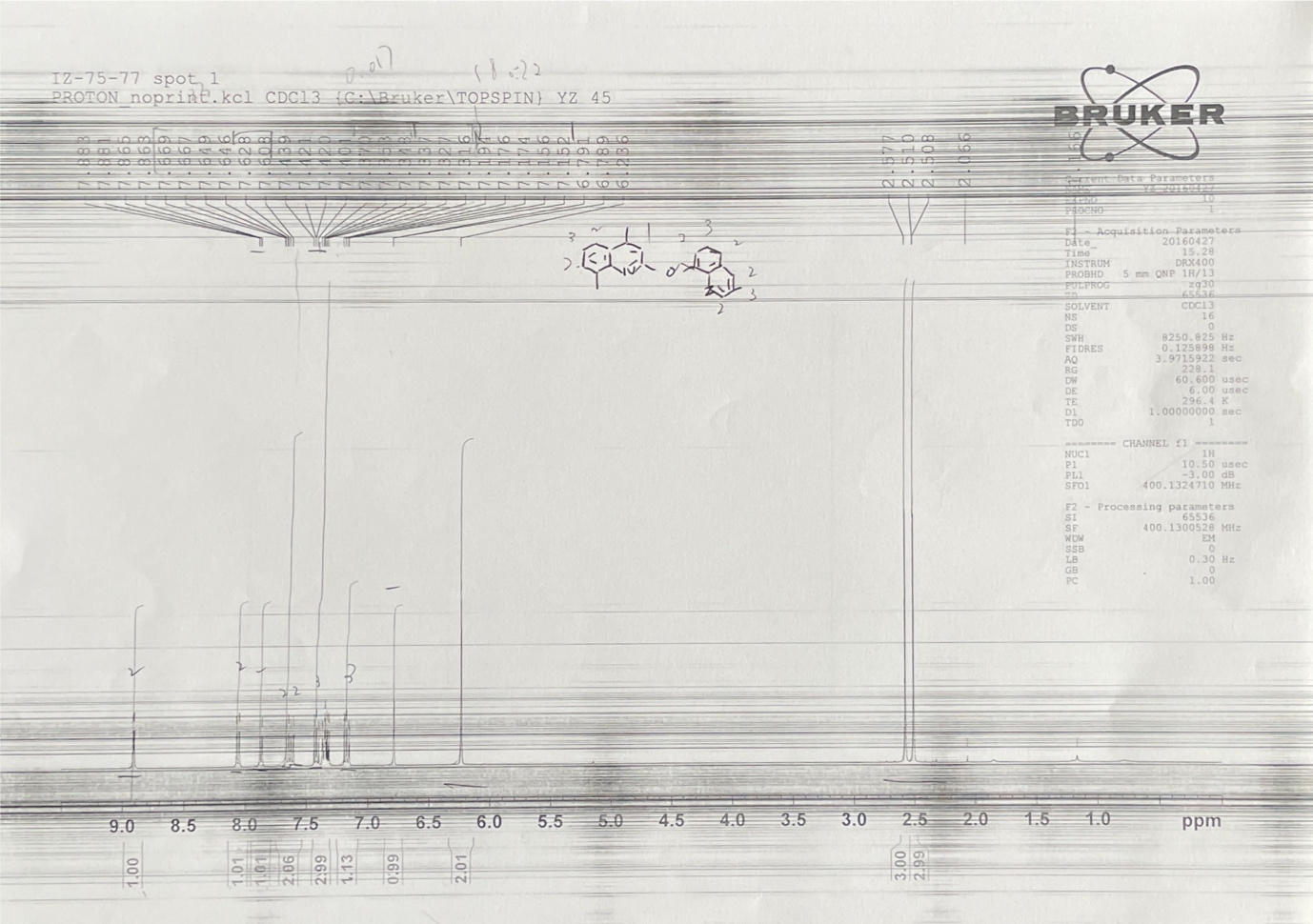

**^1^H-NMR of compound 12**

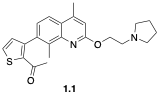

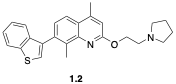

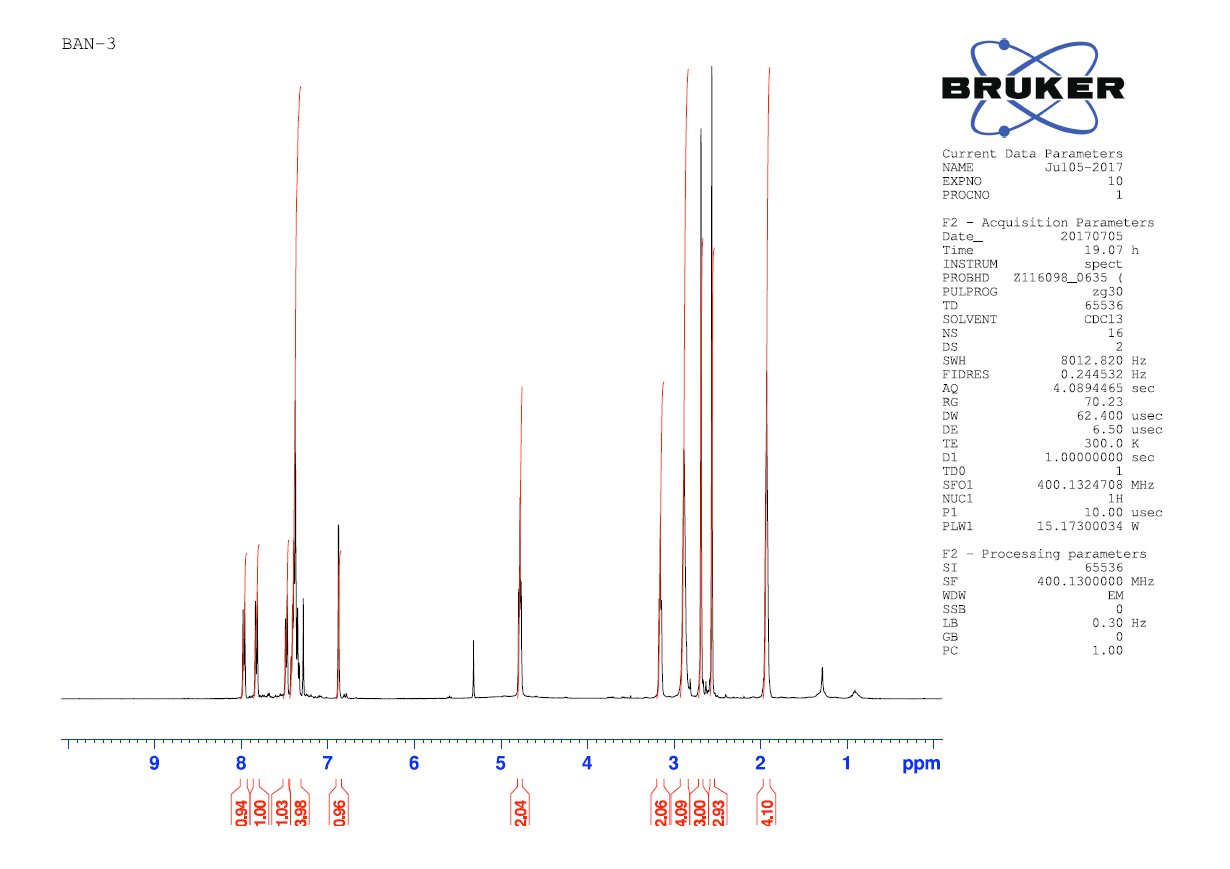

**Figure S24. ^1^H-NMR of compound 1.2**

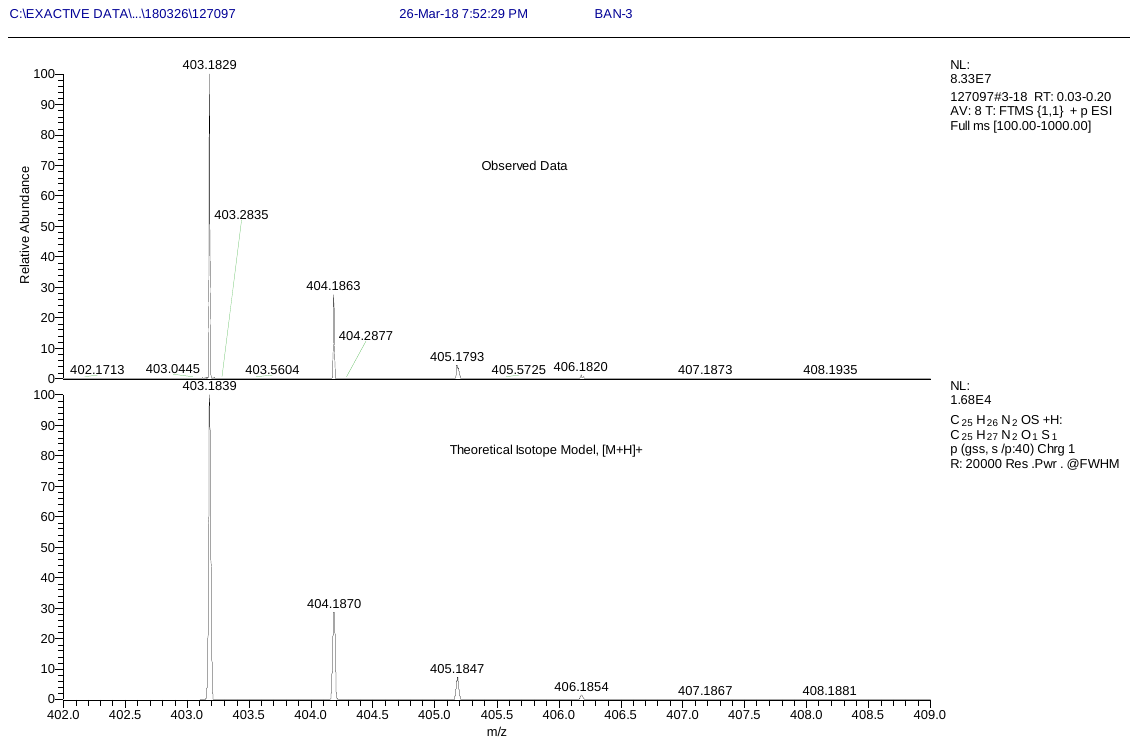

**HR-MS of compound 1.2**

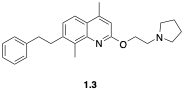

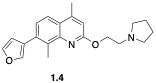

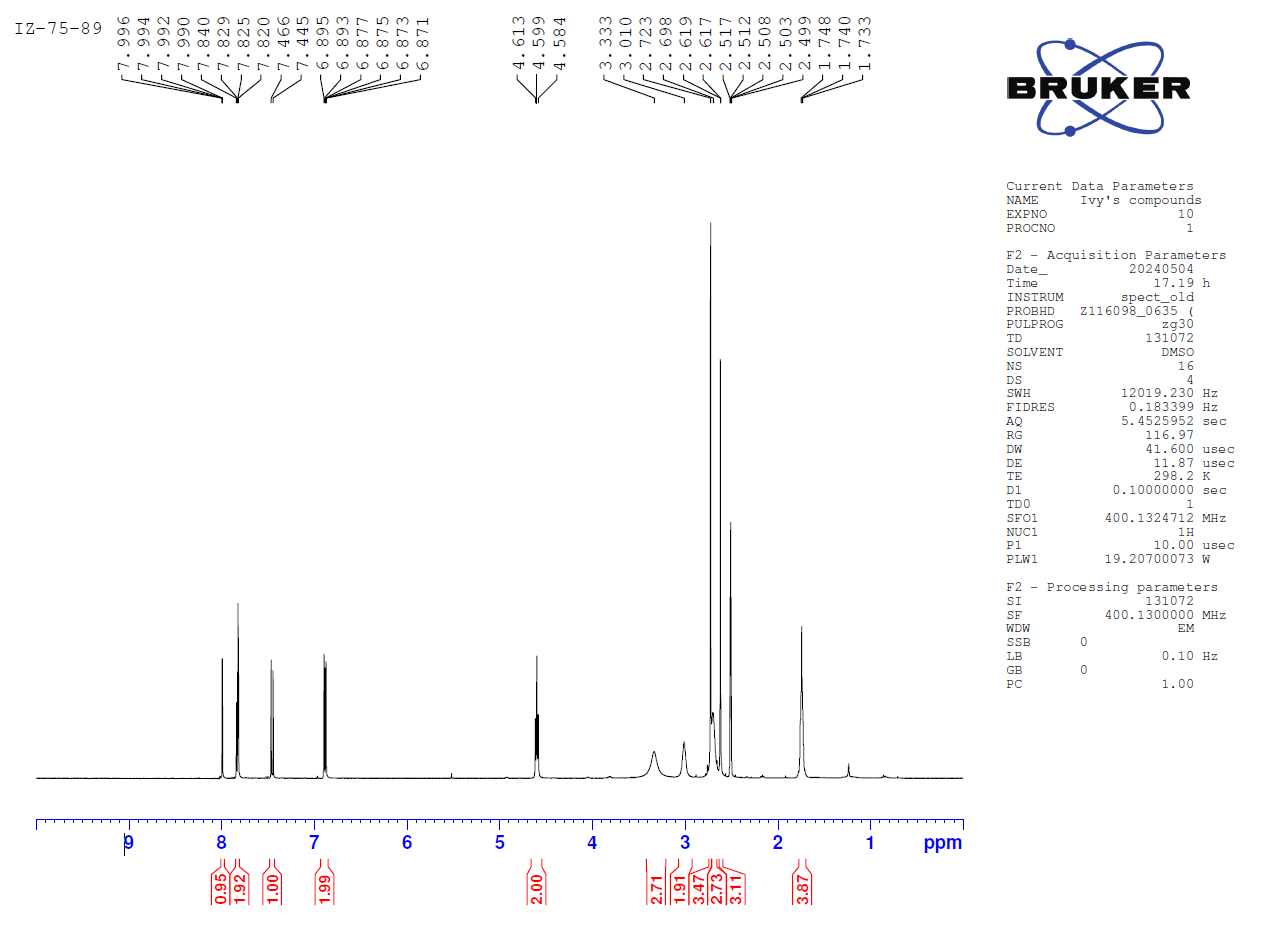

**^1^H-NMR of compound 1.4**

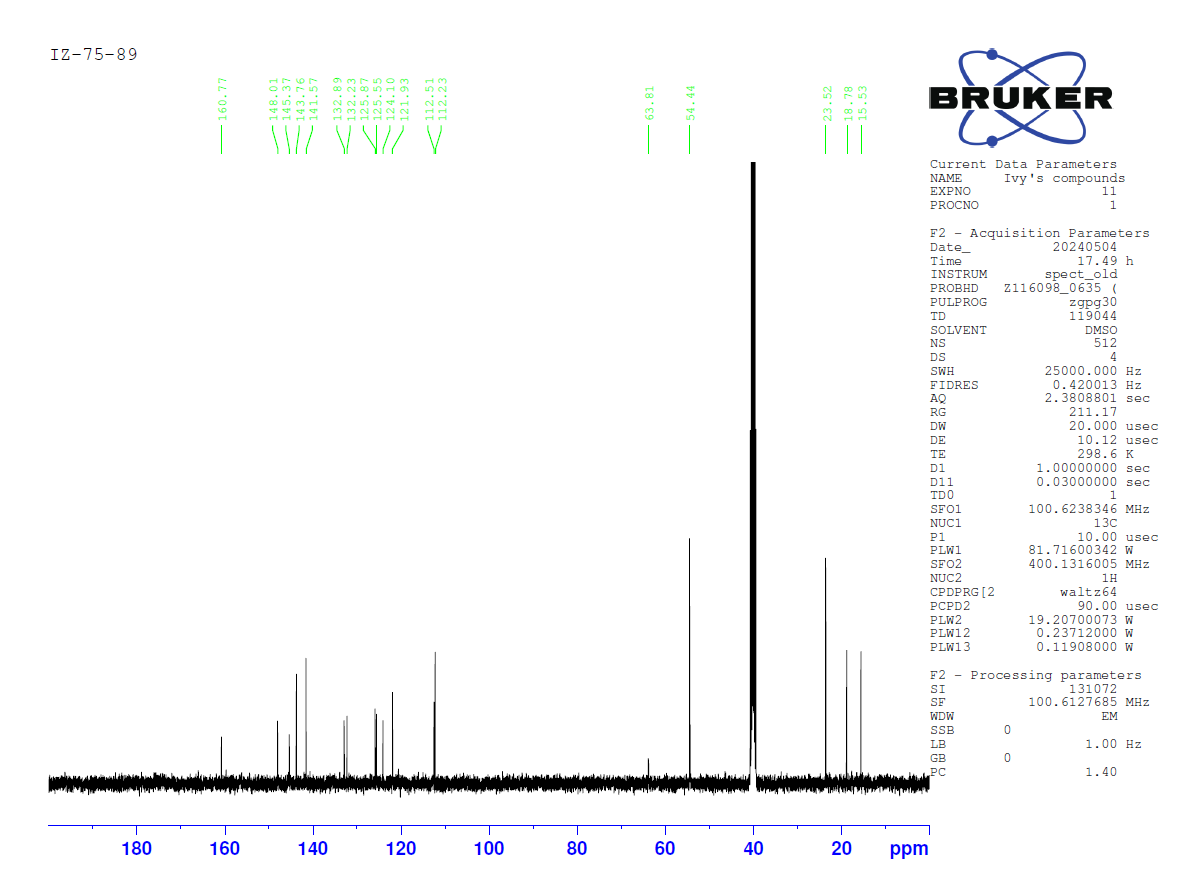

**^13^C-NMR of compound 1.4**

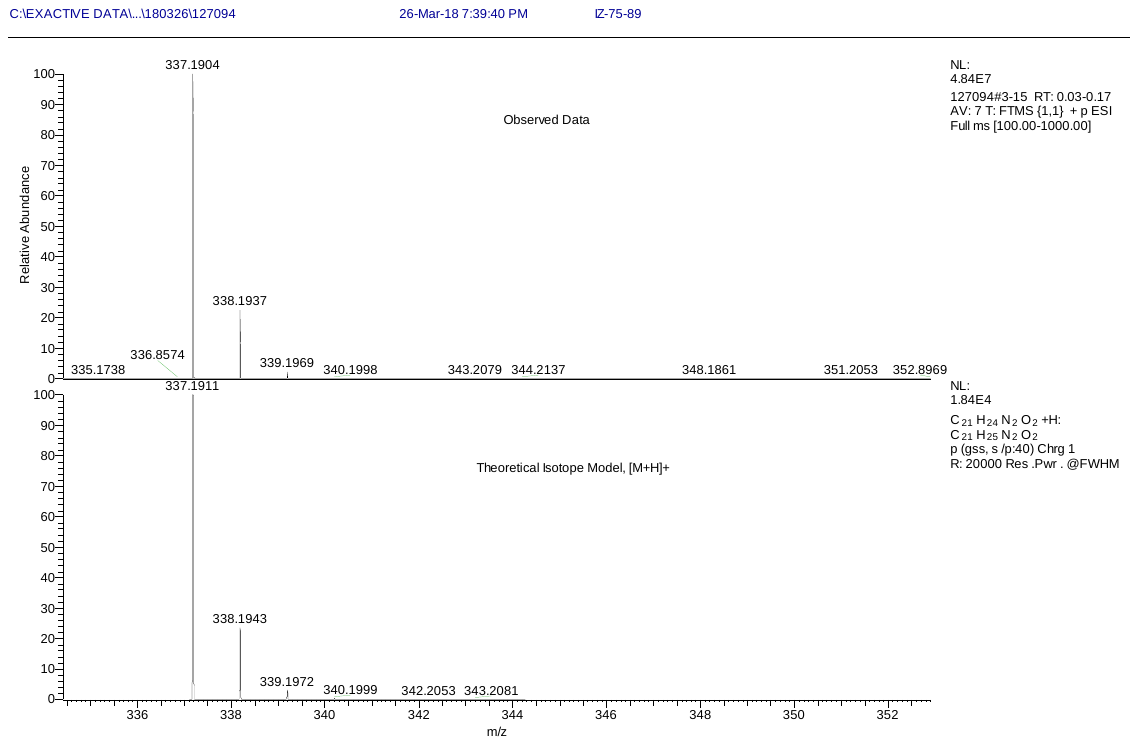

**HR-MS of compound 1.4**

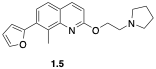

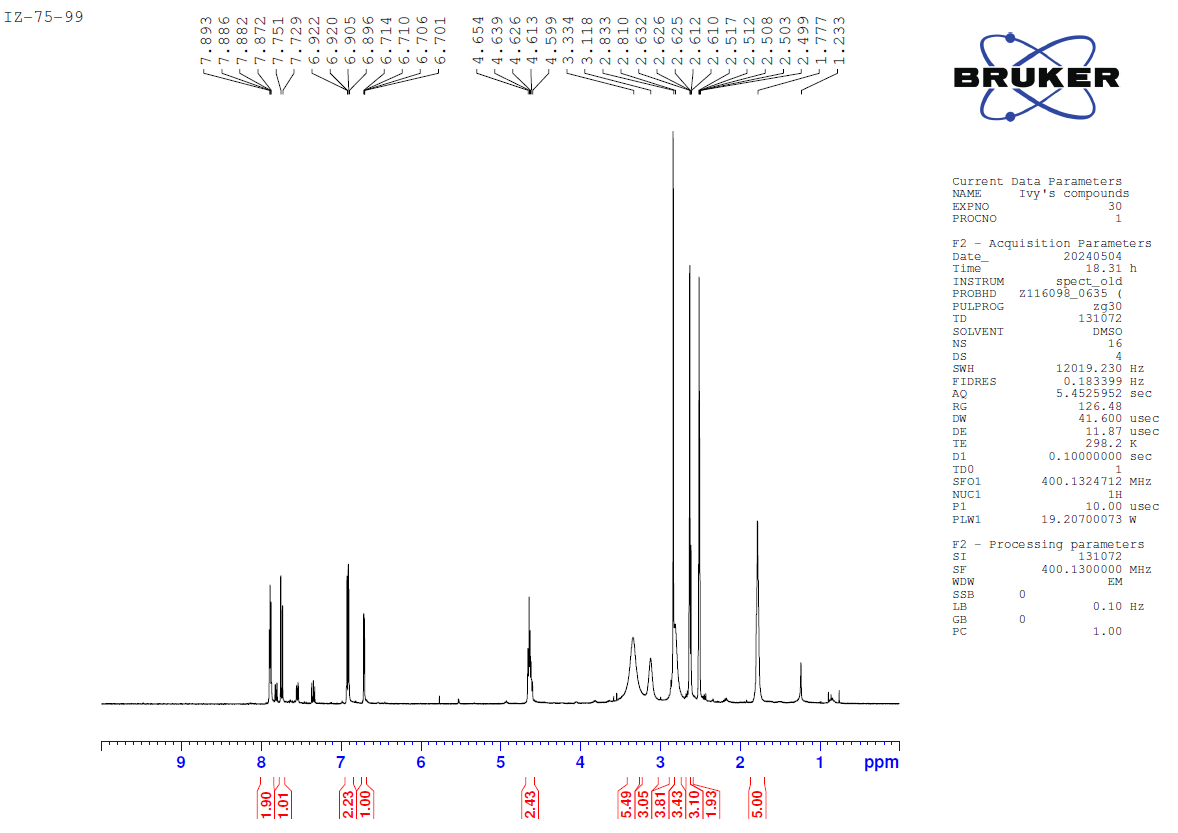

**^1^H-NMR of compound 1.5**

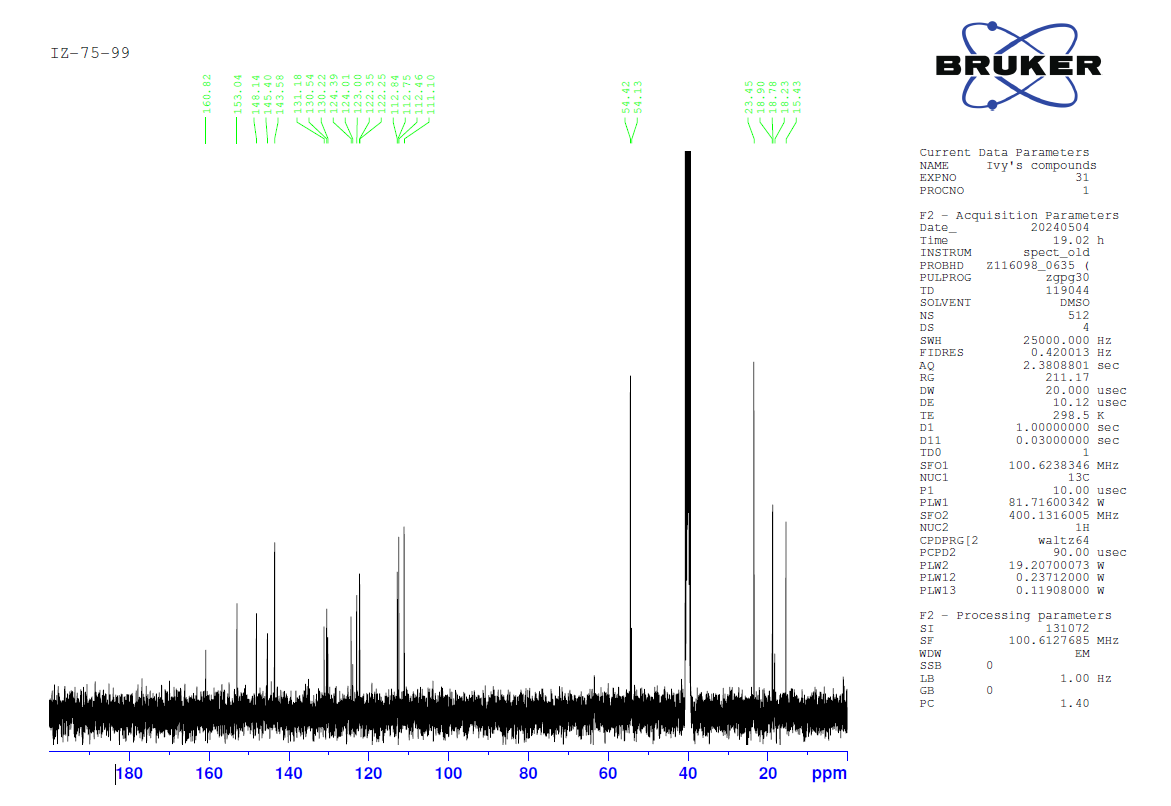

**^13^C-NMR of compound 1.5**

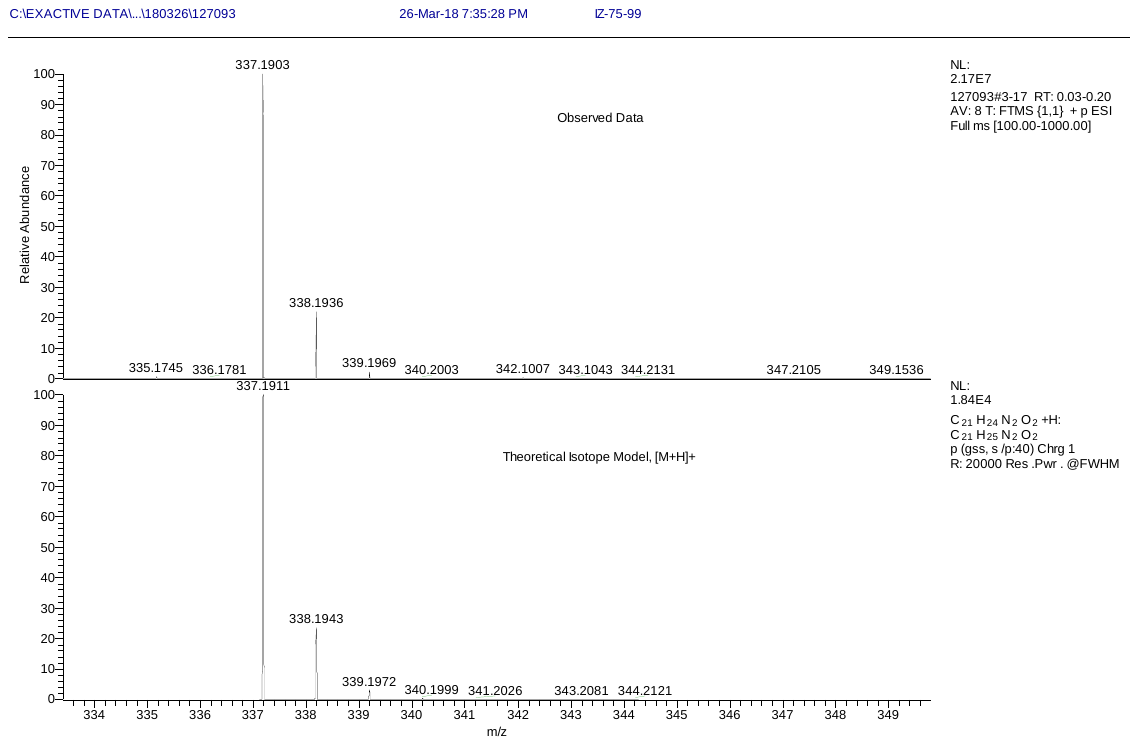

**HR-MS of compound 1.5**

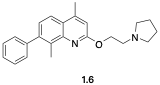

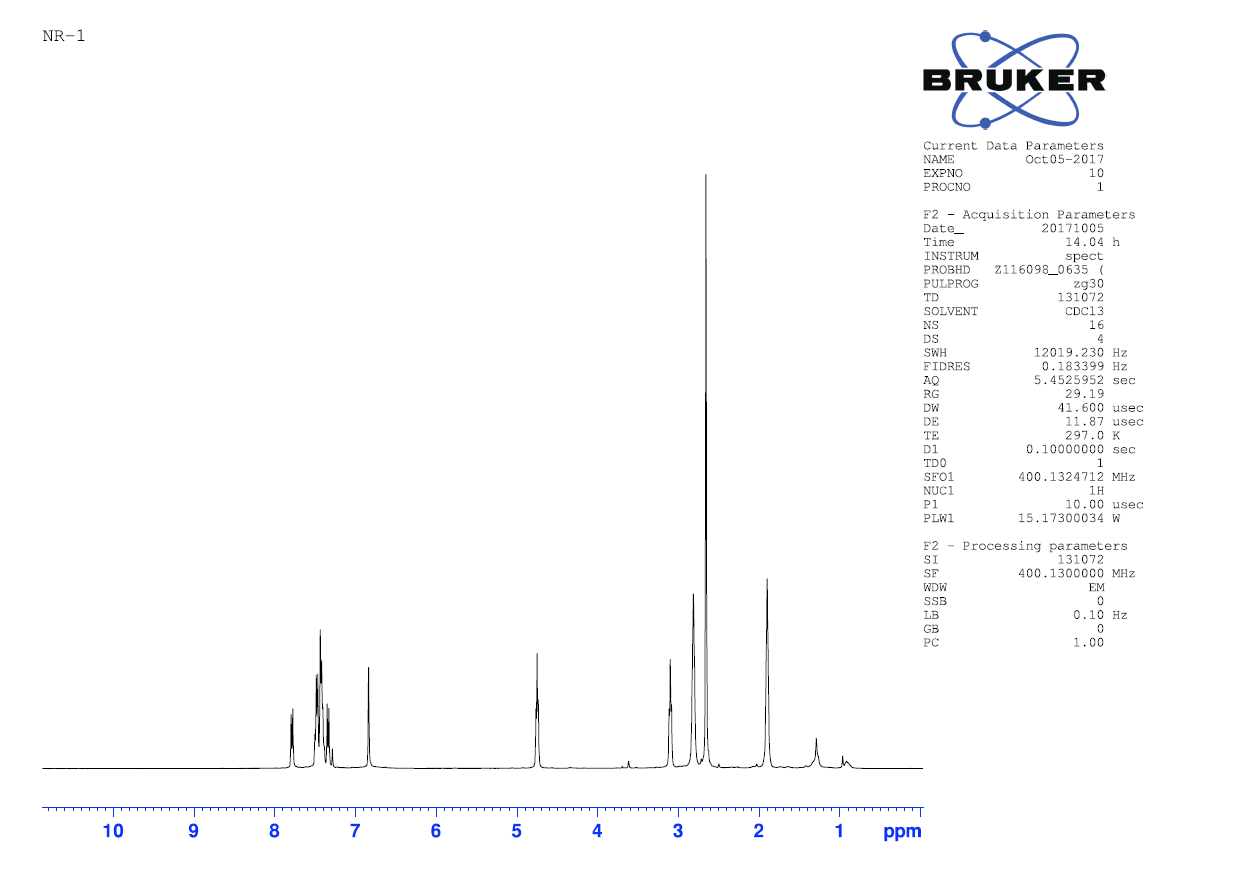
 **^1^H-NMR of compound 1.6**

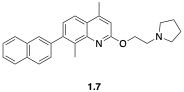

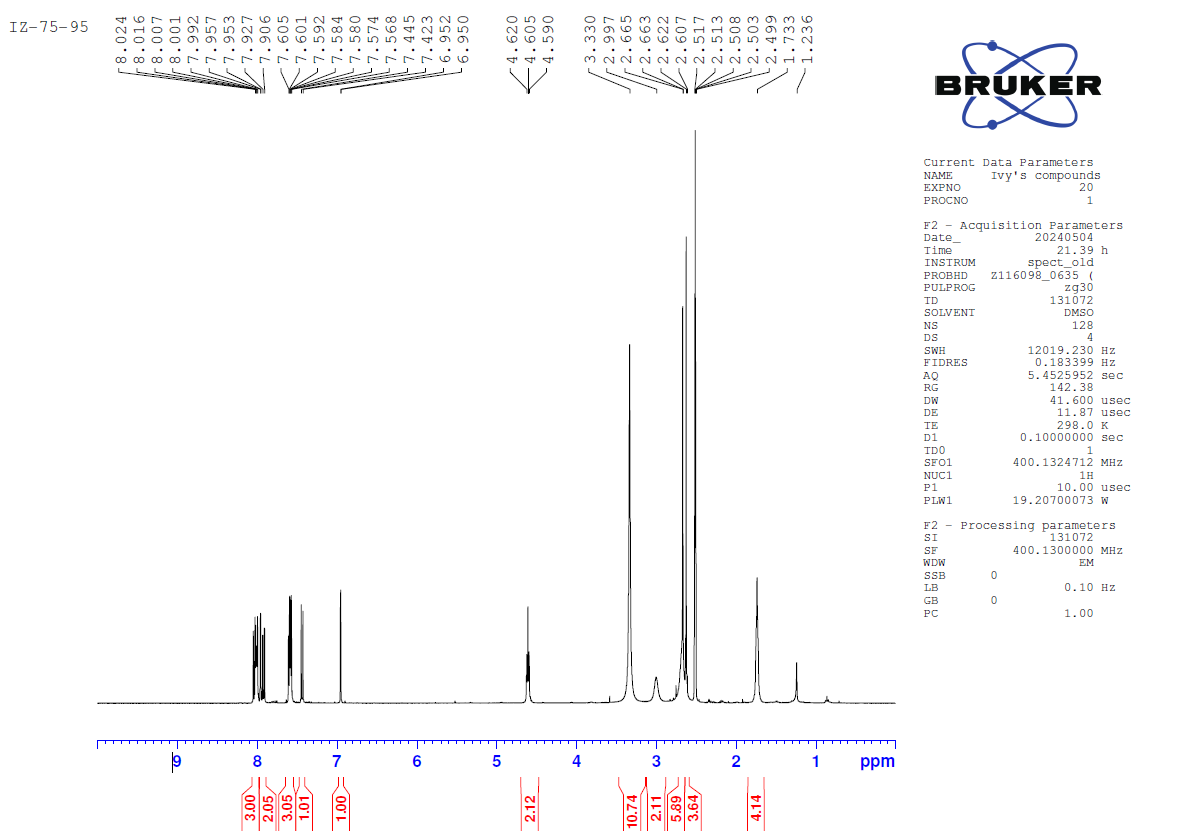

**^1^H-NMR of compound 1.7**

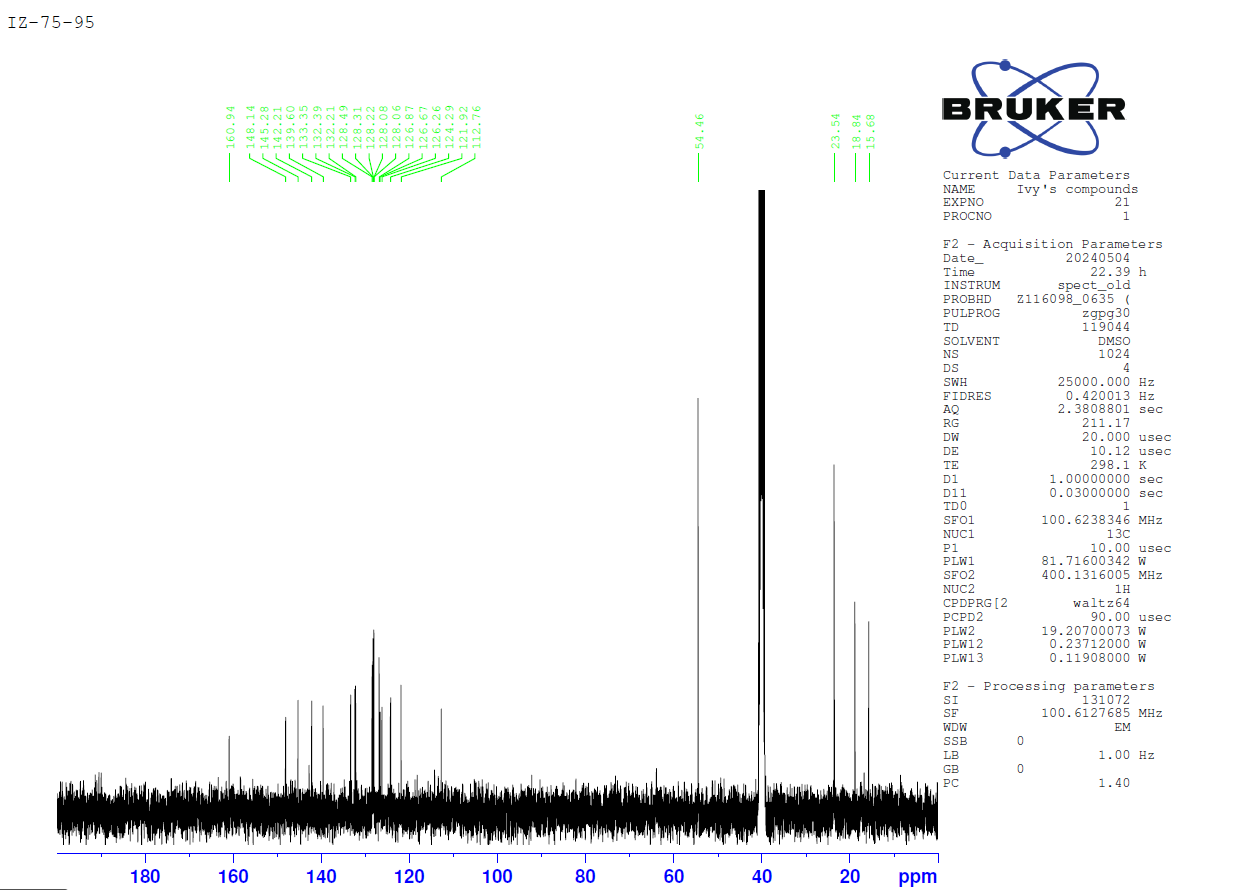

**^13^C-NMR of compound 1.7**

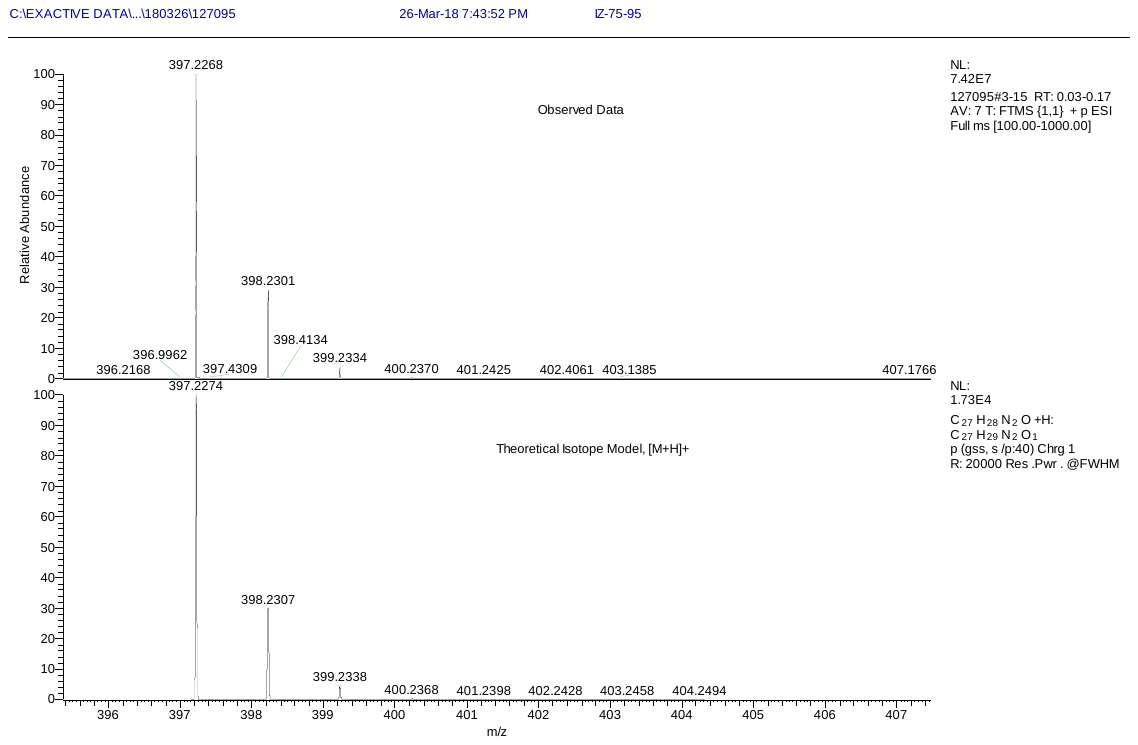

**HR-MS of compound 1.7**

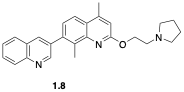

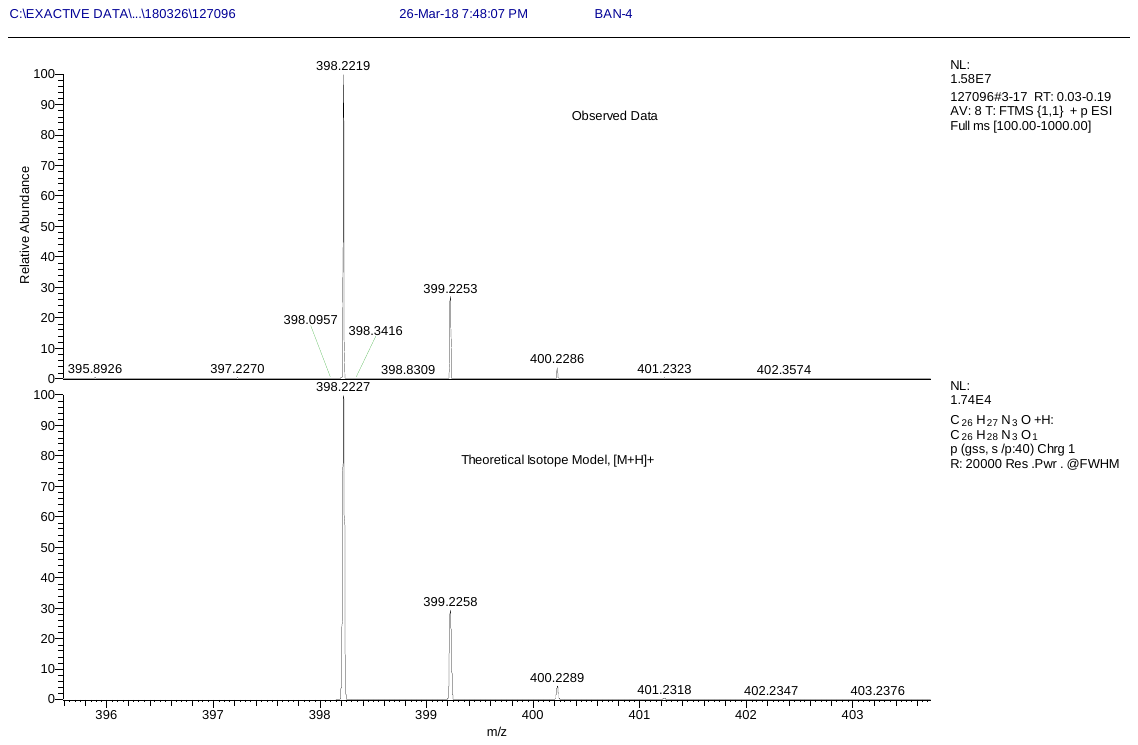

**HR-MS of compound 1.8**

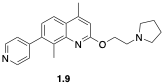

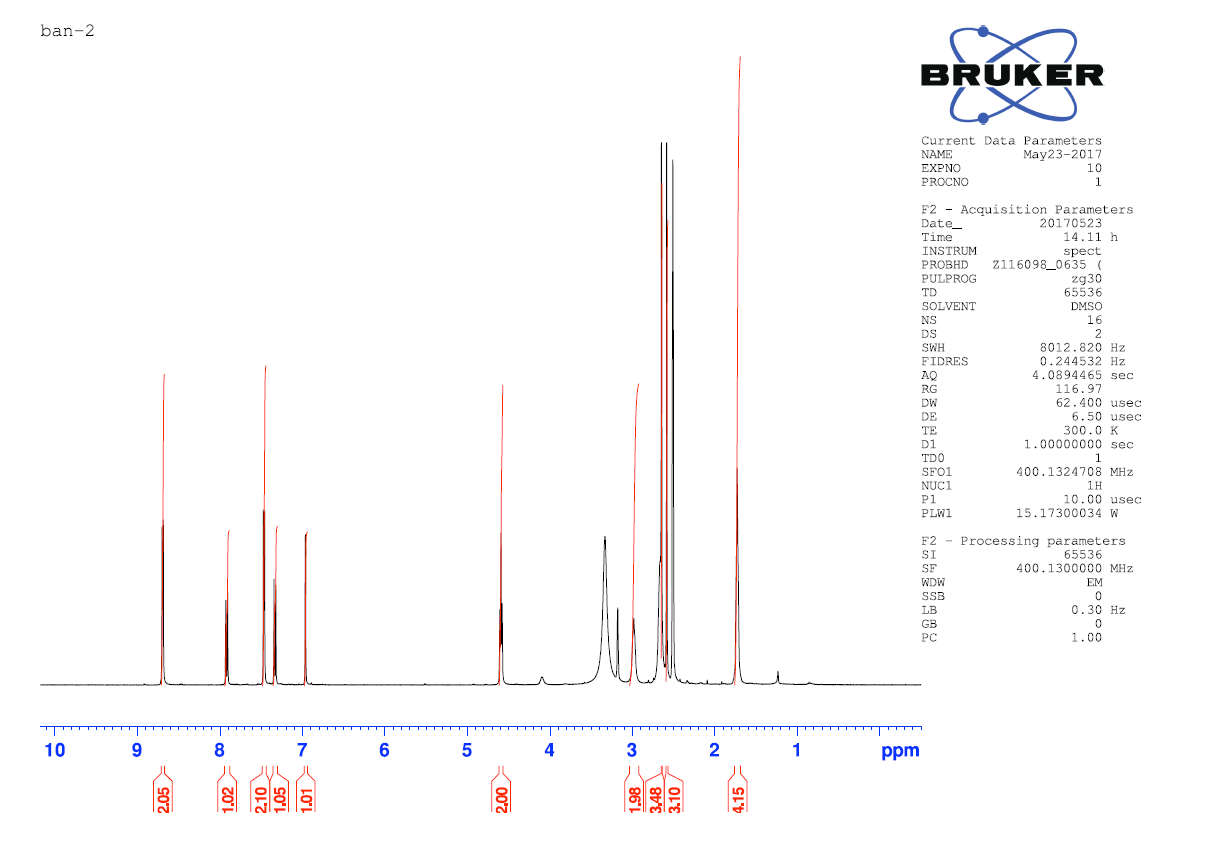

**^1^H-NMR of compound 1.9**

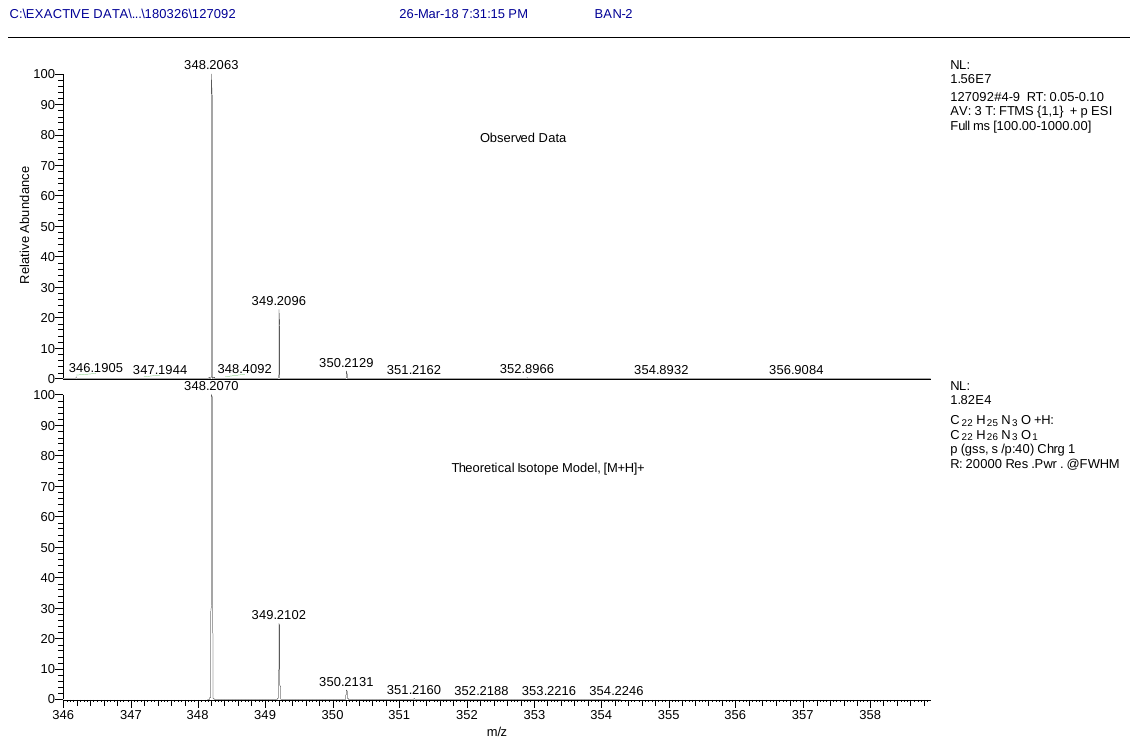

**HR-MS of compound 1.9**

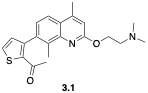

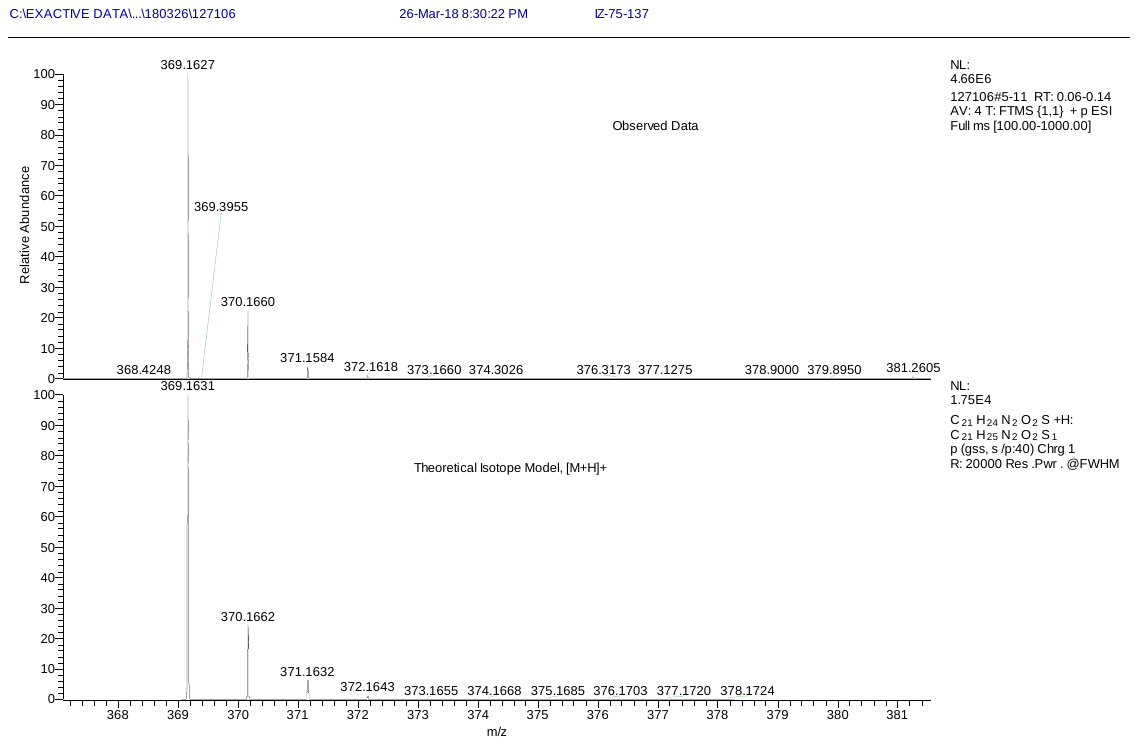
 **HR-MS of compound 3.1**

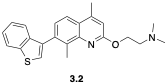

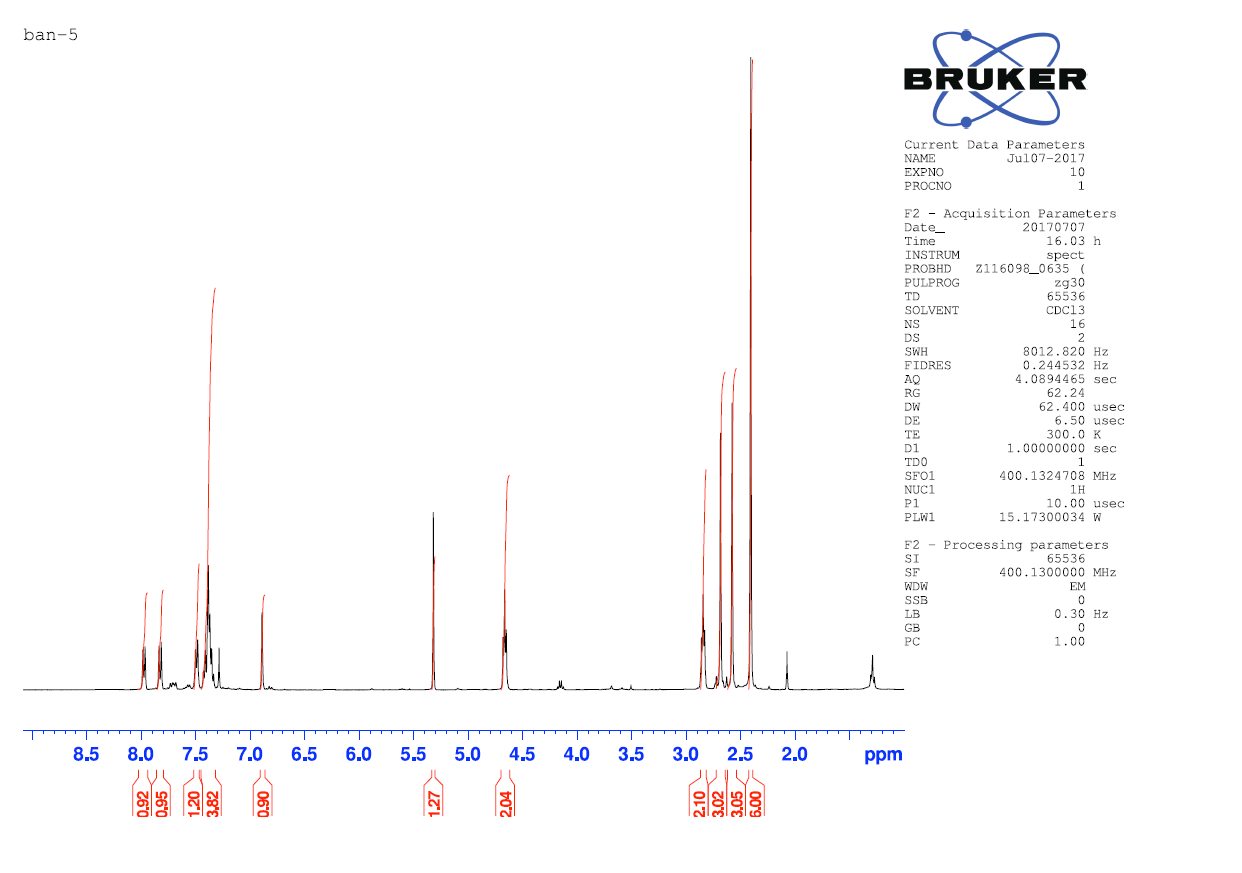

**^1^H-NMR of compound 3.2**

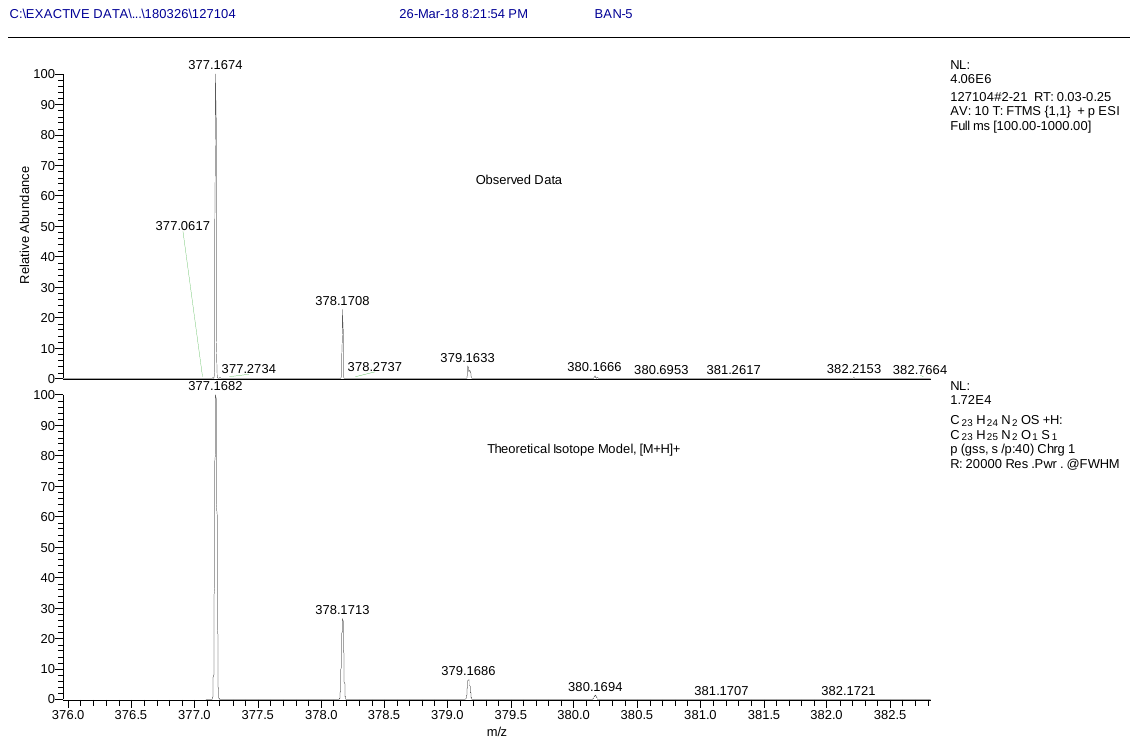

**HR-MS of compound 3.1**

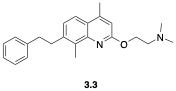

**3.4**

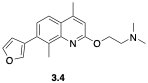

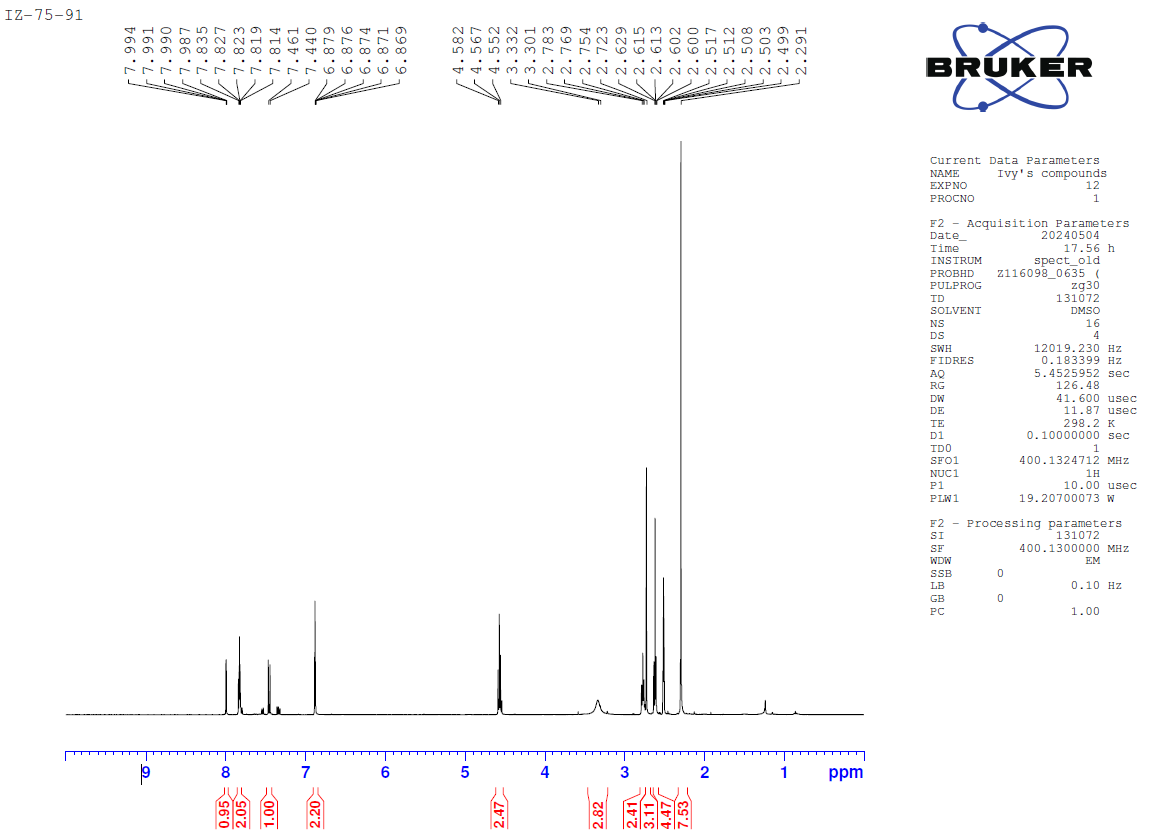

**^1^H-NMR of compound 3.4**

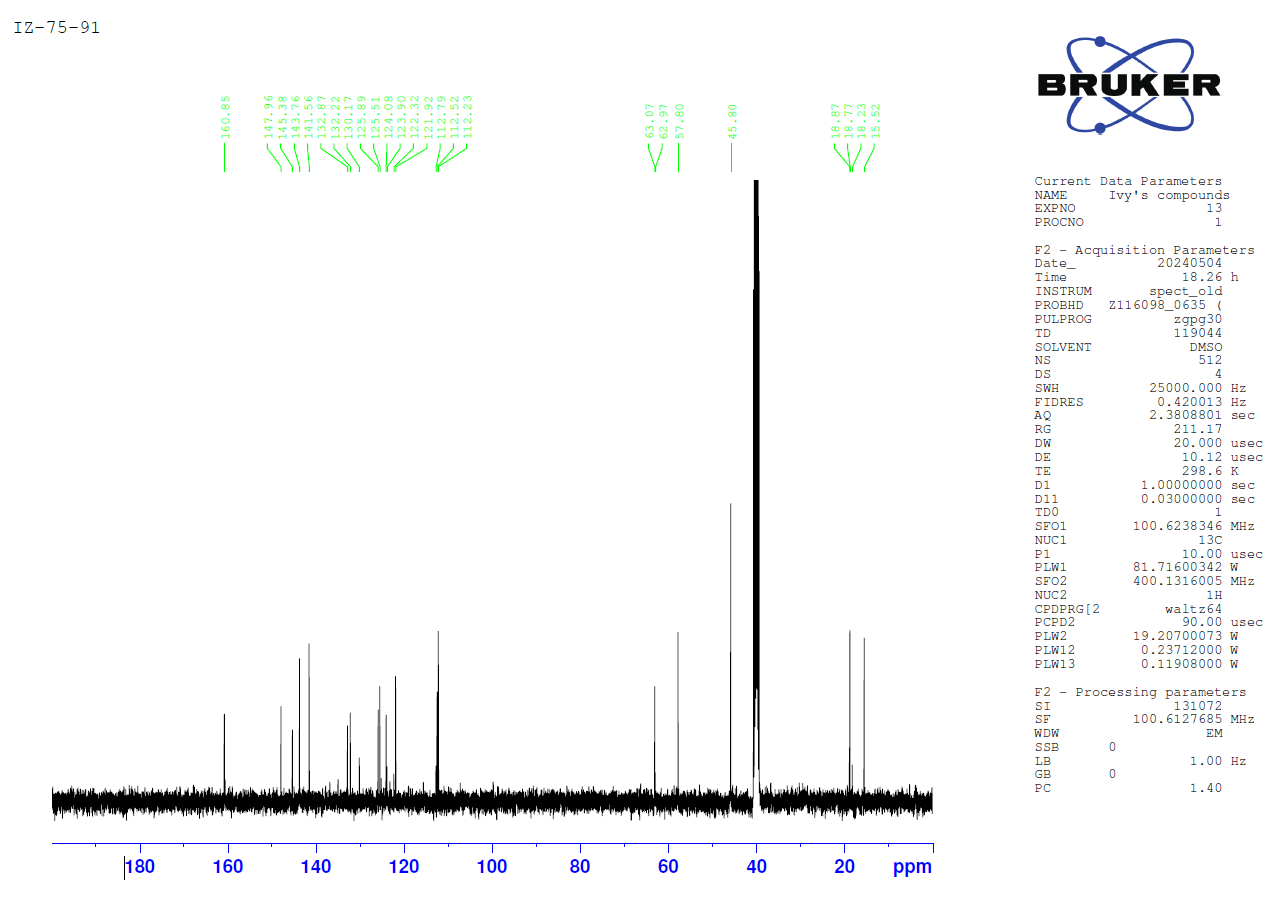

**^13^C-NMR of compound 3.4**

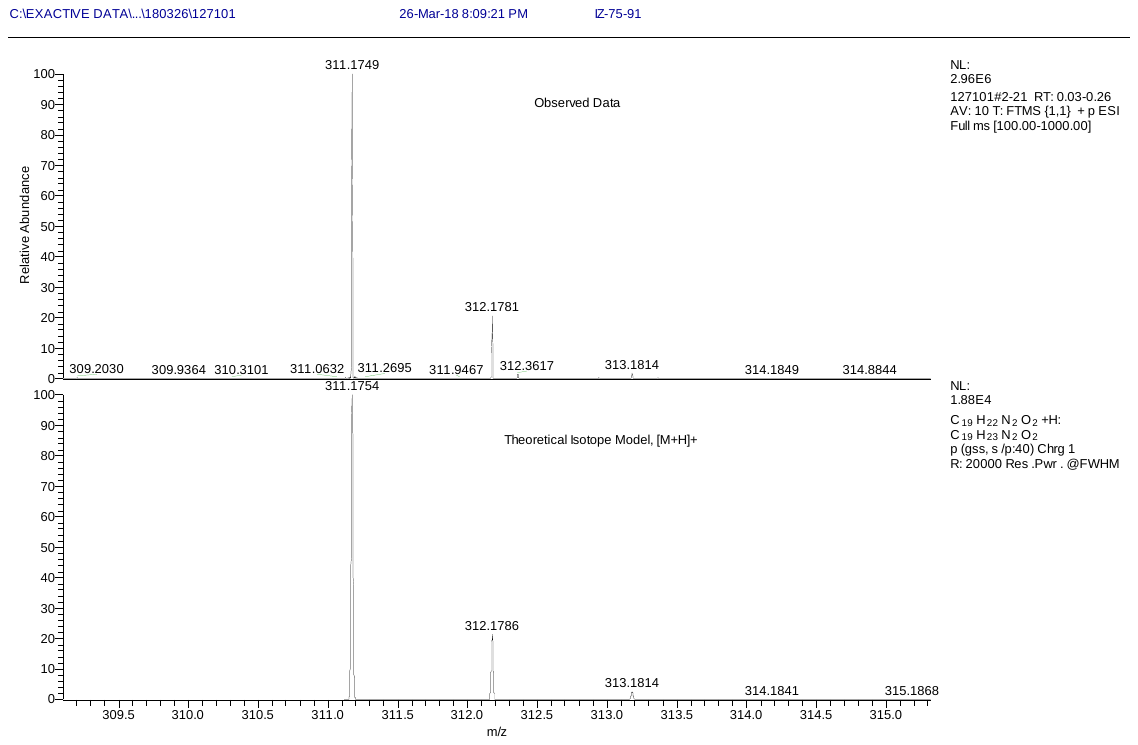

**HR-MS of compound 3.4**

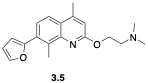

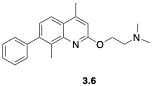

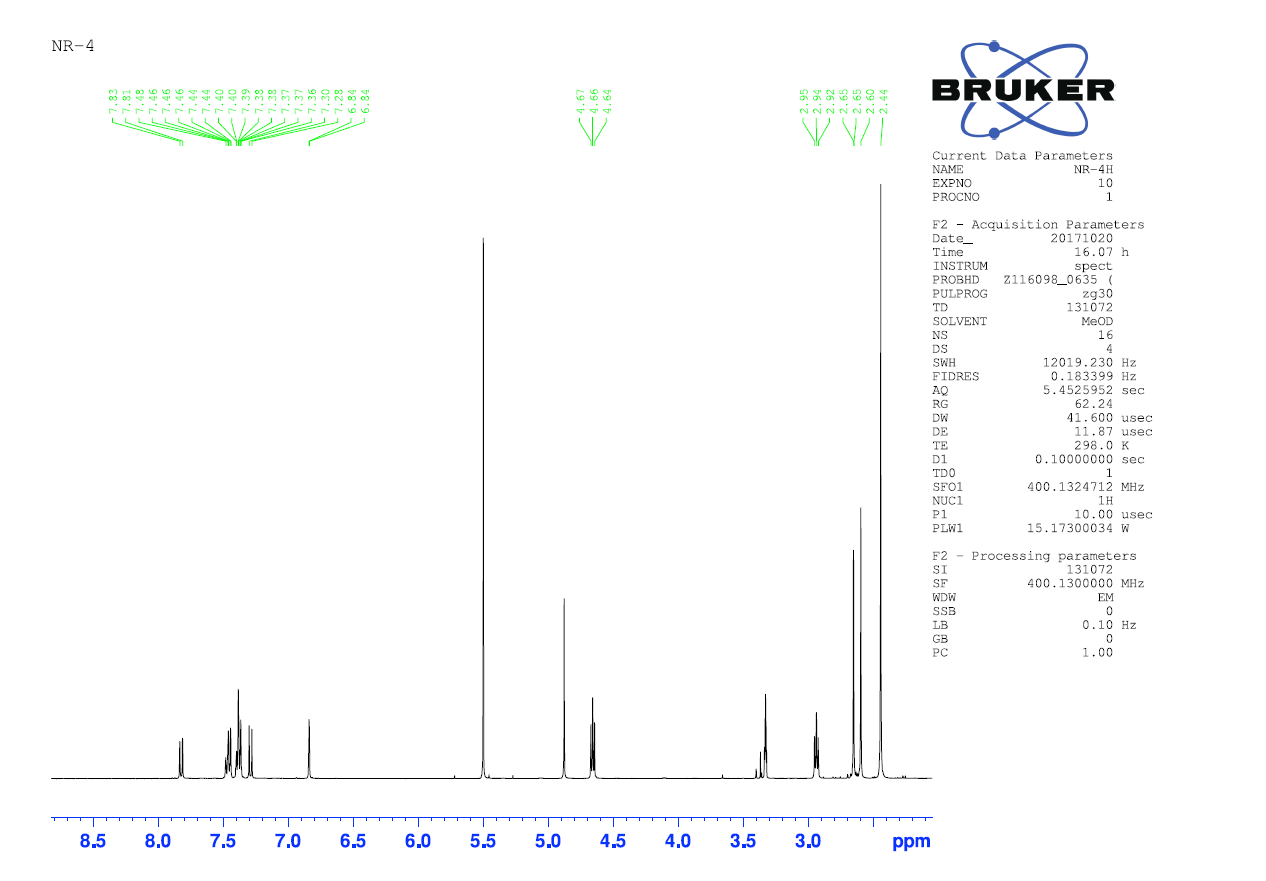

**^1^H-NMR of compound 3.6**

**3.7**

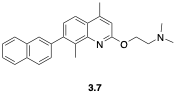

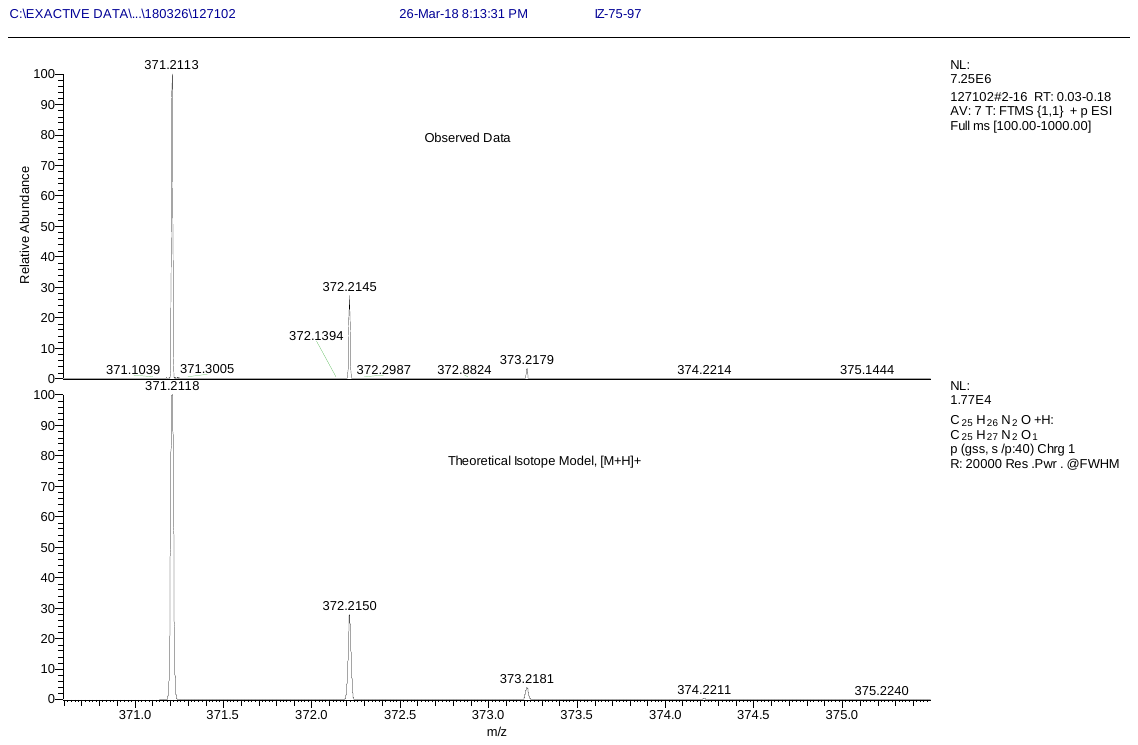

**HR-MS of compound 3.4**

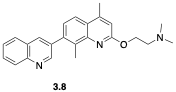

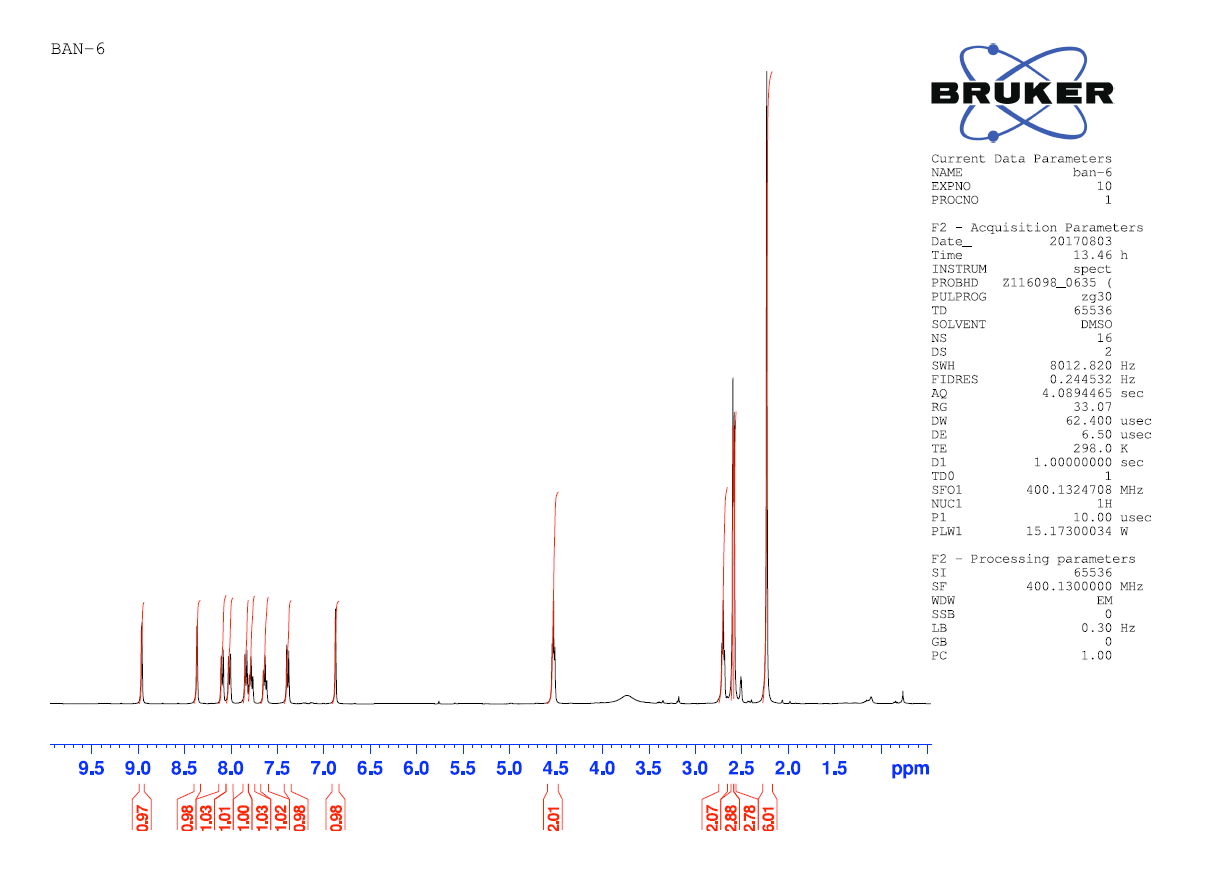

**^1^H-NMR of compound 3.8**

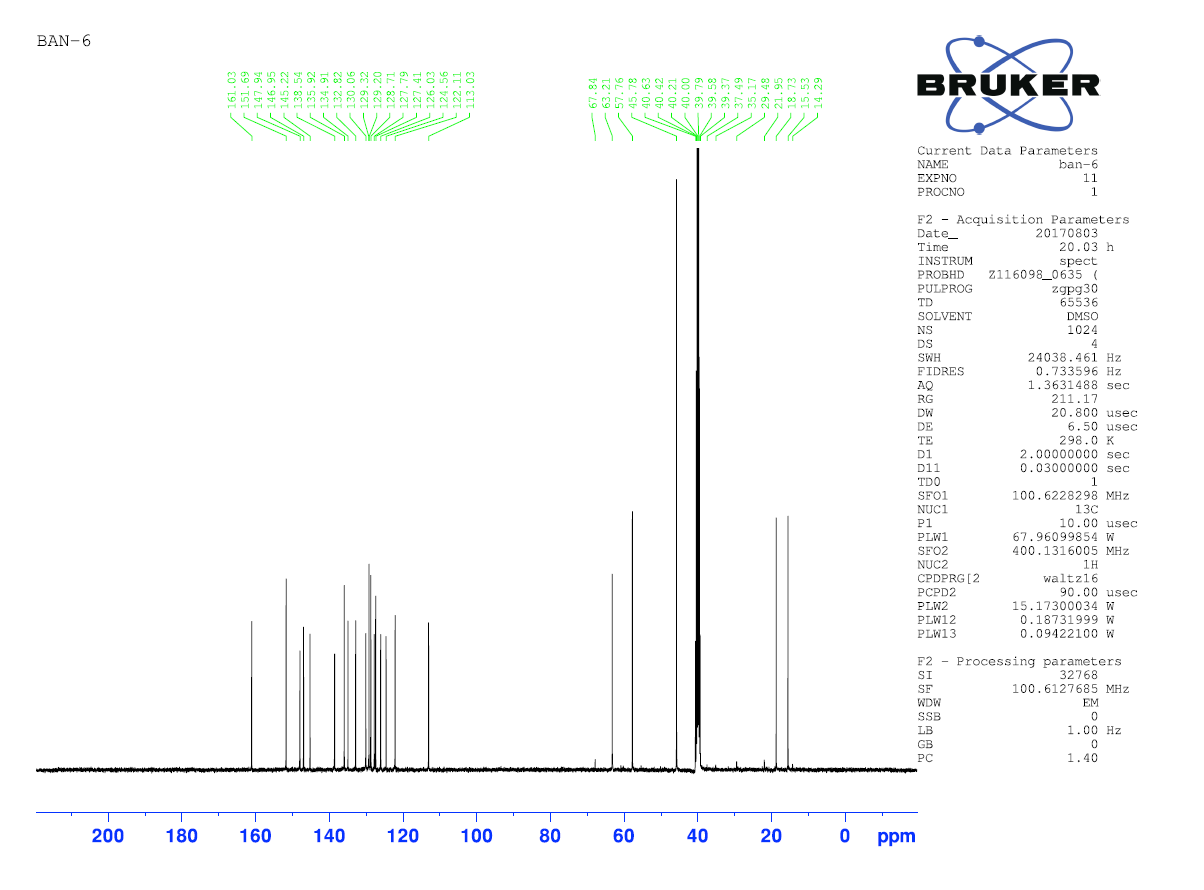

**^13^C-NMR of compound 3.8**

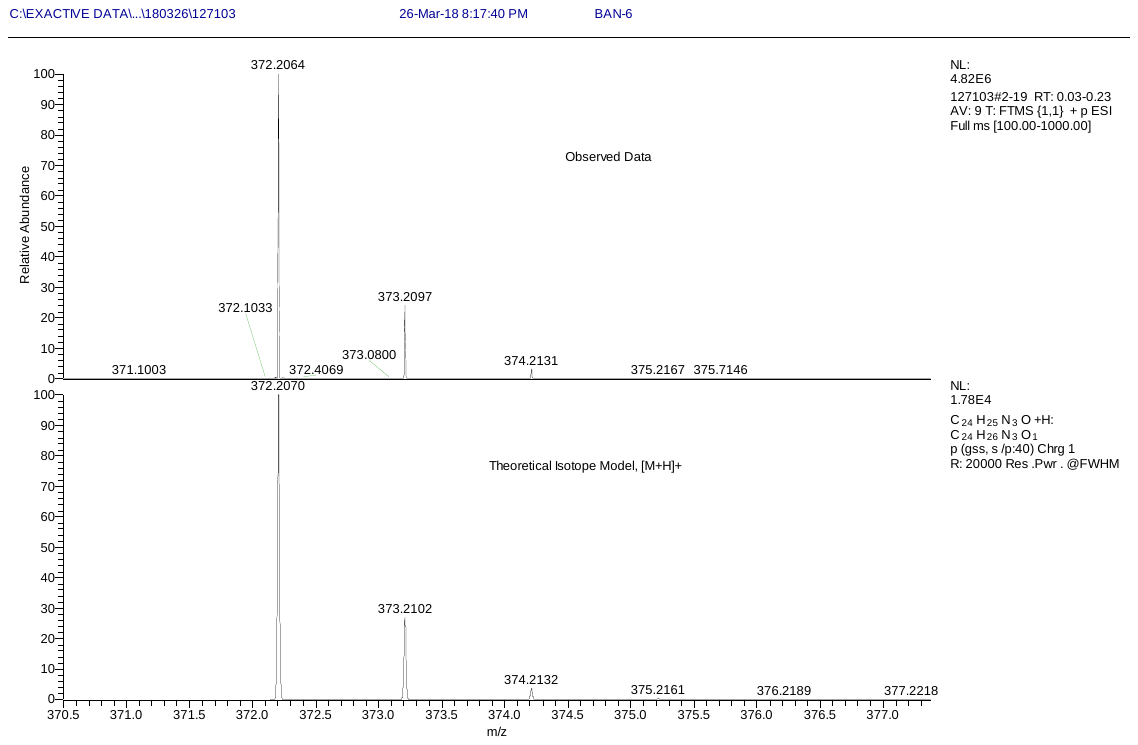

**HR-MS of compound 3.8**

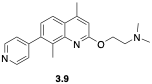

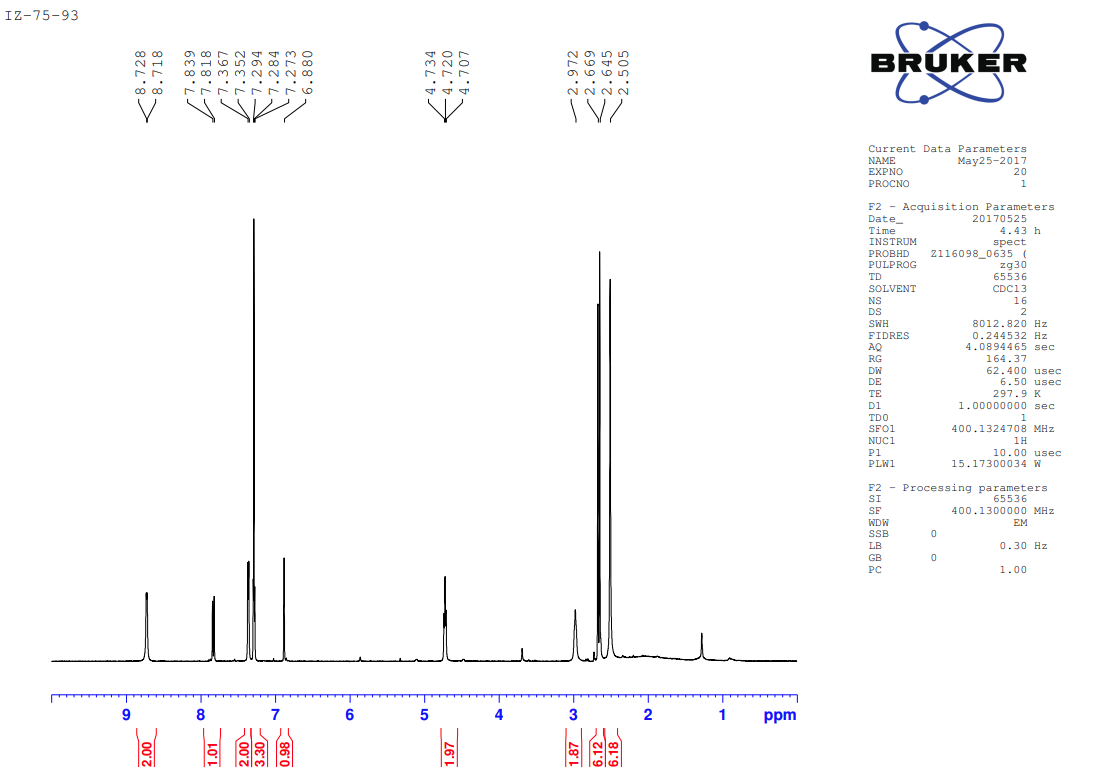

**^1^H-NMR of compound 3.9**

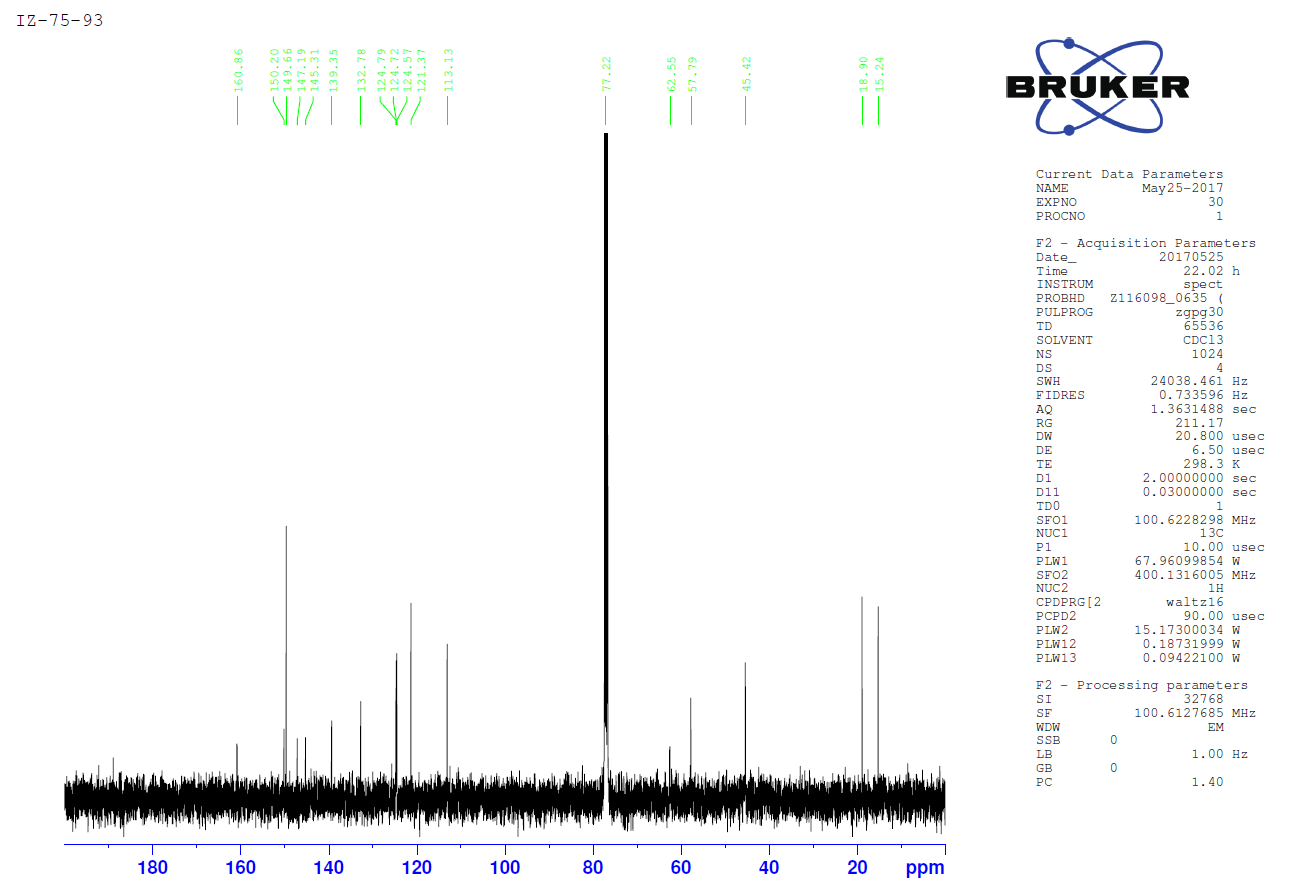

**^13^C-NMR of compound 3.9**

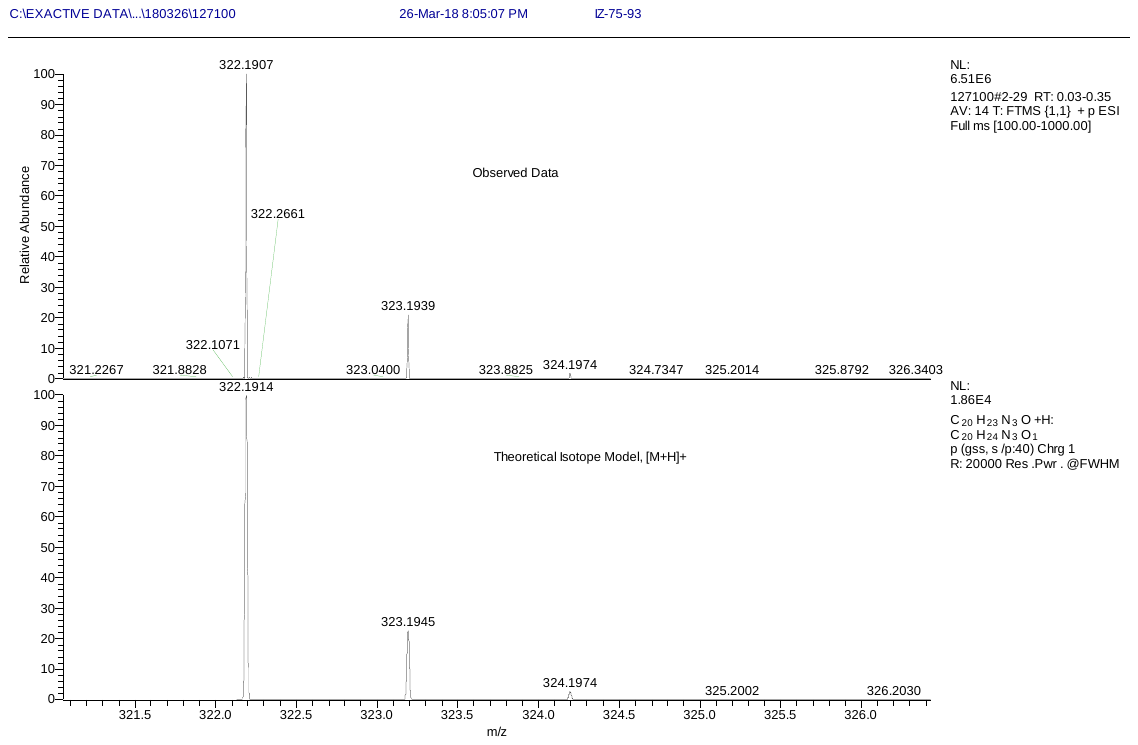

**HR-MS of compound 3.9**

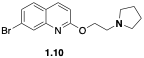

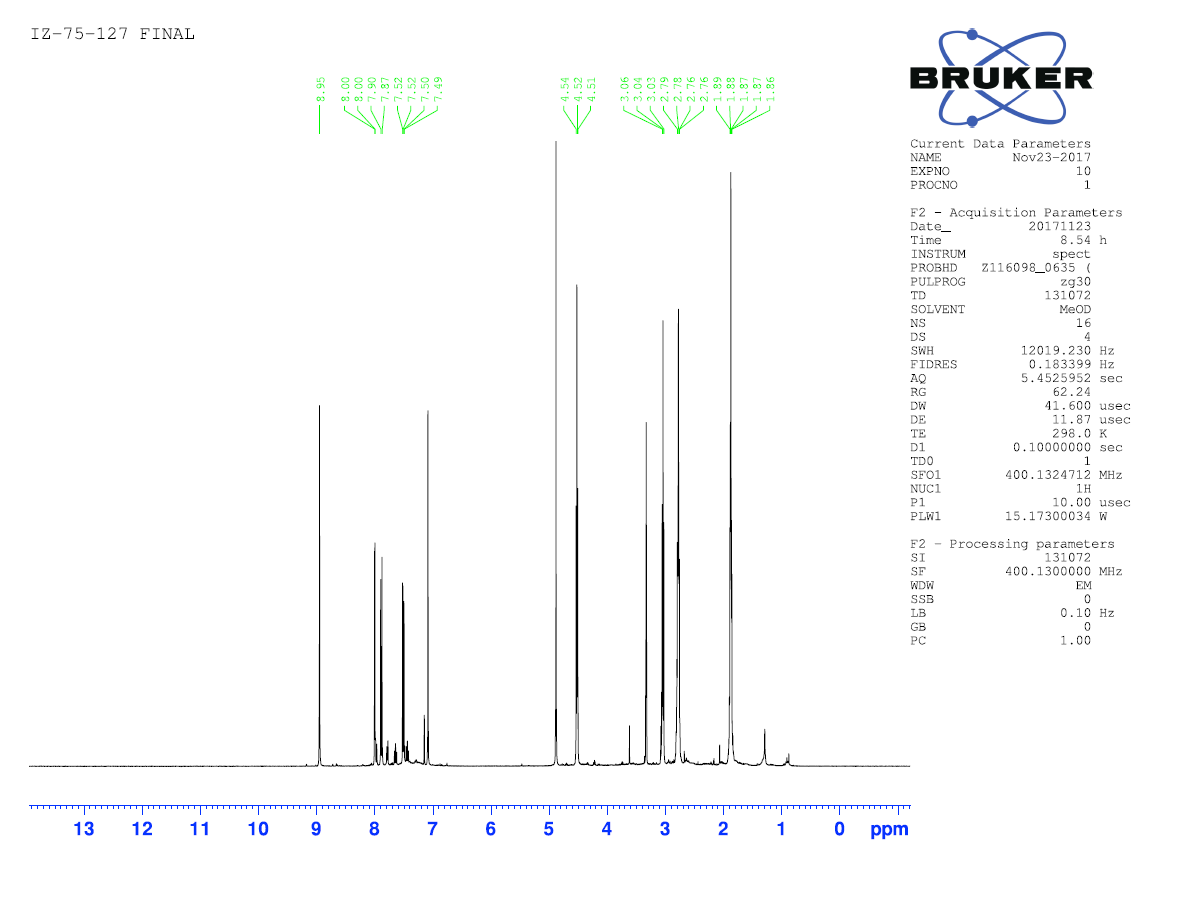

**^1^H-NMR of compound 1.10**

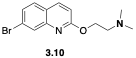

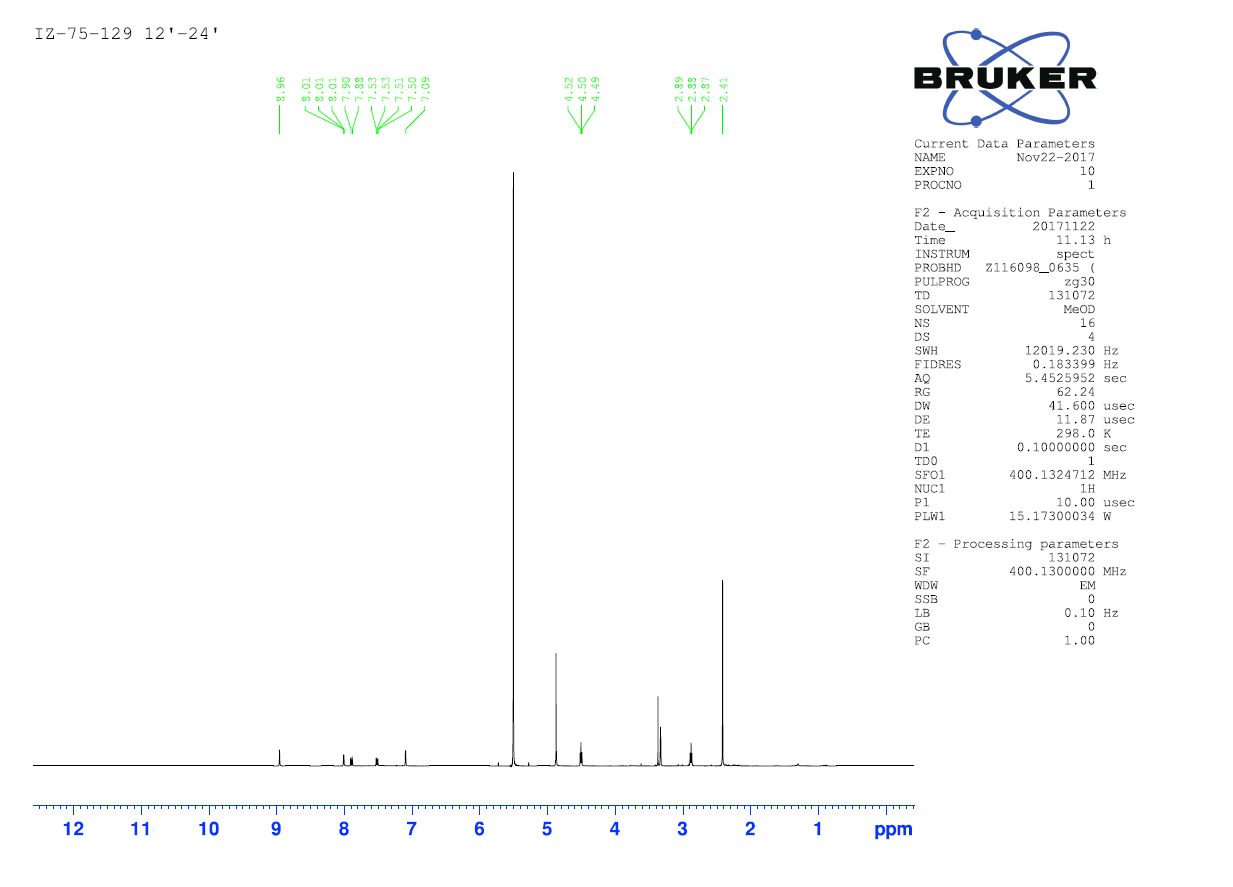

**^1^H-NMR of compound 3.10**

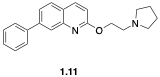

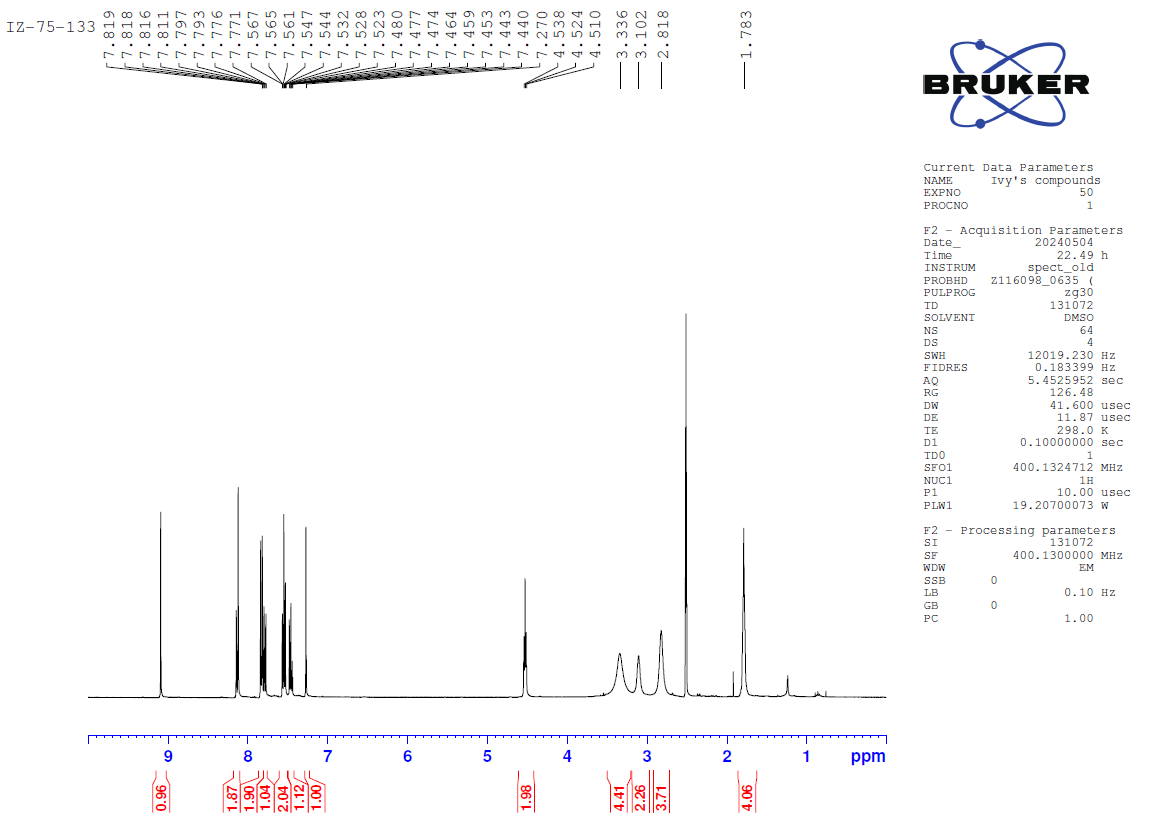

**^1^H-NMR of compound 1.11**

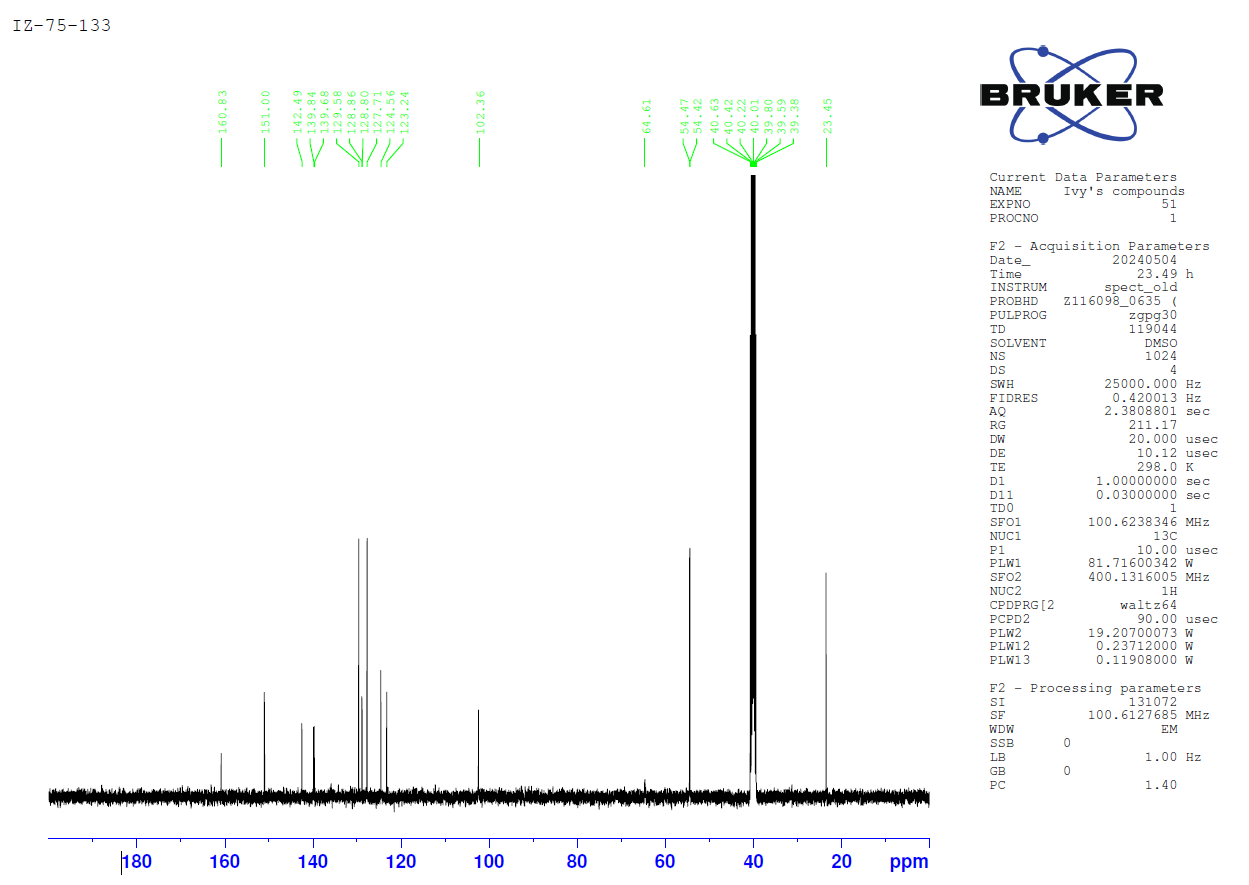

**^13^C-NMR of compound 1.11**

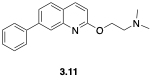

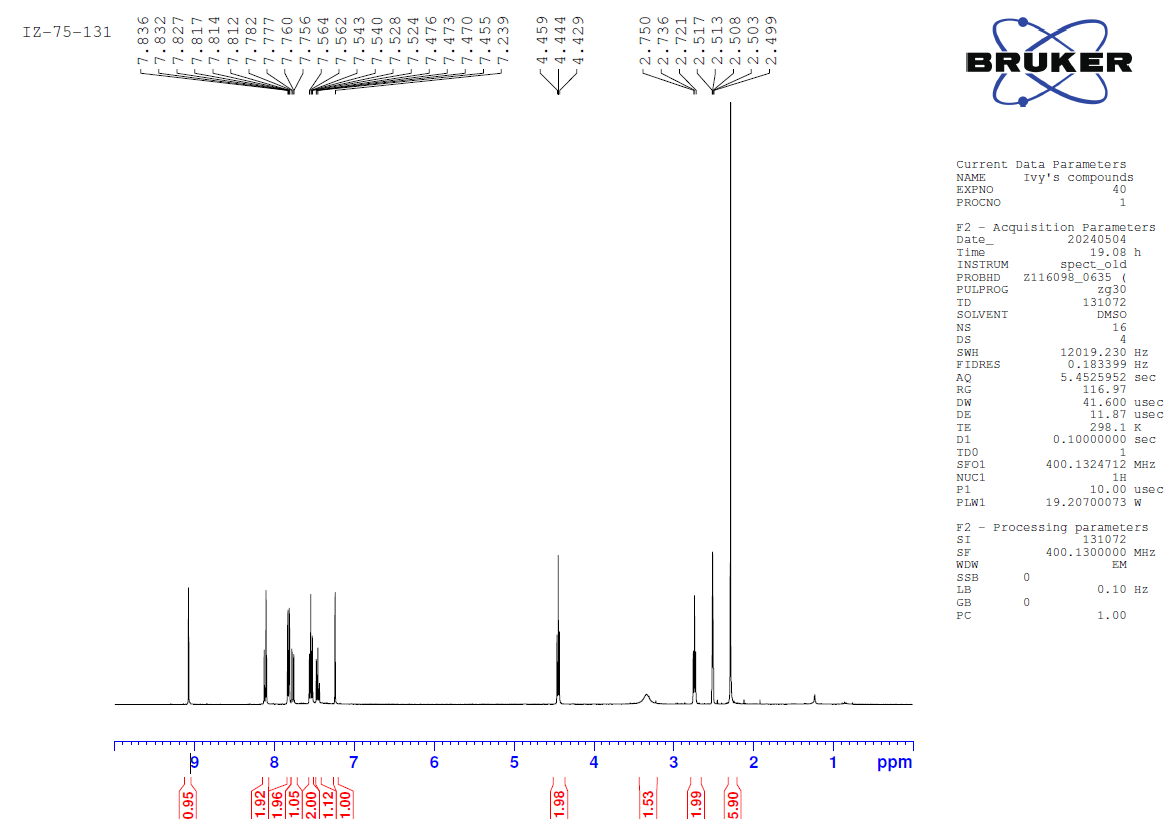

**^1^H-NMR of compound 3.11**

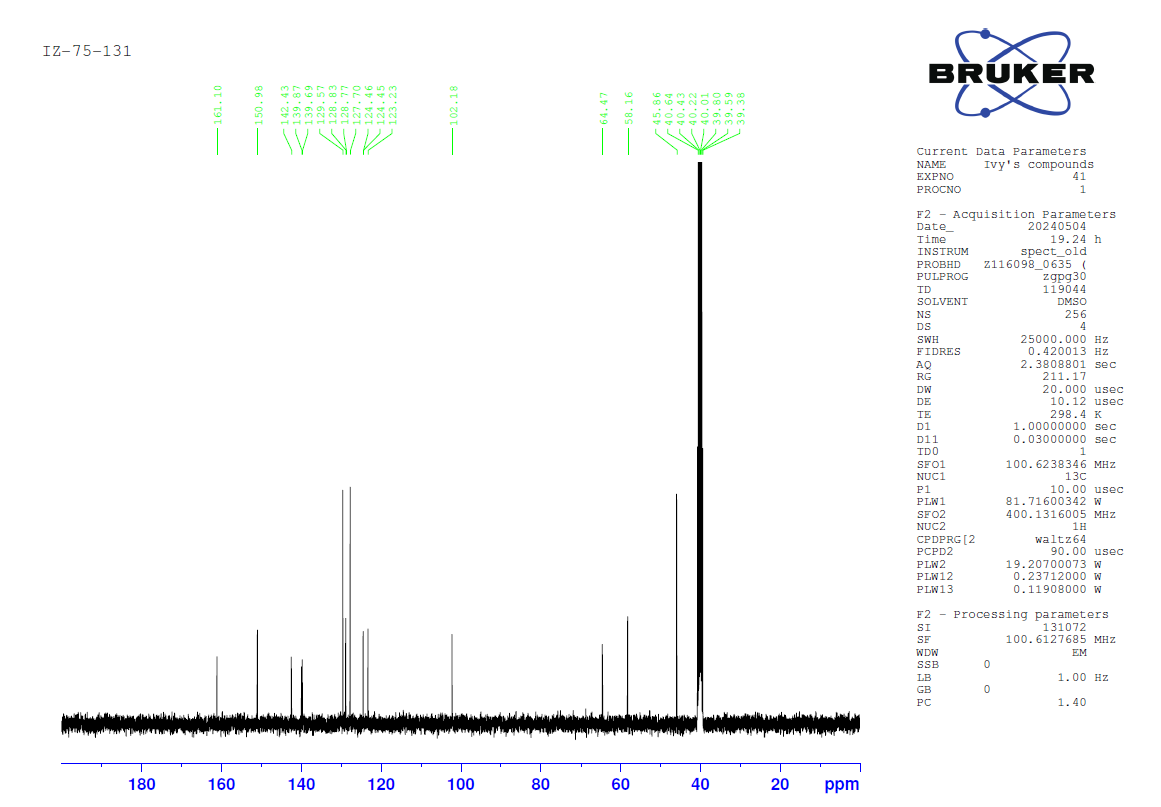

**^13^C-NMR of compound 3.11**

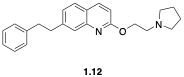

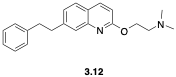

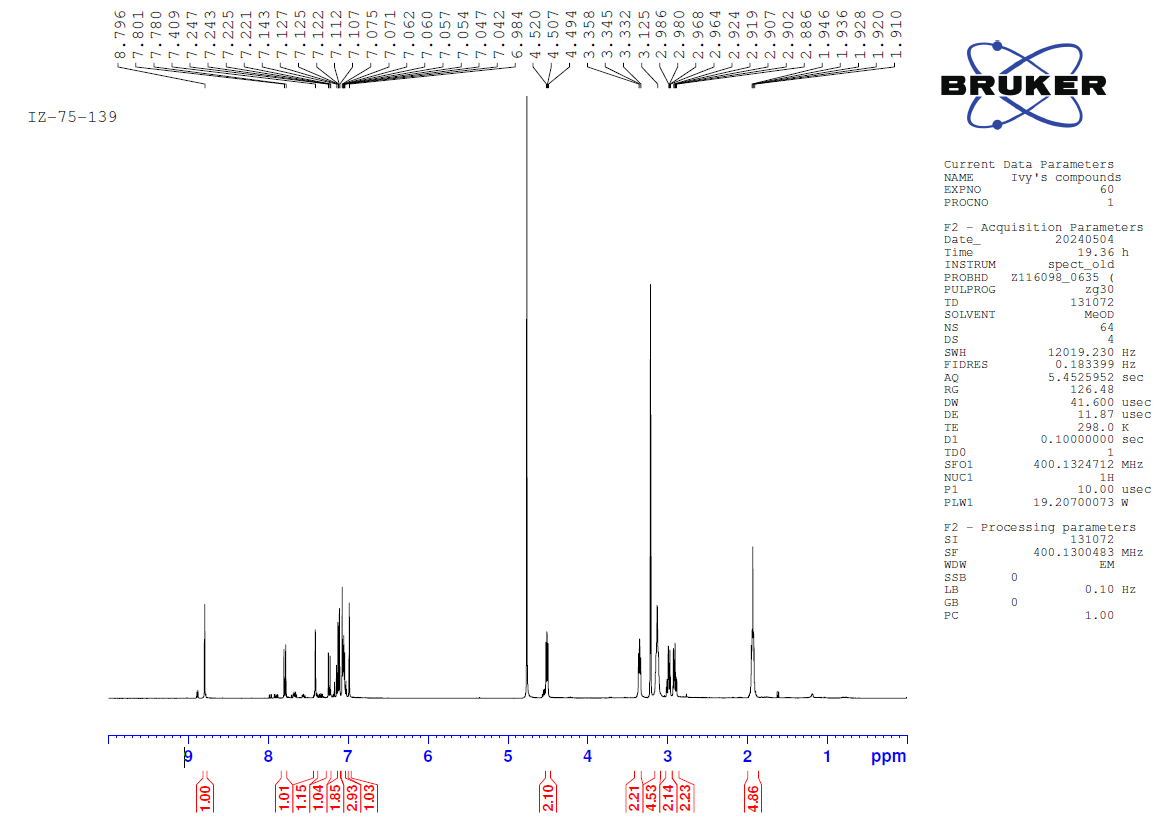

**^1^H-NMR of compound 3.12**

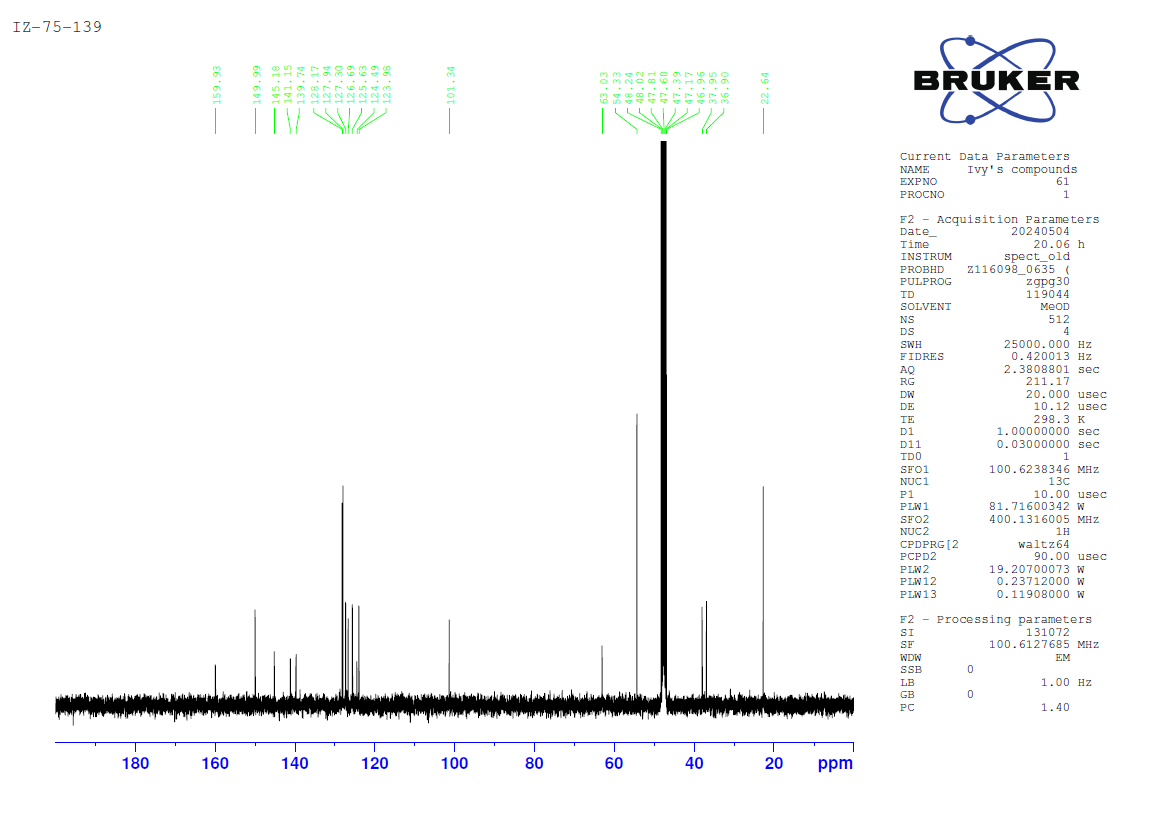

**^13^C-NMR of compound 3.12**
